# Epigenetic changes with age primes mammary luminal epithelia for cancer initiation

**DOI:** 10.1101/2021.02.12.430777

**Authors:** Rosalyn W. Sayaman, Masaru Miyano, Parijat Senapati, Sundus Shalabi, Arrianna Zirbes, Michael E. Todhunter, Victoria Seewaldt, Susan L. Neuhausen, Martha R. Stampfer, Dustin E. Schones, Mark A. LaBarge

## Abstract

Aging causes molecular changes that manifest as stereotypical phenotypes yet aging-associated diseases progress only in certain individuals. At lineage-specific resolution, we show how stereotyped and variant responses are integrated in mammary epithelia. Age-dependent directional changes in gene expression and DNA methylation (DNAm) occurred almost exclusively in luminal cells and implicated genome organizers *SATB1* and *CTCF*. DNAm changes were robust indicators of aging luminal cells, and were either directly (anti-)correlated with expression changes or served as priming events for subsequent dysregulation, such as demethylation of *ESR1*-binding regions in DNAm-regulatory *CXXC5* in older luminal cells and luminal-subtype cancers. Variance-driven changes in the transcriptome of both luminal and myoepithelial lineages further contributed to age-dependent loss of lineage fidelity. The pathways affected by transcriptomic and DNAm changes during aging are commonly linked with breast cancer, and together with the differential variability found across individuals, influence aging-associated cancer susceptibility in a subtype-specific manner.

## Introduction

The stereotyped aging phenotypes exhibited by organisms, organs, and tissues represent the integration of accumulated, stochastically incurred damages to individual cells that result commonly understood hallmarks of aging [Todhunter, Sayaman et al., 2018, López-Otín, 2013]. Age-associated directional changes in transcriptomes and methylomes of whole tissues are well documented [Volkova et al., 2005, de Magalhães et al., 2009, Glass et al., 2013, Peters et al., 2015, Christensen et al., 2009, Hernandez et al., 2011, Bell et al., 2012]. Age-dependent changes in DNA methylation (DNAm) has led to the development of epigenetic clocks [Horvath, 2013, Hannum et al., 2013, Weidner et al., 2014]. These directional molecular changes explain, at least in part, the noticeable phenotypic changes that accompany aging. Yet, in certain individuals, physiologic aging does not equate to chronologic aging. This discordance of physiologic and chronologic age reflects the variability that exists across individuals that may itself be an important molecular phenotype of aging [de Jong, et al., 2019]. As outliers, these individuals provide avenues for understanding biological processes that could explain why certain individuals appear physiologically younger or older than their chronological age. Indeed, variance between individuals arises in the contexts of tumors, diet, and aging [Xie et al., 2011, Slieker et al., 2016, Bashkeel, et al., 2019]. Increased susceptibility to a plethora of diseases, including cancers, is a prominent consequence of aging but the manifestation and onset of diseases vary between same-aged individuals. As shown in monozygotic twin studies, epigenetic changes with age modulate gene expression, differentially shaping phenotypes and significantly affecting their likelihood to progress towards a diseased state [Fraga et al., 2005]. Therefore, it is important to dissect how aging leads to both directional changes – where aging causes phenotypic shifts in one direction in a stereotyped manner; and variant changes – where aging causes increased variance that gives rise to diverging phenotypes. We propose that integration of directional and variant changes in aging may regulate the differential susceptibility between individuals.

The breast is an excellent model system for examining aging at the molecular and cellular levels because normal tissue across the adult lifespan is available from common cosmetic and prophylactic surgeries. Human mammary epithelial cells (HMEC) are readily isolated as primary cells enabling detailed and reproducible molecular studies that span the lifespan, and that model malignant progression [Stampfer et al., 2013]. Well-established lineage-specific markers and cell-sorting protocols facilitate experimentation at lineage-specific resolution. Moreover, breast tissue provides an ideal model for studying aging-associated cancer susceptibility as 82% of new breast cancers are diagnosed in women ≥50y [DeSantis et al., 2019]. Likewise, the breast provides an opportunity to study how subtype etiology is linked to aging as age-associated incidence rates of breast cancers vary in a subtype-specific manner – basal subtype estrogen receptor negative (ER-) breast cancer incidence rates stabilize whereas luminal A and B subtype ER+ breast cancer incidence rates continue to rise after age 50y [Tarone and Chu, 2002, Howlander, et al., 2014]. Whole breast tissue analyses have previously identified significant directional changes in gene expression and DNAm with age, which included changes that are associated with breast cancer biological processes [Yau et al., 2007, Lee and Lee, 2017, Song et al., 2017; Johnson et al., 2017]. However, aging is also associated with significant shifts in proportions of breast cell lineages, including epithelial and stromal populations [Garbe et al., 2012, Benz et al., 2014], so it is unclear how tissue-level molecular changes in normal aging reflected changes in cell-intrinsic and microenvironment states. Lineage-specific analyses are necessary to unravel such mechanisms.

The mammary epithelium, the origin of breast carcinomas, is a bilayer of two major cell lineages. Myoepithelial cells are basally-located, contractile, and thought to have tumor suppressive properties [Pandey et al., 2011]. Luminal epithelial cells are apically-located, secretory, and include subpopulations of hormone receptor positive cells [Booth and Smith, 2006]. Luminal cells and the mechanisms that maintain their lineage-specificity merit deep consideration in the context of breast cancer susceptibility because the cancer cells-of-origin of intrinsic breast cancer subtypes are thought to arise within the differentiation hierarchy of mammary stem cells to luminal progenitors to mature luminal cells [Prat and Perou, 2011, Lim et al., 2009; Molyneux et al., 2010, Pommier et al., 2020]. Luminal A and B intrinsic subtype expression and genomic profiles are closest to mature luminal cells [Tharmapalan et al., 2019, Prat and Perou, 2011]. Multipotent progenitors with basal differentiation bias accumulate with age. Luminal cells in older women exhibit expression of cytokeratins and other markers typically associated with myoepithelial cells, e.g. YAP1, keratin (K)14 and *TP63*; decreased expression of luminal markers, e.g. K19 and *ELF5*; and share protein expression signatures with immortalized HMECs [Garbe et al., 2012, Pelissier Vatter et al., 2018]. We describe this aging phenomenon as a loss of lineage fidelity, where the faithfulness of expression of certain lineage-specific markers with age diminishes without loss of lineage-specific expression of other canonical markers and gross phenotypic and histological differences between luminal and myoepithelial cells are maintained (Miyano, Sayaman et al., 2017). We hypothesize that the aging mechanisms that impinge upon maintenance of lineage fidelity are drivers of susceptibility to cancer initiation in breast tissue.

Here we show how age-dependent transcriptional and epigenetic stereotyped and variant responses integrate in breast epithelia and demonstrate how these changes may lead to increased susceptibility to cancer initiation. We examined genome-wide transcription in primary luminal and myoepithelial cells from 43 individuals across a wide spectrum of ages and cancer risk profiles, and matched DNA methylomes in a subset of these individuals. This approach allowed us to distinguish between directional and variant effects of aging. We showed that loss of lineage fidelity in gene expression was a genome-wide phenomenon recapitulated in DNAm. Age-dependent differential expression and DNAm explained, in part, the observed loss of lineage fidelity, while our model of the overall increase in variances with age also accounted for a considerable fraction of this loss. Strikingly, directional changes in expression and DNAm with age occurred almost exclusively in luminal cells, whereas changes in variance were found in both epithelial lineages. Genome-wide changes in luminal cells involved dysregulation of chromatin and genome organizers with age, including decrease of *SATB1* expression and changes in methylation at *CTCF* binding sites. Loss of lineage fidelity led to enrichment of genes and biological processes commonly dysregulated in cancers, as well as alteration of the luminal-myoepithelial interactome that was significantly modulated by apical cell-cell junction proteins, such as *GJB6*. DNAm changes were robust indicators of aging in luminal cells, and were either associated with canonical regulation of gene expression or served to prime aged luminal cells for subsequent dysregulation through secondary events such as binding transcriptional activators and repressors, DNAm regulatory proteins and chromatin remodeling complexes. An exemplar of this priming phenomenon is the demethylation of ESR1-binding regions in a key DNAm-regulatory protein CXXC5 in luminal cells of older women, which was also seen in age-related luminal-subtype cancers concordant with *CXXC5* and *ESR1* up-regulation. Many DNAm changes that were ascribed to early stage luminal-subtype breast tumors were already present in aged LEPs and represented the majority of loss of lineage fidelity events observed in the older cells. We show that lineage-specific interrogation revealed how lineage fidelity is lost through integration of genome-wide age-dependent stereotyped and variant changes in transcriptomes and methylomes. Primed DNAm states and biological variability across individuals underlie differential cancer susceptibility and could explain the differential prevalence of luminal breast cancer subtypes with age.

## Results

### Genome-wide loss of lineage-specific expression with age

To identify age-dependent changes in regulation of gene expression within the two main epithelial lineages of the breast (Figure 1A), RNA sequencing was performed on FACS-enriched luminal epithelial cells (LEPs) and myoepithelial cells (MEPs) isolated from 4^th^ passage (p4) finite-lifespan HMEC from reduction mammoplasties from two age cohorts: younger <30y women considered to be premenopausal (age range 16-29y, n=32 LEP and MEP samples, m=11 subjects) and older >55y women considered to be postmenopausal (age range 56-72y, n=22, m=8) (**Table S1**). We restricted analysis to the 17,165 genes with comparable dynamic ranges of gene expression levels and consistent lineage-specific expression between primary organoid and p4 LEPs and MEPs (linear regression *R^2^* ≥ 0.88 to 0.91, *p* < 2.2×10^-16^) (See methods) (**Figure S1A-B**).

**Figure 1.**
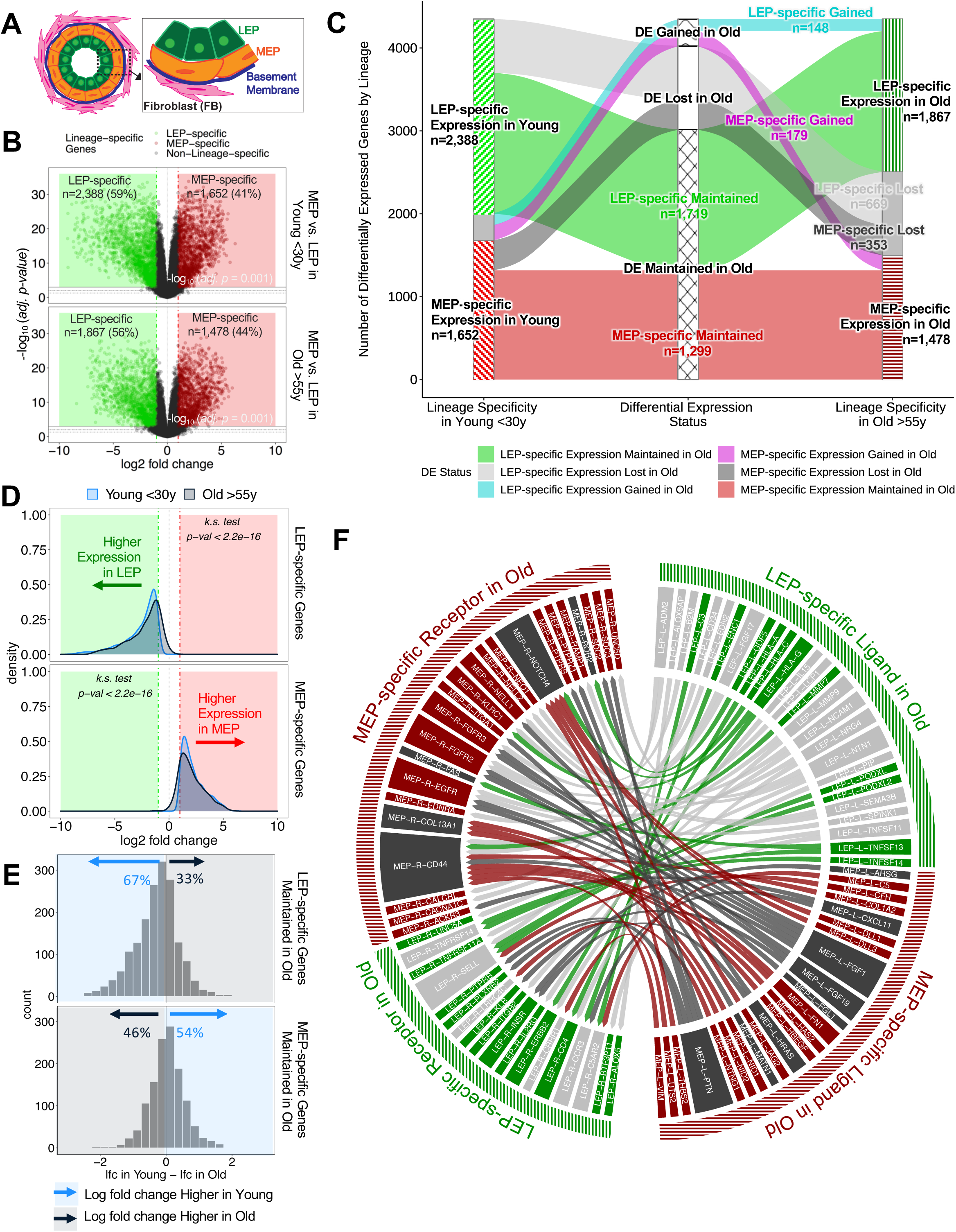
Genome-wide loss of lineage-specific expression with age. (**A**) A schematic depicting the two major epithelial lineages comprising the bilayered mammary epithelium. An apical layer of secretory luminal epithelial cells (LEPs) are surrounded by a basal layer of contractile myoepithelial cells (MEPs). The basement membrane surrounds the epithelial layer and separates it from the stromal compartment represented here by fibroblasts (FB). (**B**) Differential expression (DE) analysis identifies LEP-specific and MEP-specific genes (*limma* linear model *adj. p* < 0.001, fold change ≥ 2) in younger <30y (top panel) and older >55y (bottom panel) epithelial cells shown in a volcano plot of log_2_ fold changes (lfc) and -log_10_ BH adj. *p*-values. Number and percent of lineage-specific genes in each age cohort indicated. (**C**) Strata plot shows changes in lineage-specific DE with age, showing the number of LEP- and MEP-specific genes gained (cyan and magenta), lost (light and dark gray), and maintained (green and red) in older women (*adj. p* < 0.001, fold change ≥ 2). (**D**) Distribution of lfc in expression between LEPs and MEPs in younger (light blue) and older subjects (blue gray) for either DE LEP-specific (top panel) or MEP-specific (bottom panel) genes (*adj. p* < 0.001, fold change ≥ 2). LEP-specific genes are shown with (-) lfc values and MEP-specific-genes with (+) lfc values relative to each other; Kolmogorov-Smirnov (KS) test *p*-values for equality of distributions of lfc between younger and older women are annotated. (**E**) Histogram of pair-wise differences in lfc in expression between LEPs and MEPs in younger women vs. older women for genes that maintained DE of LEP-specific (top panel) and MEP-specific (bottom panel) genes with age. The percent of genes with higher lfc in younger women (light blue) or higher lfc in older women (blue gray) are indicated. (**F**) Interactome map of the ligand-receptor pairs (LRPs) [Ramilowski et al., 2015] with loss of lineage-specific DE of ligands and/or their cognate receptors (*adj. p* < 0.001, fold change ≥ 2) in older LEPs (light gray) or MEPs (dark gray). LRPs are connected by chord diagrams from the cell type expressing the ligand (cell type-L-gene symbol) to the cell type expressing the cognate receptor (cell type-R-gene symbol). See also **Figure S1**.

In lineage-specific analysis, the number of differentially expressed (DE) genes between LEPs and MEPs decreased with age (*limma* Benjamini-Hochberg (BH) *adjusted p-value, adj. p* < 0.05, < 0.01, < 0.001) (**Figure S1C**). Restricting analysis to genes with strong lineage-specific bias (*limma* DE *adj. p* < 0.001, fold change ≥ 2), we found 4,040 genes (23% of all genes analyzed) with highly significant lineage-specific DE in young women – of which 59% were LEP-specific and 41% were MEP-specific (Figure 1B top panel). In contrast, 3,345 genes had highly lineage-specific DE in older women – of which 56% were LEP-specific and 44% were MEP-specific (Figure 1B bottom panel). Shifts in lineage-specific expression with age were illustrated in the strata-plot in Figure 1C. Loss of lineage-specific expression with age was detected in 1,022 genes – a majority of which (65.5%) were LEP-specific. A gain in lineage-specific expression with age was also detected in 327 genes – of which 45% were LEP-specific and 55% were MEP-specific. These changes in lineage-specific expression occur genome-wide (**Figure S1D**).

We defined loss of lineage fidelity as a loss of the faithful expression of lineage-specific markers with age. Statistically, we described this loss as a phenomenon whereby the magnitude of gene expression differences that distinguish LEPs from MEPs decreases with age. This is seen as shifts in distributions of log_2_ fold changes (lfc) between lineages to smaller values in the older cohort (Kolmogorov-Smirnov two-sample test, KS *p* < 2.2×10^-16^) (Figure 1D). We found that 76% of LEP-specific genes (one-sided t-test *p* < 2.2×10^-16^) and 63% of MEP-specific genes (one-sided t-test *p* < 2.2×10^-16^) had higher lfc between lineages in younger cells compared to older cells (**Figure S1E**). These percentages indicated this phenomenon was not restricted to genes that had lost lineage-specific expression. Indeed, within the subset of genes where lineage-specific DE was maintained with age by significance threshold, the majority – 67% of LEP-specific genes (one-sided t-test *p* < 2.2×10^-16^) and 54% of MEP-specific genes (one-sided t-test *p* = 0.005), still showed larger lfc differences between LEPs and MEPs in younger women (Figure 1E). These data expand on our earlier findings that identified loss of lineage fidelity in a fairly small set of lineage-specific probes [Miyano, Sayaman et al., 2017], and underscore the genome-wide nature of this phenomenon whereby gene expression differences that distinguish lineages shift towards smaller lfc in women as they age.

### Loss of lineage fidelity with age leads to disrupted lineage-specific signaling

Because loss of lineage-specific expression could upset the relative balance of ligands and receptors synthesized by each lineage, we explored how loss of lineage fidelity could lead to disrupted or dysregulated cell-cell communication between neighboring cell types. We defined the breast interactome as a set of possible ligand-receptor interactions between cell populations in normal breast tissue based on the DE of cell-specific ligands and their cognate receptors [Sayaman, 2016]. Using published ligand-receptor pairs (LRPs) [Ramilowski et al., 2015], we first identified 224 candidate lineage-specific LRPs in young LEPs and MEPs based on the DE of 62 LEP-specific and 66 MEP-specific ligands, and 45 LEP-specific and 47 MEP-specific cognate receptors (**Figure S1F**). Protein-protein interaction (PPI) functional enrichment of lineage-specific LRPs identified top KEGG pathways (*stringdb* functional enrichment FDR *p* < 0.001) (**Figure S1G**), with ligands and receptors related to cytokine-cytokine receptor interaction and PI3K-Akt signaling commonly enriched in both LEPs and MEPs. LEP-specific ligands and receptors were enriched for cell adhesion molecules (CAMs) involved in cell-cell and cell-ECM interactions, and axon guidance molecules (AGMs), which have been found to play roles in mammary gland proliferation, adhesion, and migration [Harburg and Hinck, 2011]. MEP-specific ligands and receptors were enriched for ECM-receptor interaction and focal adhesion LRPs. LEP- and MEP-specific ligands and MEP-specific receptors were enriched for MAPK signaling and Rap1 signaling, which is involved in cell adhesion, cell-cell junction formation, and cell polarity processes. Enrichment of cytokine, immune, and infection-related pathways including antigen processing and presentation (LEP-specific ligands), complement and coagulation cascade (MEP-specific ligands), and hematopoietic cell lineage (MEP-specific receptors) pathways further suggested lineage-specific interactions between epithelial and immune cells.

Loss of lineage fidelity with age led to disruption of 74 LRPs based on the loss of lineage-specific expression of ligands and/or their cognate receptors (Figure 1F). For each lineage, we considered KEGG pathways (FDR *p* < 0.01) that were likely to exhibit dysregulated signaling either through direct disruption of the LRPs due to loss of lineage-specific signaling of the ligand or receptor, or indirect loss of signaling homeostasis via dysregulation of the cognate pair (**Figure S1H**). Loss of lineage-specific expression of LEP LRPs with age was enriched for KEGG pathways involved in (1) cell-cell and cell-ECM interactions including CAMs, AGMs and adherens junctions, and (2) cytokine, immune and infection-related pathways. Loss of lineage-specific expression of MEP LRPs with age were associated with (1) pathways in cancer; (2) pathways involved with MAPK, EGFR, NOTCH and PI3K-AKT signaling; and (3) MEP contractility. These findings suggest that loss of lineage fidelity with age has the potential to affect a wide range of biological processes regulating lineage-specific function and signaling, including potential dysregulation of cancer-related processes and immune-specific signaling by the epithelia.

### Loss of lineage fidelity with age is recapitulated in genome-wide DNA methylation

To understand how loss of lineage fidelity is maintained and made metastable during aging, we examined epigenetic regulation of gene expression via DNAm in a subset of FACS-enriched LEP and MEP samples (<30y, n=10 LEP and MEP samples, m=4 individuals; >55y, n=10, m=5) encompassing 347,280 probes (**Table S2**). We found a substantial decrease in the number of differentially methylated (DM) CpG sites and differentially methylated regions (DMRs) between LEPs and MEPs with age across multiple significance thresholds (*limma* and *DMRcate adj. p* < 0.05, < 0.01, < 0.001), suggesting our interpretation was not unduly affected by establishing arbitrary cutoffs (**Figure S2A-B**). Restricting analysis to highly lineage-specific CpG sites, we found 36,488 sites (11% of all CpG sites analyzed) with lineage-specific DM in younger cells (*adj. p* < 0.001, fold change ≥ 2) (Figure 2A, **S2A**), including 4,570 lineage-specific DMRs (*DMRcate* FDR *p* < 0.001) (**Figure S2B**). In stark contrast, only 15,866 sites showed lineage-specific DM in older cells (*adj. p* < 0.001, fold change ≥ 2) (Figure 2A**, S2A**), including 2,386 lineage-specific DMRs (FDR *p* < 0.001) (**Figure S2B**). This represented a loss of lineage-specific DM in 21,521 CpGs (7,276 genes) – 59% of all DM CpG sites with age (Figure 2A), which represent a much larger fraction of lineage-specific changes than seen in transcription. In addition, 899 CpGs (605 genes) with *de novo* lineage-specific DM unique to older cells were detected (Figure 2A). Moreover, loss of lineage fidelity phenomenon was recapitulated in DNAm, with the distribution in lfc at lineage-specific DM sites (*adj. p* < 0.001, fold change ≥ 2) shifting towards much smaller lfc between LEPs and MEPs in the older cohort (KS test *p* < 2.2×10^-16^) (**Figure S2C**) with 95.5% of LEP-specific CpG sites (one-sided t-test *p* < 2.2×10^-16^) and 93.6% of MEP-specific CpG sites (one-sided t-test *p* < 2.2×10^-16^) having higher lineage-specific lfc in younger women (**Figure S2D**). Loss of lineage fidelity in DNAm was arguably more striking than the loss of lineage fidelity observed in gene expression.

**Figure 2.**
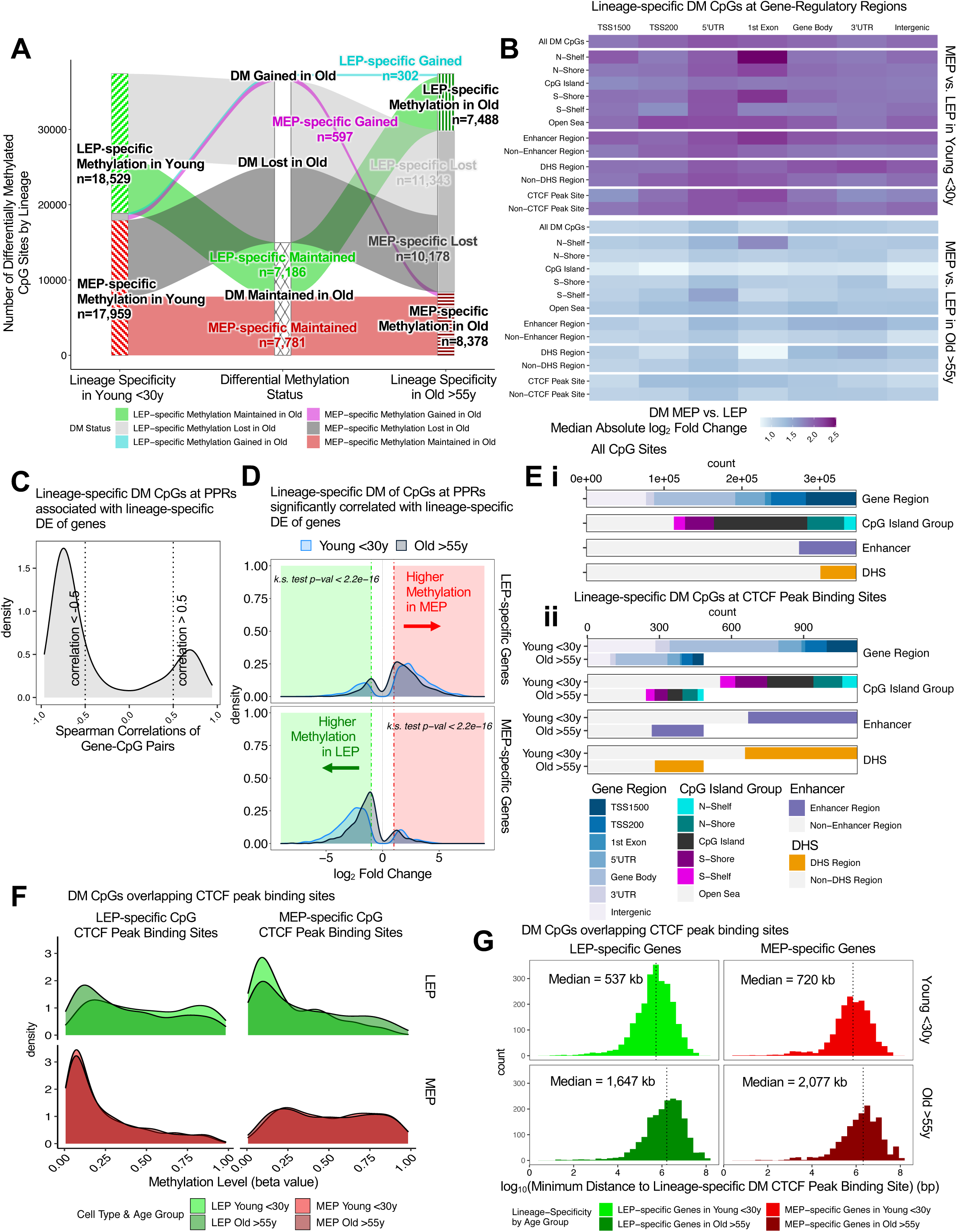
Loss of lineage fidelity with age is recapitulated in genome-wide DNA methylation. (**A**) Strata plot tracks changes in lineage-specific differential methylation (DM) with age, showing the number of LEP- and MEP-specific CpG sites which gain (cyan and magenta), lose (light and dark gray) or maintain (green and red) lineage-specific DM in older women (*limma adj. p* < 0.001, fold change ≥ 2). (**B**) Heatmap of the median value of DM absolute lfc between LEPs and MEPs in young (top panel) and older subjects (bottom panel) across gene regions: TSS1500, TSS200, 5’UTR, 1^st^ exon, gene body, and 3’UTR for all CpG sites and for specific annotated regulatory features: CpG island groups (CpG islands, N- and S-shores, N- and S-shelves and open seas), enhancer element regions, DNaseI hypersensitivity sites (DHS), and overlapping CTCF peak-binding sites. (**C**) Density distribution of Spearman rank-order correlation coefficients for gene-CpG pairs at lineage-specific DM CpGs located at promoter proximal regions (PPRs) that are associated with lineage-specific DE genes (DM and DE *adj. p* < 0.001, fold change ≥ 2). (**D**) Distribution of lfc in DNA methylation (DNAm) between LEPs and MEPs in young (light blue) and older subjects (blue gray) at DM CpG sites located at promoter proximal regions (PPRs) that are significantly correlated (Spearman cor *adj. p* < 0.05) with expression of either DE LEP-specific (top panel) or MEP-specific genes (bottom panel). CpG sites with higher DNAm in LEPs are shown with (-) lfc values and CpG sites with higher DNAm in MEPs with (+) lfc values relative to each other; KS test *p*-values for equality of distributions of lfc between younger and older women are annotated. (**E**) Total number of CpGs and proportions of annotated features: gene regions, CpG island groups, enhancer regions, DNaseI hypersensitivity sites across (**i**) All CpGs sites, and (**ii**) Lineage-specific DM CpGs (*adj. p* < 0.001, fold change ≥ 2) that overlap CTCF peak binding sites [ENCODE Project Consortium, 2011, 2012] in younger and older epithelial cells. (**F**) DNAm levels (beta values) of DM CpGs that overlap CTCF peak binding sites in LEPs (top) and MEPs (bottom) in younger epithelial cells, and their corresponding levels in older epithelial cells. (**G**) Histogram of genomic distances between lineage-specific DM CTCF binding sites and DE LEP-specific (left) or MEP-specific (right) genes in younger (top) vs. older (bottom) epithelial cells. Median genomic distances annotated. See also **Figure S2**.

Maintenance of lineage fidelity in younger epithelia was associated with lineage-specific DM at gene-regulatory regions that were lost with age. We first looked at DNAm across annotated genomic locations specifically at the promoter proximal regions (PPRs) that encompass: the regions 200-1200 bases (TSS1200) and 0-200 bases (TSS200) upstream of the transcription start site (TSS), 5’UTR between the TSS and ATG start site, and 1^st^ Exons, that are associated with canonical transcriptional regulation, as well as gene bodies, 3’UTRs and intergenic regions (**Figure S2E**). The largest differences (lfc) in DNAm between LEPs and MEPs in young women occurred at CpGs mapping to 1^st^ exons and 5’UTRs at shelves and shores of CpG islands (Figure 2B, **S2F**). We also saw strong differences in lineage-specific DM at enhancer regions at PPRs and at gene bodies, which suggested a role for exonic and intronic splicing enhancers, as well as at DNaseI hypersensitivity sites (DHS) associated with open chromatin regions. Further, we found lineage-specific DM at CTCF peak binding sites at 5’UTRs and 1^st^ exons, which suggested a role for chromatin organization in maintenance of lineage fidelity (Figure 2B). In contrast, there was an overall reduction in the magnitude of lineage-specific differences (lfc) in DNAm across all regions in older women (Figure 2B). N-shelfs located at 1^st^ exons remained as sites with largest differences in DNAm between LEPs and MEPs independent of age. DNAm of 1^st^ exons is more tightly linked to transcriptional silencing than upstream promoter regions [Brenet et al., 2011], suggesting 1^st^ exons may be a key regulatory site for lineage-specificity in breast epithelia.

### Loss of lineage-specific DNA methylation is highly correlated with loss lineage-specific expression

Enrichment of lineage-specific DM sites at PPRs and gene-regulatory regions suggested that DNAm regulates lineage-specific expression of LEP and MEP. We tested the correlation of DNAm and gene expression for 11,946 genes mapping to 184,678 CpG probes in the subset of individuals with matched data. Of the mapped genes, 26% (n=3,076) had expression levels that were significantly correlated (Spearman correlation, cor *adj. p* < 0.05) with methylation level of at least one associated CpG (**Figure S2Gi**). In contrast, of all the mapped CpG probes analyzed, only 5% (n=8,576) had methylation levels correlated (cor *adj. p* < 0.05) with gene expression (**Figure S2Gii**). That is, DNAm at 95% of all probed CpG sites had no direct correlation with gene expression (cor *adj. p* ≥ 0.05) (**Figure S2Hi**) regardless of genomic location or CpG island content (**Figure S2Hii**).

Lineage-specific genes (*adj. p* < 0.001, fold change ≥ 2) constituted 24% (n=2,839) of all mapped gene-CpG pairs, whereas they accounted for 56% (n=1,732) of significantly correlated gene-CpG pairs (cor *adj. p* < 0.05) (**Figure S2Gi**). Similarly, lineage-specific CpG sites (DM *adj. p* < 0.001, fold change ≥ 2) represented only 10% (n=18,780) of all mapped gene-CpG pairs, but accounted for 64% (n=5,517) of significantly correlated gene-CpG pairs (cor *adj. p* < 0.05) (**Figure S2Gii**) – a level of enrichment suggesting that metastability of lineage-specific expression was significantly regulated by DNAm. Considering gene-CpG pairs that had both lineage-specific expression and DNAm, we found 1,528 genes and 3,924 CpGs to be both lineage-specific and significantly correlated (DE and DM *adj. p* < 0.001, fold change ≥ 2, cor *adj. p* < 0.05) (**Figure S2Gi-ii, S2Ii**). Methylation of highly lineage-specific CpG sites located at gene bodies and 3’UTR were either correlated (cor > 0.5) or anti-correlated (cor < −0.5) with expression (**Figure S2Iii**). In contrast, methylation of CpG sites located at PPRs were mainly anti-correlated with expression of the associated gene (80% with cor *adj. p* < 0.05) (Figure 2C**, S2Iii**), which indicated a primary role for canonical regulation of gene expression through DNAm of PPRs. Indeed, in younger women, LEP-specific genes generally show higher methylation levels in MEPs, and MEP-specific genes show higher methylation in LEPs, at PPRs (Figure 2D**, S2J**). However, in older women, the lfc differences in methylation were much smaller in magnitude across all genomic locations between LEPs and MEPs at CpGs associated with lineage-specific genes (**Figure S2J**), and this decrease was most pronounced at PPRs (KS test *p* < 2.2×10^-16^) (Figure 2D).

These results establish that canonical transcriptional regulation via DNAm at PPRs plays a significant role in maintenance of epithelial lineage-specificity. Non-canonical regulation of expression also may occur at intragenic and intergenic regions, 3’UTRs and distal regulatory elements, as well as at gene bodies, which is thought to involve alternative splicing [Reddington et al., 2013, Maunakea et al., 2013, Lev Maor et al., 2015]. However, we lacked splicing-level resolution to further investigate this potential effect, and thus largely focused on the changes that occured at PPRs in the remainder of the paper.

### Loss of lineage-specific DNA methylation leads to disrupted CTCF binding

CCCTC-binding factor (CTCF), a TF that binds unmethylated DNA at CTCF binding sites with cell-type specific occupancies, is involved in chromatin organization, and plays a role in orchestrating genome-wide regulation of lineage-specificity [Loguercio et al., 2018, Kubo et al., 2020]. There were 20,450 CpG sites that overlapped HMEC CTCF uniform peak binding sites [ENCODE Project Consortium, 2011, 2012]. We found 1,125 lineage-specific DM sites in young women (*adj. p* < 0.001, fold change ≥ 2) that overlapped CTCF peak binding sites – of which 60% were lost in the older cohort. In contrast, only 484 lineage-specific DM sites were found in older women that overlapped CTCF peak binding sites – of which 6.4% were gained with age. Because CTCF is known to bind at enhancers, gene promoters, and inside gene bodies [Holwerda and de Laat, 2013], we compared the distributions of annotated genomic features across CTCF lineage-specific DM sites in younger cells relative to their distributions across all CpG sites. We found enhancer regions (41% CTCF DM CpG sites vs. 21% all CpG sites), gene bodies (40% vs. 30%), open seas (49% vs 32%) and DHS (42% vs. 13%) were relatively enriched, whereas PPRs (29% vs 45%) and CpG islands (17% vs 35%) were relatively depleted (Figure 2Ei-ii). Moreover, LEP-specific CTCF sites were largely unmethylated in MEPs, and MEP-specific CTCF sites were largely unmethylated in LEPs across genomic features, consistent with known CTCF TF binding of unmethylated regions (Figure 2F). Loss of lineage-specific methylation with age at CTCF sites was seen across genomic features (Figure 2Eii) and was associated with hypomethylation of LEP-specific CTCF CpG sites and hypermethylation of MEP-specific CTCF CpG sites in aged LEPs (Figure 2F).

We next examined the distance of lineage-specific DE genes (adj. *p* < 0.001, fold change ≥ 2) to their nearest lineage-specific DM CTCF peak binding sites within each age group. The median distance between lineage-specific DE genes and the nearest DM CTCF sites was three times farther in older cells than in younger cells (Figure 2G). Lineage-specific DE of the nearest gene or DM of the nearest CTCF peak binding sites were lost in older cells (**Figure S2K**).

These results suggest that changing DNAm patterns at CTCF peak binding sites in older epithelia, particularly in older LEPs, disrupts the ability of CTCF to form chromatin loops with other CTCF binding sites, enhancers, and their targets genes. Such alterations in 3D chromatin architecture may play a role in the loss of lineage fidelity, which reflects concerted genome-wide dysregulation.

### Models of loss of lineage fidelity in breast epithelia

To understand the changes within each cell population that contribute to the observed aging-associated loss of lineage fidelity, we explored two models that could explain the decrease in DE between LEPs and MEPs with age. The first model took into account age-dependent directional changes, either through stereotypic up-regulation or down-regulation that led to loss of lineage-specific expression – e.g., LEPs acquire MEP-like expression patterns and/or MEPs acquire LEP-like expression patterns in the older cohort (Figure 3Ai). The second model considered aging-associated increase in variances in the expression of lineage-specific genes in LEPs and/or MEPs from older women, leading to a loss of detection of DE between lineages (Figure 3Aii). We describe the contributions of each in the following sections.

### The luminal lineage is a hotspot for age-dependent directional changes

We examined age-dependent DE of genes and DM of CpG sites within each epithelial lineage. We found an extreme lineage bias in the numbers of DE genes, DM CpG sites, and DMRs between younger and older cells, with the majority of age-dependent changes occurring in LEPs. This phenomenon held regardless of the significance thresholds that were applied (*limma* and *DMRcate adj. p* < 0.05, < 0.01, < 0.001) (**Figure S3A-C**). In LEPs, 471 genes were DE, and 18,522 CpG sites and 3,141 regions were DM as a function of age (*limma* and *DMRcate adj. p* < 0.05) (Figures 3B, 3C). In contrast, in MEPs only 29 genes were DE, and only 20 CpG sites and 1 region were DM with age (*adj. p* < 0.05) (Figures 3B, 3C). This represented a ∼900-fold difference in age-dependent DM in LEPs compared to MEPs. Moreover, we identified age-dependent changes that showed lineage independence, with five genes commonly DE, and 14 CpG sites mapping to seven genes commonly DM with age in both LEPs and MEPs. This left only 24 genes and six CpG sites that changed with age exclusively in MEPs. Relative to MEPs, LEPs had significantly lower lineage-specific expression of DNA methyltransferase, *DNMT1* (*adj. p* = 1.3×10^-12^), which is involved primarily in maintenance of DNAm [Cedar and Bergman, 2011]; and significantly higher expression of ten-eleven-translocation enzyme, *TET2* (*adj. p* = 2.2×10^-9^), which catalyzes DNA demethylation through oxidation of 5-methylcytosine to 5-hydroxymethylcytosine [Pastor et al., 2013] (Figure 3D). These DNAm regulatory enzymes showed similar lineage-specific expression in organoids (*adj. p* = 3.8×10^-7^ and *adj. p* = 8.7×10^-12^ respectively) (**Figure S3D**), suggesting that intrinsic properties of the lineage play a role in its ability to maintain DNAm levels over a time course of physicochemical insults. That stereotypic directional changes associated with aging were almost exclusively found in LEPs suggests that this lineage could serve as a primary indicator of aging – a proverbial canary in the coalmine.

**Figure 3.**
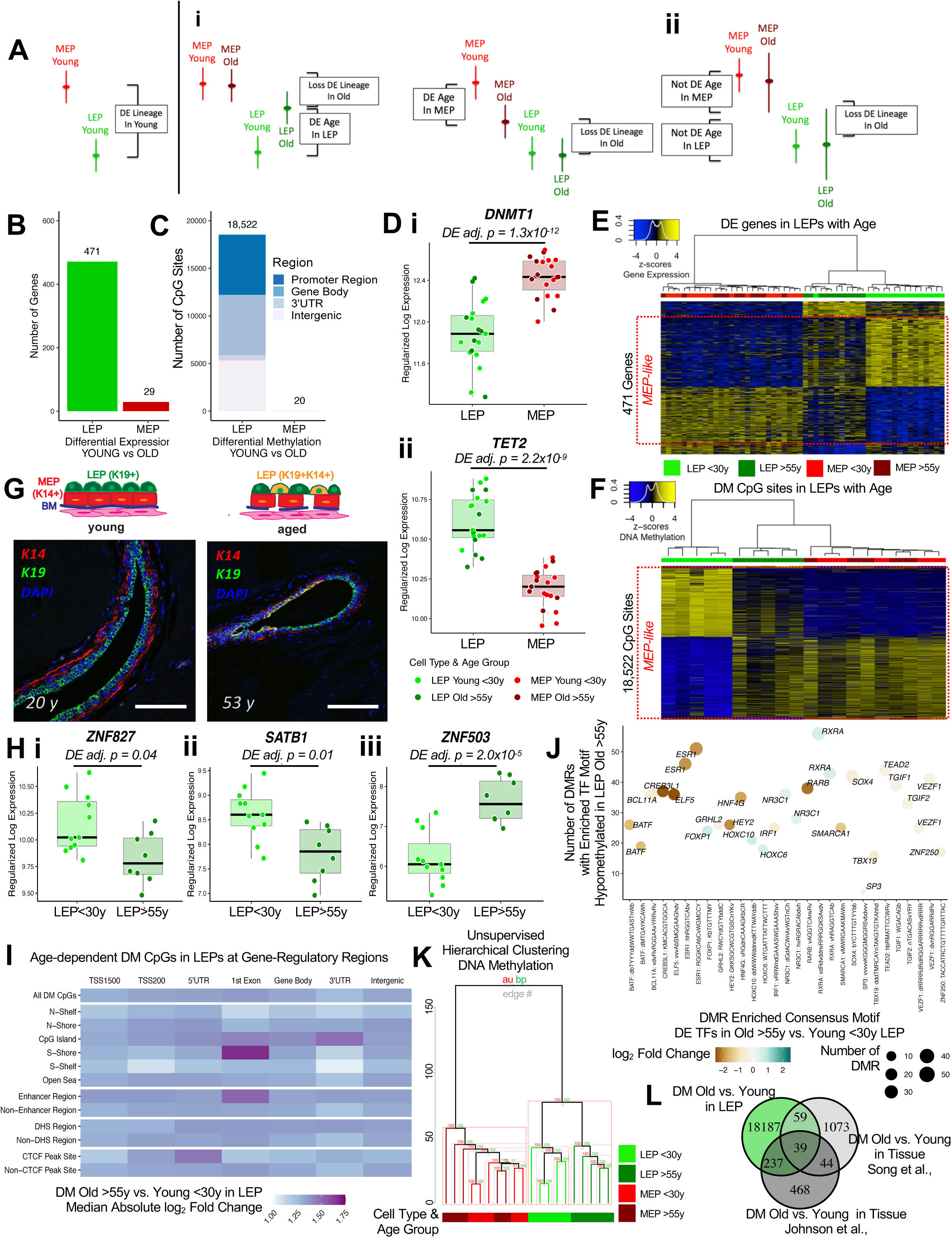
The luminal lineage is a hotspot for age-dependent directional changes. (**A**) Models of loss of lineage fidelity illustrate hypothesized mechanisms leading to loss of lineage fidelity (**i**) Age-dependent DE shifts in gene expression in LEPs and/or MEPs of older relative to young cells; or (**ii**) An increase in gene expression variance in older LEPs and/or MEPs that lead to loss of detection of lineage-specific DE between LEPs and MEPs with age. (**B-C**) Number of (**B**) age-dependent DE genes (*limma* DE *adj. p* < 0.05) and (**C**) age-dependent DM CpG sites (*limma* DM *adj. p* < 0.05) between younger and older LEPs or MEPs. (**D**) Boxplots of gene expression rlog values of (**i**) *DNMT1* and (**ii**) *TET2* in LEPs and MEPs. MEP vs. LEP DE adj. *p*-values (*limma*) annotated. (**E-F**) Hierarchical clustering of all samples based on (**E**) age-dependent DE genes and (**F**) age-dependent DM CpG sites in LEPs (DE and DM *adj. p* < 0.05). Scaled rlog values of DE genes and scaled beta values of DM CpG sites are shown as a heatmap. Clustering performed using Euclidean distances and Ward agglomerative method. (**G**) LEP-specific K19 and MEP-specific K14 immunofluorescence staining in breast tissue from a younger and an older reduction mammoplasty. (**H**) Boxplots of gene expression rlog values of (**i**) *ZNF827*, (**ii**) *SATB1* and (**iii**) *ZNF503* in LEPs of younger and older LEPs. Age-dependent DE adj. *p*-values (*limma*) are indicated. (**I**) Heatmap of the median value of DM absolute lfc between younger and older LEPs across gene regions: TSS1500, TSS200, 5’UTR, 1^st^ exon, gene body, and 3’UTR for all CpG sites and for specific annotated regulatory features: CpG island groups (CpG islands, N- and S-shores, N- and S-shelves and open seas), enhancer element regions, DHS, and overlapping CTCF peak-binding sites. (**J**) Number of DMRs (*DMRcate* FDR *p* < 0.05) that lost DNAm with age with enriched motifs (*ELMER* FDR *p* < 0.05) for DE TFs in LEPs (*adj. p* < 0.05). Point sizes reflect the number of DMRs with motif enrichment and are color coded based on DE lfc of the corresponding TFs. (**K**) Unsupervised hierarchical clustering of LEPs and MEPs from younger and older women based on DNAm beta values of all CpG sites. Clusters with approximately unbiased *p*-value, AU ≥ 95 (*pvclust*) are highlighted (dashed red, with the largest supported cluster in solid red). Clustering performed using Euclidean distances and Ward agglomerative method. (**L**) Venn diagram of CpG sites identified to be DM with age in LEPs in our study (*adj. p* < 0.05) compared to those identified in bulk tissue from two independent datasets [Song et al., 2017, Johnson et al., 2017]. See also **Figure S3**.

Age-dependent up-regulation and down-regulation of gene expression, as well as gain and loss of DNAm at CpG sites, occurred at comparable frequencies in LEPs (*adj. p* < 0.05) (Figure 3E-F). In LEPs, 82% of the genes that were DE and 97% of the DM CpG sites with age changed in a direction towards acquiring MEP-like patterns (Figure 3E-F). These directional changes in methylation patterns in older LEPs were more pronounced than expression shifts, and led to clustering of older LEPs more closely to MEPs than to younger LEPs (Figure 3F). In particular, gain of methylation at over half of CpG sites (53%) in older LEPs led to DNAm levels comparable to MEPs, whereas loss of methylation (47%) in older LEPs led to more partially-methylated DNAm levels intermediary to those observed in young LEPs and MEPs (Figure 3F). These molecular changes recapitulated previous phenotypic observations of luminal basalization where LEPs in breast tissues of older women acquired expression of MEP-specific marker K14, while also maintaining expression of LEP-specific marker K19 [Petersen et al., 2010; Garbe et al., 2012] (Figure 3G). Although changes in MEPs were far fewer, we note that shifts in expression and methylation in older MEPs led to “LEP-like” patterns (**Figure S3E-F**).

Expression of key LEP regulatory TFs [Lambert et al., 2018] shifted towards more MEP-like patterns in LEPs with age. Fifty TFs were DE between young and old LEPs (**Figure S3G**), the majority of which have known roles in breast cancer progression. Of these TFs, 88% changed expression in older LEPs towards the direction of MEP-like expression. These included highly expressed TFs in young LEPs that were down-regulated with age, such as: *ELF5*, *GRHL2* (**Figure S3Hi**), *HES4* (**Figure S3Hii**), *SGSM2* (**Figure S3Hiii**), *SOX4*, *TEAD2*, *ZNF641*, *ZNF827* (Figure 3Hi), and genome organizer *SATB1* (Figure 3Hii)*. ZNF827* mediates telomere homeostasis through recruitment of DNA repair proteins [Vilas et al., 2018]. Loss of *GRHL2* and *SGSM2* are associated with down-regulation of E-cadherin and epithelial-to-mesenchymal transition (EMT) in mammary epithelial cells [Xiang et al., 2012, Lin et al., 2019]. *HES4* is a canonical target gene of Notch1 which plays an important role in normal breast epithelial differentiation and cancer development [Kontomanolis et al., 2018]. And *SATB1* has genome organizing functions in stem cells and tumor progression [Kohwi-Shigematsu et al., 2013]. In addition, several TFs gained expression in older LEPs, including: *SP3* (**Figure S3Hiv**)*, RXRA, SOX15* and *ZNF503* (Figure 3Hiii). *ZNF503* inhibits *GATA3* expression, a key regulator of mammary LEP differentiation, and down regulation is associated with aggressive breast cancers [Shahi et al., 2017, Kouros-Mehr et al., 2006]. And *SP3* silencing inhibits Akt signaling and breast cancer cell migration and invasion [Mansour, 2020]. Aging caused significant changes in TFs in the LEP lineage that are known to have roles in breast cancer.

To better understand how age-dependent DNAm changes in LEPs related to gene expression, we examined the regulatory regions and features where the largest changes in DNAm occurred with age. Comparing younger to older LEPs, the largest median absolute-lfc across all DM sites occurred at: CpG islands in 5’UTRs, 1^st^ exons, gene bodies and 3’UTRs (Figure 3I**, S3I**), as well as S-shores and enhancer regions at 1^st^ exons, and at CTCF binding sites in 5’UTRs (Figure 3I) – further implicating chromatin remodeling in age-dependent changes in LEPs. The 1^st^ exon region continued to be a hotspot region exhibiting the largest changes in both lineage-specific and age-dependent DE (Figure 2B, 3I). Of 391 age-dependent DE genes in LEPs (*adj. p* < 0.05) that also had mapped methylation data, 206 (52%) had expression levels correlated with methylation of at least one CpG site (cor *adj. p* < 0.05), and 128 (33%) additionally had CpG sites that were significantly DM with age (*adj. p* < 0.05) (**Figure S3Ji**). The corresponding 277 CpG sites associated with significantly correlated and age-dependent gene-CpG pairs in LEPs (**Figure S3Jii-S3Ki**) occurred primarily at PPRs (55%) that were predominantly anti-correlated (92%) (cor < −0.5, *adj. p* < 0.05) with expression; and at gene bodies (41%) that showed both correlation (34%) and anti-correlation (65%) with expression (cor < −0.5 or cor > 0.5, *adj. p* < 0.05) (**Figure S3Kii**). These results suggest both canonical and non-canonical roles of DNAm in aging phenotypes through regulation of PPR and gene body methylation.

Next, we performed TF motif enrichment analysis at each of the 3,141 age-dependent DMRs (*ELMER* FDR *p* < 0.05) [Silva et al., 2019]. Half (50%) of DMRs between younger and older LEPs had motifs enriched within +/-250bp of the given CpG sites. We highlighted the most common consensus motifs (HOCOMOCO v11) [Kulakovskiy et al., 2017], and the top 60 TFs associated with DMRs that lost methylation (n ≥ 50 DMRs) in either younger (**Figure S3Li**) or older LEPs (**Figures S3Lii**). One highly common consensus motif enriched in DMRs (hvYCGCGAGAn) was associated with transcriptional regulator *ZBTB33* (*KAISO*), which is known to have bimodal DNA-binding specificity – capable of binding methylated CGCG motif as well as non-methylated consensus motifs, and to repress transcription through recruitment of the chromatin remodeling machinery [Buck-Koehntop, et al., 2012]. Moreover, two of the most common enriched motifs were associated with LEP-specific TF *ELF5* (E74 Like ETS Transcription Factor 5), which was highly expressed in LEPs of younger women (*adj. p < 0.05*) and were highly enriched in DMRs that were hypomethylated in the younger cohort. Canonical MEP-specific TF *TP63* which was expressed higher in older LEPs at nominal significance (*unadj. p* < 0.005, *adj. P* < 0.1) and was highly enriched in DMRs that were hypomethylated in the older cells (**Figures S3Li-ii**), supporting our earlier findings based on a more limited set of whole genome bisulfite sequencing (WGBS) data in LEPs [Senapati et al., 2020].

To further understand how changes in TF expression with age were linked to hypomethylation events in older LEPs, we examined the 24 TFs DE with age (*adj. p* < 0.05) that had enriched motifs within the 913 DMRs that lost methylation in older LEPs (Figure 3J). Of these, 18 TFs were down-regulated in older LEPs, suggesting that loss of TF expression could lead to alternate binding of other regulatory proteins at these hypomethylated sites. Another six TFs up-regulated in old LEPs were enriched in DMRs that lost methylation with age, including in 190 DMRs that overlapped known gene promoter regions (**Figure S3M**), suggesting that binding of upregulated transcriptional activators or repressors at these sites could directly lead to further downstream dysregulation of transcription with age. Of these up-regulated TFs, homeobox *HOXC6*, which is involved in mammary gland development, and *HOXC10* (**Figure S3Ni-ii**) are both regulated by estrogen and MLL histone methylases, and are overexpressed in breast cancers [Hussain et al., 2015, Ansari et al., 2012]. Thus, hypomethylation of DMRs with age could serve as priming events for subsequent binding of overexpressed TFs associated with aging and breast cancers.

The striking LEP lineage bias in age-dependent DE and DM suggested that epithelial lineages of the breast age via different mechanisms or at different rates. Indeed, predicted ages of isogenic MEP and LEP pairs, calculated using Horvath’s 353-CpG pan-tissue DNAm clock [Horvath, 2013], showed that MEPs exhibit physiologic ages that were accelerated on average 12+/-6y compared to their isogenic LEP counterparts (**Figure S3O**). Further, older subjects exhibited larger variances in the difference in the biological ages of LEP and MEP pairs. Thus, even though LEPs and MEPs originate from common progenitors and reside juxtaposed in tissue, their biological ages may be out of phase and biological clocks may be sensitive to the changing composition of the breast with age. Indeed, 77 of the 353 CpG markers showed lineage-specific DM in younger cells, 31 showed lineage-specific DM in the older cells (*adj. p* < 0.05), and 29 CpGs showed age-dependent DM only in LEPs (*adj. p* < 0.05).

To understand the contribution of age-dependent directional changes in driving loss of lineage fidelity (Figure 3Ai), we examined the overlap of age-dependent DE genes or DM sites with genes or CpG sites that lost lineage-specific DE or DM, respectively. Only 9% of the loss in lineage-specific DE was explained by age-dependent DE in LEPs or MEPs at *adj. p* < 0.05. If we considered all genes with at least 2-fold change DE with age, these age-dependent directional changes accounted for only 21% of loss of lineage-specific expression events. These findings suggested that other mechanisms play a substantial role in regulating lineage fidelity, and that molecular changes associated with aging are not limited to stereotyped directional changes. Indeed, unsupervised hierarchical clustering of expression data only clustered samples by lineage (approximately unbiased, AU, *p* ∼ 100), but not by age, indicating that age-dependent expression changes were not robustly stereotypic (**Figure S3P**). In contrast, 43% of the losses in lineage-specific DM were explained by age-dependent DM in LEPs or MEPs at *adj. p* < 0.05. While these age-dependent changes explained less than half of the loss in lineage fidelity in methylation, unsupervised hierarchical clustering of DNAm data none-the-less robustly clustered samples by lineage (AU ∼ 100), and LEP samples by age (AU ∼ 100) (Figure 3K), showing that DNAm changes in LEPs are robust predictors of aging. The 18,528 CpG sites that were DM with age in LEPs and/or MEPs mapped to 5,903 genes – a number far larger than the number of genes that were DE with age. This suggested that in specific instances, DNAm changes with age may precede gene expression changes.

### Age-dependent directional changes in the luminal lineage are indicators of aging breast tissue

Because LEPs dominated the age-specific signal in the epithelia, we examined if the contribution of the luminal lineage could be detected in age-dependent changes observed in bulk tissue. We performed DE analysis on publicly available microarray expression data from normal primary tissue samples (GSE102088, m=114 subjects, 35 <30y, 68 >30y <55y, 11 >55y) that were generated for expression-DNAm correlation analysis by Song et al., [Song et al., 2017]. We detected a small number of DE genes with age (*limma adj. p* < 0.05), with 97 genes DE between younger <30Y and older >55y tissues, three genes DE between middle-aged >30y <55y and younger tissues, and no genes DE between middle-aged and older tissues. Of the 471 genes DE between younger and older LEPs, only the genome organizer *SATB1* (Figure 3Hii) was also DE and significantly down-regulated with age in bulk tissue (*adj. p 0.05*) (**Figure S3Qi**). Further, SATB1 showed significant LEP-specific expression compared to MEPs (adj. *p* < 0.001, fold change ≥ 2) (**Figure S3Qii**), and was also down-regulated in PAM50 LumA, LumB, Her2 relative to Basal and Normal intrinsic subtypes (Kruskal-Wallis, KW test adj. *p* < 0.0001) (**Figure S3Qiii**). Together, these results suggest that *SATB1*-mediated genome organization plays a key role in the maintenance of the luminal lineage and its dysregulation with age, and that genome reorganization in luminal-subtype cancers may track that of aged LEPs based on common downregulation of *SATB1*. We also considered DE genes that were nominally significant (*limma unadj. p* < 0.05) given the differences in platforms (RNA-seq vs. microarray), and found 87 genes significantly DE in LEPs and DE at nominal significance in bulk tissue, and 526 genes DE at nominal significance in both datasets. The majority of LEP age-dependent expression signals were more muted, but detectable, in bulk tissue.

To compare age-dependent DNAm changes in LEPs and bulk tissue, we used two independent published findings of age-dependent DM in normal primary breast tissue [Song et al., 2017, Johnson et al., 2017]. Song et al., found 1,214 CpGs DM with age (GSE101961, m=121 subjects), while Johnson et al. found 787 CpGs DM with age (GSE88883, m=100 subjects). Of these, 276 CpGs were commonly DM in the two independent datasets. Of the 18,522 CpG sites DM with age in LEPs (*adj. p 0.05*), 39 CpGs were also DM with age in the two bulk tissue datasets (Figures 3L). These 39 CpGs robustly stratified 95% younger <30y and 100% of older >55y samples from the Song et al., and Johnson data sets (approximately unbiased, AU, p ≥ 85); whereas the majority of middle-aged samples clustered with older tissues (**Figure S3Ri-ii**). Although detection of LEP age-dependent DNAm changes in bulk tissue was limited, there was sufficient signal to identify indicators of aging in breast tissue driven by DNAm changes in the luminal lineage.

### Aging-associated increase in variance contributes to loss of lineage fidelity

Next, we explored the alternate model that incorporated measures of variance as an explanation for the loss of lineage-specific expression in older epithelia (Figure 3Aii). Gene expression means (Figure 4A-B) and variances (Figure 4C-D) of LEPs and MEPs from younger cells were categorized into quantiles. Corresponding categories of gene expression means and variances in older cells relative to younger showed that gene expression means shifted minimally between younger and older cells (Figure 4A-B), whereas shifts in variances occurred at a much higher frequency (Figure 4C-D). Genes with very low or low variances in younger cells that had very high or high variances in older women were distributed across the range of gene expression levels in LEPs and MEPs (**Figure S4A-B**). Though the dynamic ranges of gene expression in LEPs and MEPs changed as a function of age, these changes were not stereotyped across individuals – i.e. different aged individuals had different sets of genes that deviated from the range of expression seen in younger samples. Indeed, principal component (PC) analyses showed that aged individuals did not cluster together but instead were scattered across the first two PCs, reflecting the contributions of genes with increases in variances with age to the principal axes (**Figure S4C-D**).

**Figure 4.**
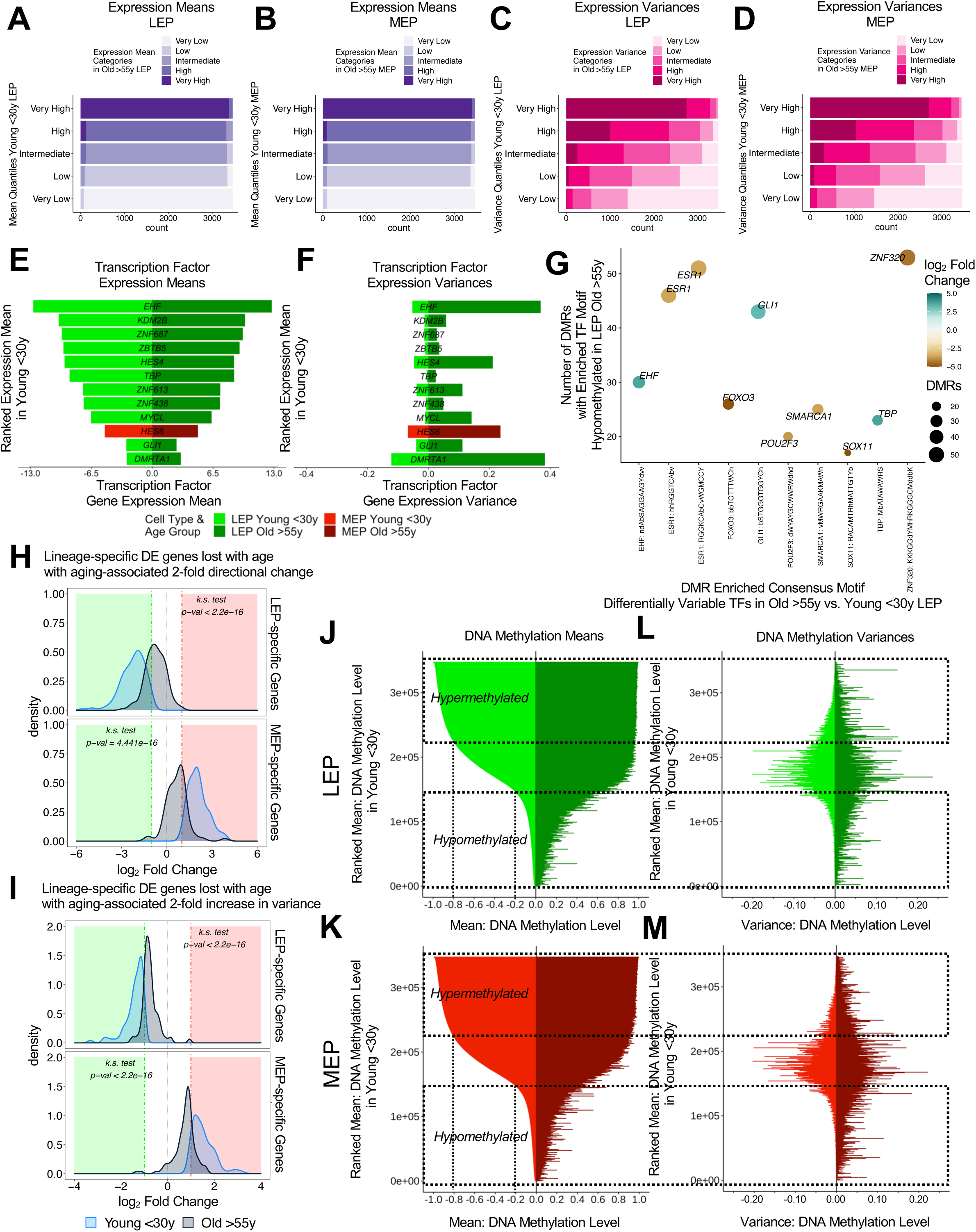
Aging-associated increase in variance contributes to loss of lineage fidelity. **(A-D**) Gene expression (**A, B**) means and (**C, D**) variances of LEPs and MEPs from younger women are categorized into quantile levels: very low, low, intermediate, high, very high. Corresponding categories of gene expression means and variances in older cells, as defined by threshold values in younger cells, are fractionally represented in color. (**E-F**) Gene expression (**E**) means and (**F**) variances of transcription factors (TFs) with significant increase in variances in older cells (*MDseq* differential variability (DV) *adj. p* < 0.05). (**G**) Number of DMRs (*DMRcate* FDR *p* < 0.05) that lost DNAm with age with enriched motifs (*ELMER* FDR *p* < 0.05) for DV TFs in LEPs (*adj. p* < 0.05). Point sizes reflect the number of DMRs with motif enrichment and are color coded based on DV lfc of the corresponding TFs. (**H-I**) Distribution of lfc in expression between LEPs and MEPs in younger (light blue) and older (blue gray) for either DE LEP-specific (top panel) or MEP-specific (bottom panel) genes that are lost with age (DE *adj. p* < 0.001, fold change ≥ 2) and that have at least an (**H**) age-dependent 2-fold directional change (*limma*) or (**I**) aging-associated 2-fold increase in variance (*MDSeq*) in the older cohort. LEP-specific genes are shown with (-) lfc values and MEP-specific-genes with (+) lfc values relative to each other; KS test *p*-values for equality of distributions of lfc between young and older women are annotated. (**J-K**) DNAm means of (**J**) LEPs and (**K**) MEPs from young (left) and older women (right) are shown in rank order from unmethylated to methylated CpG sites based on DNAm levels in the younger cohort. (**L-M**) Corresponding DNAm variances of (**L**) LEPs and (**M**) MEPs in younger (left) and older (right) women for the ranked ordered CpG sites. Unmethylated (beta values < 0.2) and methylated (beta values > 0.8) regions are indicated by dashed lines. See also **Figure S4**.

We performed gene expression differential variability (DV) analysis on the 14,601 genes whose variances could be estimated, and found 137 genes in LEPs and 128 genes in MEPs with significant age-dependent DV (*MDSeq adj. p* < 0.05) (**Figure S4E**). Twelve regulatory TFs in either LEPs or MEPs that had tuned windows of expression in younger cells were dysregulated in older cells through a significant increase in variance (*adj. p* < 0.05) (Figure 4E, 4F) and included *EHF*, *KDM2B*, *HES4, TBP*, *MYCL, GLI1, ZBTB5*, *DMRTA1*, and zinc fingers *ZNF687*, *ZNF613*, and *ZNF438* in LEPs, and *HES6* in MEPs. Notch target *HES4* was also DE with age. *GLI1* activates the hedgehog pathway in mammary stem cells [Bhateja et al., 2019]. Estrogen-regulated *HES6*, is known to enhance breast cancer cell proliferation [Hartman et al., 2009]. And lastly, *KDM2B (FBXL10)*, which is was down-regulated in a subset of older women (**Figure S4F**), is a histone demethylase ZF-CxxC protein that binds unmethylated DNA, and recruits polycomb repressor complex-1 to CpG islands and protects polycomb-bound promoters from *de novo* methylation [Farcas et al., 2012, Boulard et al., 2015]. Another 96 TFs had a 2-fold nominally significant increase in variance in LEPs and MEPs of older women (*unadj. p < 0.05*). We next mapped the motif enrichment of the 12 TFs that had age-dependent DV in LEPs (*adj. p* < 0.05) to the DMRs that lost methylation in older LEPs. Nine of these TFs were enriched in 334 hypomethylated DMRs (*ELMER* FDR *p* < 0.05) (Figure 4G), 246 of which overlapped gene promoter regions (**Figure S4G**). These analyses suggested that age-dependent variability in expression across individuals can lead to differential outcomes as different downstream targets are modulated in different individuals.

To understand how age-dependent variability changes affected lineage-specific expression, we focused on genes that lost lineage-specific expression with age and that showed at least a 2-fold increase in dispersions in the older cohort. Genes with 2-fold increases in variances with age explained 27% of the observed loss of lineage-specific expression events, on par with the proportion (21%) explained by genes that had 2-fold changes in DE (**Figure S4H**). Both of our models of directional and variant changes with age led to a significant decrease in the magnitude difference between LEP- and MEP-specific expression in the older cells (Figure 4H-I). Furthermore, 50% of genes with 2-fold age-dependent increase in variance and loss of lineage-specific expression were expressed at levels significantly correlated with methylation of at least one CpG site (cor. *adj. p* < 0.05) and those associations were predominantly anti-correlated in PPRs, suggesting that gene expression variances were impacted by DNAm.

As with expression, categorical shifts in mean DNAm levels represented a smaller fraction of changes between younger and older LEPs and MEPs (**Figure S4I, S4J**, respectively), whereas categorical shifts in variance were more pronounced (**Figure S4K, S4L**). In wine-glass plots, DNAm means and variances of LEPs (Figure 4I, 4K) and MEPs (Figure 4J, 4L) from younger (left) and older women (right) were shown in rank order from unmethylated to methylated CpG sites based on DNAm levels in the younger cohort. Partially-methylated regions showed high variances in both younger and older cells, whereas large shifts towards increased variances in older samples arose in the regions that were unmethylated and methylated in younger cells (Figure 4K, 4L). In gene region-specific analysis of CpG sites with very low and low variances in younger cells, the largest variances in older cells were largely restricted to unmethylated (beta values < 0.2) and methylated regions (beta values > 0.8), and were enriched at PPRs (**Figure S4Mi-ii**). Taken together, these data suggest that increased variances in transcription, as well as epigenetic regulation, are considerable drivers of the loss of lineage fidelity in breast epithelia. Such variances across individuals may underlie the age-dependent dysregulation of susceptibility-associated biological processes in specific individuals.

### Hallmark pathways associated with cancer are dysregulated with age in luminal and myoepithelial lineages

We next performed gene-set enrichment analysis to identify hallmark signatures of coherently-expressed gene sets that represented well-defined biological processes (Molecular Signatures DB) [Subramanian, et al., 2005, Liberzon et al., 2015] that were dysregulated with age. We found a number of biological hallmark gene sets enriched in older LEPs and MEPs that are known to be dysregulated in breast cancers (Figure 5A-B).

**Figure 5.**
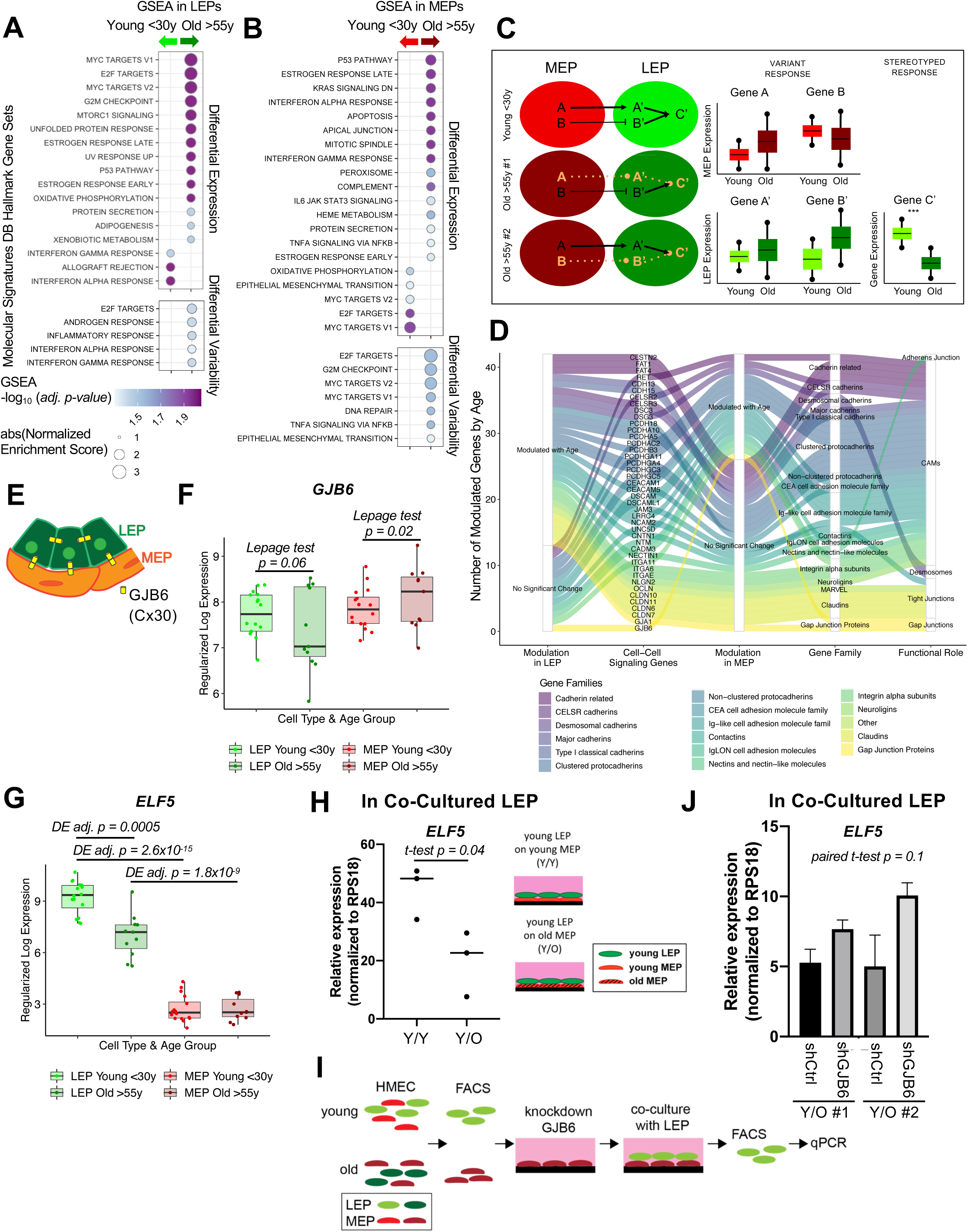
Hallmark pathways associated with cancer are dysregulated with age in luminal and myoepithelial lineages. (**A-B**) Molecular Signatures database (MSigDB) Hallmark gene sets identified by GSEA to be enriched (*fgsea adj. p < 0.05*) in younger and older (**A**) LEPs and (**B**) MEPs based on age-dependent DE analysis (top) (*limma adj. p* < 0.05) or DV analysis (bottom) (*MDSeq adj. p* < 0.05). Point colors show the GSEA -log_10_ of the adj. *p*-values, and point sizes indicate the absolute value of the GSEA normalized enrichment scores. (**C**) Schematic illustrating integration of directional and variant responses in older epithelial cells. Different genes become dysregulated in LEPs and MEPs of older individuals leading to an increase in variance in expression across aged cells. Through cell-cell signaling, variant responses in MEPs (gene A or gene B) can lead to variant responses in LEPs (gene A’ or gene B’). Where these variant responses integrate and affect common downstream genes in LEPs (gene C’) lead to detectable measures (***) of age-dependent directional changes that are seen as stereotyped responses in the lineage. (**D**) Modulation of apical junction-associated genes in LEPs and MEPs (Lepage test *p < 0.05).* Genes are annotated with their respective HUGO Gene Nomenclature Committee (HGNC) gene family and functional role in adherens junctions, cell adhesion molecules, desmosomes, tight junctions or gap junctions. (**E**) Schematic of Cx30 (*GJB6*) homotypic channel formation between LEPs and between LEP and MEP. (**F**) Boxplot of *GJB6* gene expression rlog values in LEPs and MEPs in young and older women. Lepage test *p*-values are indicated. (**G**) Boxplot of *ELF5* gene expression rlog values in LEPs and MEPs in young and older women. Adj. *p*-values for age-dependent DE in LEPs (*adj. p* < 0.05) and lineage-specific DE in younger and older cells (*adj. p* < 0.001, fold change ≥ 2) are indicated. (**H**) Relative expression of *ELF5* in young LEPs co-cultured with either young MEPs (Y/Y) or old MEPs (Y/O). Two-tailed t-test *p*-value indicated. (**I**) Schematic of co-culture methodology with HMEC cells from younger and older women enriched by FACS for LEPs and MEPs; *GJB6* is knocked-down in the older MEP feeder layer by shRNA; younger LEPs are co-cultured on top of the older MEP feeder layer for 10 days; LEPs are separated from MEPs by FACS; and LEP expression levels are measured by qPCR. (**J**) Relative expression of *ELF5* in young LEPs co-cultured with either shControl or shGJB6 older MEPs in two individuals. One-tailed paired t-test *p*-value indicated. See also **Figure S5**.

Seventeen hallmark gene sets were significantly modulated in LEPs (*fgsea adj. p* < 0.05) based on DE (Figure 5A top). Three immune-related gene sets were enriched in younger LEPs and included genes up-regulated in response to interferon IFN-alpha and -gamma, and during allograft rejection. In contrast, 14 gene sets were enriched in older LEPs, which included: two subgroups of genes regulated by MYC; genes encoding cell-cycle related targets of E2F TFs and involved in the G2/M checkpoint; genes upregulated by mTORC1 complex activation, during unfolded protein response, in response to UV radiation, and during adipocyte differentiation (adipogenesis); genes involved in the p53 and protein secretion pathways; genes defining early and late response to estrogen; and genes encoding proteins involved in oxidative phosphorylation and in processing of xenobiotics.

Twenty hallmark gene sets were significantly modulated in MEPs (*fgsea adj. p* < 0.05) based on DE (Figure 5B top). Five gene sets were enriched in younger MEPs, including: MYC targets and E2F targets; genes involved in oxidative phosphorylation; and genes defining epithelial-mesenchymal transition (EMT). In contrast, 15 gene sets were enriched in older MEPs, including: genes involved in p53 pathways; genes down-regulated by KRAS activation; genes mediating programmed cell death by caspase activation (apoptosis); immune-related gene sets upregulated in response to IFN-alpha, IFN-gamma and by IL-6 via STAT3, genes regulated by NF-kB in response to TNF and genes encoding components of the innate complement system; genes defining early and late response to estrogen; genes encoding components of peroxisome, apical junction complex, and mitotic spindle assembly; and genes involved in protein secretion and in metabolism of heme and erythroblast differentiation.

In addition, five gene sets were significantly modulated (*fgsea adj. p* < 0.05) based on DV and were enriched in older LEPs (Figure 5A bottom). These included: E2F targets that were similarly enriched via DE; genes defining responses to androgen and inflammation; and genes upregulated in response to IFN-alpha and IFN-gamma – gene sets that in contrast were enriched via DE in LEPs of younger women. Seven gene sets were significantly modulated (*fgsea adj. p* < 0.05) based on DV and enriched in older MEPs (Figure 5B bottom). These included: genes involved in DNA repair and G2/M checkpoint; genes regulated by NF-kB in response to TNF – a gene set similarly enriched via DE; as well as MYC targets, E2F targets, and genes defining EMT – gene sets that in contrast were enriched via DE in MEPs from younger women.

Several enriched gene sets were involved in processes that were disrupted with age either via DE or DV in both LEPs and MEPs, and such overlaps likely suggest integration of stereotyped and variant responses and reflect their impact in common biological processes (Figure 5C). Furthermore, the divergence in the age-dependent enrichment of cellular processes between DE and DV suggests that genes that become variable with age are associated with pathways that are otherwise important in maintaining lineage-specificity and function in younger cells.

### Gap Junction protein *GJB6* is a mediator of the non-cell autonomous mechanism of aging in breast

We showed previously that MEPs from older women can impose an aging phenotype on LEPs from younger women that acquire expression patterns of older LEPs when co-cultured in bilayers on the apical surfaces of MEPs [Miyano, Sayaman et al., Aging 2017]. This non-cell autonomous mechanism of aging requires direct cell-cell contact between LEPs and MEPs [Miyano, Sayaman et al., Aging 2017], and because changes in MEPs are predominantly associated with DV rather than DE, this suggests that LEPs serve as integration nodes for dysregulation in MEPs.

Apical junction-associated genes that play a key role in direct cell-cell interactions were significantly enriched with age in MEPs (Figure 5B). We explored dysregulation of known adherens, tight, and gap junctions, desmosomes, and cell adhesion molecules (CAM) in LEPs and MEPs to identify candidate genes that may regulate communication between the lineages. Because age-dependent changes involve both DE and DV, we performed a non-parametric Lepage test for location (central tendency) and scale (variability) on the 198 genes corresponding to cell-surface junction proteins, and found 42 genes that were modulated with age in LEPs and/or MEPs (*p* < 0.05) (Figure 5D). Genes encoding for junctional proteins that showed age-dependent modulation include tight junctions *CLDN10* and *CLDN11* (**Figure S5Ai-ii**), desmosomal cadherins *DSG3* (desmoglein) and *DSC3* (desmocollin) (**Figure S5Bi-ii**) – which are expressed in both LEPs and MEPs [Garrod and Chidgey, 2008], and gap junction *GJB6* (Connexin-30) – which is also expressed in both lineages in the normal mammary gland and is able to form homo-(LEP-LEP) and hetero-cellular (LEP-MEP) channels [Teleki et al., 2014] (Figure 5E). *GJB6* is of specific interest as it showed modulation via an increase in variance in older MEPs (*p* = 0.02) and nominal increase in variance in older LEPs (*p* = 0.06) (Figure 5F), and as such provide an avenue for exploring changes that occur in only a subset of older women that may lead to differential susceptibility across aged individuals.

We asked whether knockdown or inhibition of *GJB6* expression in the subset of older MEPs with higher expression relative to younger MEPs could restore proper signaling between LEPs and MEPs. To test this, we used our established heterochronous co-culture system and used recovery of LEP expression of *ELF5* as a readout. *ELF5* is a highly LEP-specific TF that was significantly down-regulated in LEPs with age (Figure 5G). LEP expression of *ELF5* was dynamic and responsive to microenvironment changes [Miyano, Sayaman et al., 2017], and was down-regulated in young LEPs when co-cultured on apical surfaces of older MEPs for 10 days (two-tailed t-test *p* = 0.04) (Figure 5H). If bringing variant *GJB6* under tighter control prevents chronologically older MEPs from imposing older biological ages in young LEPs, then *ELF5* levels should not decrease in co-culture. FACs-enriched LEPs from younger women were co-cultured for 10 days on older MEPs treated with either shGJBG or scramble shRNA (shCtrl) (**Figure S5C, 5I**). When co-cultured on top of older MEP-shGJB6 relative to MEP-shCtrl, LEP-expression of *ELF5* was maintained at higher levels (nominally significant one-tailed paired t-test *p* = 0.1) (Figure 5J), consistent with higher expression levels in younger women. LEP-expression of *ELF5* likewise showed a step-wise (though non-significant) increase when older MEP feeder layers were pre-treated with increasing concentrations of a non-specific gap junction inhibitor 18 alpha-glycyrrhetinic acid (18αGA) (**Figure S5D**). Thus, reducing the level or variance of *GJB6* prevented older MEPs from imposing an older biological age in young LEPs as determined by *ELF5* expression. These data suggest that variance is a driver of stereotypical aging phenotypes at the tissue level, and that constraining specific changes caused by an increase in molecular noise during aging (such as in cell-cell communication nodes may prevent the spread of age-related cues amongst epithelia.

### Common pathways between aging and susceptibility to breast cancer initiation

Aging increases the risk for breast cancer, thus we sought to understand how these gene expression and DNAm changes with age relate to cancer susceptibility. To do so, we examined LEPs from women considered to be at higher risk to develop breast cancer based on accepted genetic or clinical risk factors. We analyzed samples with mutations in *BRCA1*, *BRCA2*, *PALB2*, or *APC*, or samples that were normal-contralateral to or -peripheral to tumor (age range 25-72y, n=48 LEP and MEP samples, m=24 individuals, **Table S1**). We collectively refer to this cohort as higher risk samples and compared their LEP expression profiles with reduction mammoplasty samples from younger women <30y with average risk. We found 529 susceptibility-associated genes (inclusive of the common age-dependent and susceptibility-exclusive genes), including 45 TFs, that were DE between LEPs from higher risk vs. younger average risk women (*limma adj. p* < 0.05) (Figure 6A). Of these, 175 genes, including 24 TFs, were also DE with age in LEPs of average risk women (Figure 6A**, S6A**). These 175 common age-dependent and susceptibility-associated DE genes changed in the same direction (56% downregulated and 44% upregulated) in both older women and women with higher risk relative to young, and drove the clustering of all LEP samples from older and higher risk women (Figure 6B**, S6A**). Moreover, 93% of the common aging and higher risk transcriptomic changes shifted towards the direction of “MEP-like” expression. Of note, a sub-cluster of older and higher risk individuals had more pronounced “MEP-like” expression level shifts, underscoring how variability between individuals within the aged and higher risk cohorts could lead to more dysregulated phenotypes (Figure 6B). These results provide a basis for concluding that the epithelia of naturally aging average risk older women share susceptibility gene expression profiles with women who are clinically understood to have increased risk of breast cancer.

**Figure 6.**
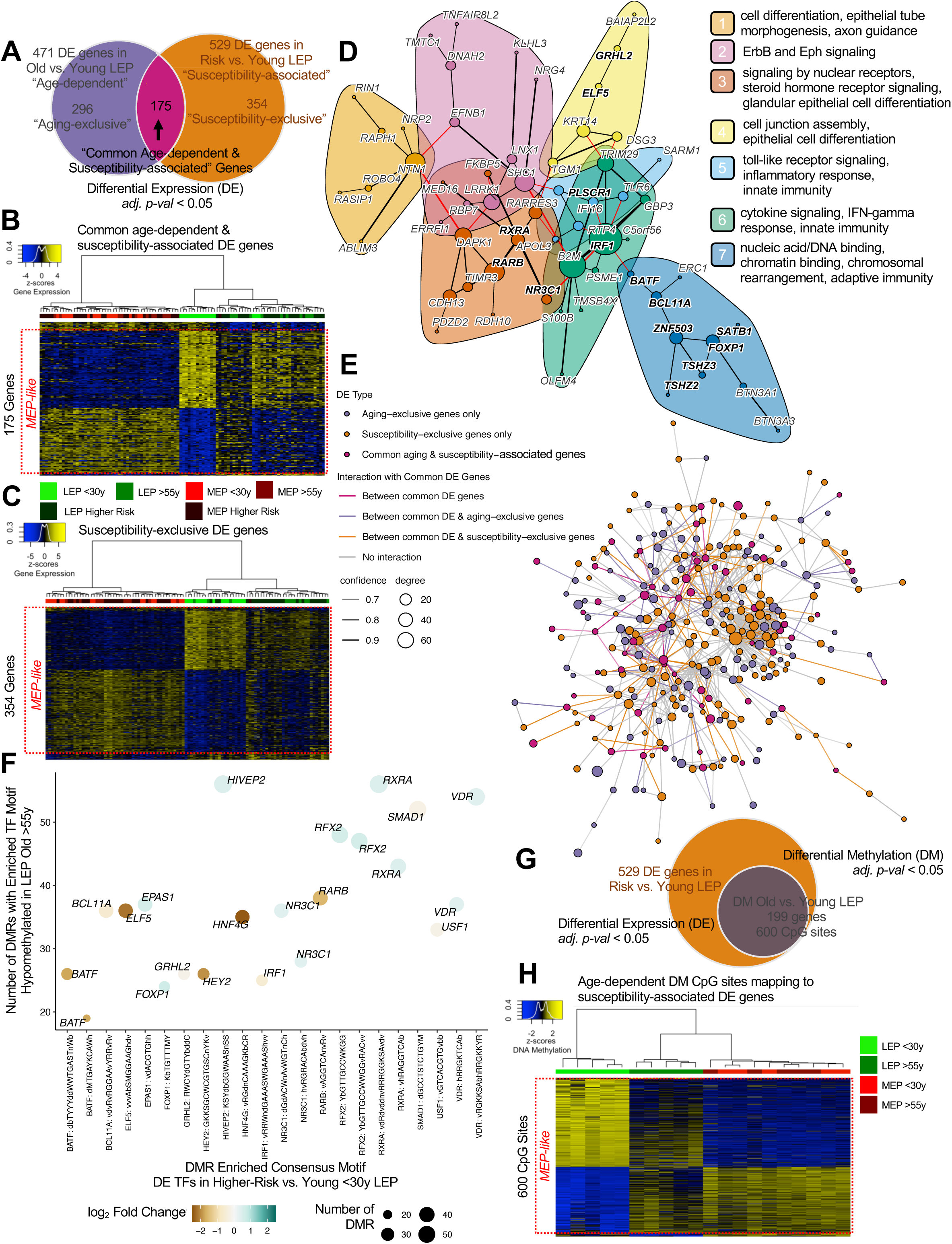
Age-dependent changes in methylation are priming events for increased susceptibility to breast cancer. (**A**) Venn diagram illustrating the intersection between age-dependent DE genes (*limma adj. p* < 0.05) identified from contrast of younger vs. older LEPs from average risk (AR) women, and susceptibility-associated DE genes (*limma adj. p* < 0.05) identified from contrast of younger LEPs from AR women vs. LEPs from women with higher risk (HR) of breast cancer. (**B-C**) Hierarchical clustering of all samples from younger AR, older AR and HR women based on (**B**) common age-dependent and susceptibility-associated DE genes; and (**C**) susceptibility-exclusive DE genes – e.g., DE only between younger AR and HR women but not DE by age. Scaled rlog values of DE genes are shown as a heatmap. Clustering performed using Euclidean distances and Ward agglomerative method. (**D**) Protein-protein interaction (PPI) network (*stringdb* interaction score ≥ 0.4, medium confidence) of common age-dependent and susceptibility-associated DE genes (*adj. p* < 0.05), with TFs annotated in bold. Seven gene communities are identified (*igraph* optimal community structure); corresponding network functional enrichment (*stringdb* enrichment FDR *p* < 0.05) of selected processes are annotated. (**E**) PPI network (*stringdb* interaction score ≥ 0.7, high confidence) between common age-dependent and susceptibility-associated DE genes (magenta), aging-exclusive DE genes (purple) and susceptibility-exclusive DE genes (orange) (*adj. p* < 0.05). (**F**) Number of DMRs (*DMRcate* FDR *p* < 0.05) that lost DNAm with age with enriched motifs (*ELMER* FDR *p* < 0.05) for susceptibility-associated DE TFs in LEPs (*adj. p* < 0.05). Point sizes reflect the number of DMRs with motif enrichment and are color coded based on DE lfc of the corresponding TFs. (**G**) Venn diagram illustrating the intersection between susceptibility-associated DE genes (*adj. p* < 0.05) and genes with age-dependent DM (*adj. p* < 0.05) at associated CpG sites. (**H**) Hierarchical clustering of samples from AR women based on age-dependent DM CpG sites mapping to susceptibility-associated DE genes (DE and DM *adj. p* < 0.05) – e.g., all genes DE between younger AR and HR women, that are also DM with age. Scaled beta values of DM CpG sites are shown as a heatmap. Clustering performed using Euclidean distances and Ward agglomerative method. See also **Figure S6**.

Of the 529 susceptibility-associated genes, 354 genes were uniquely DE between LEP of women with higher risk vs. younger average risk women (*adj. p* < 0.05) (Figure 6A). These genes defined a “susceptibility-exclusive” expression profile, and included 21 TFs, as well as the oncogenes *MET*, *EGFR* and *H3F3A* and the tumor suppressor gene *ATM* [Davoli et al., 2013] (**Figure S6B**). Hierarchical clustering based on these 354 susceptibility-exclusive genes – 41% of which are downregulated and 59% are upregulated in women with higher risk, clustered a subset of older women with the majority of higher risk women (Figure 6C**, S6B**), suggesting that some older women have acquired expression changes that are reflective of a susceptibility profile. Of these susceptibility-exclusive genes, 93% acquired “MEP-like” expression patterns, with LEP-specific genes downregulated and MEP-specific genes upregulated in LEPs from higher risk women (Figure 6C). Upon closer inspection of the expression patterns of these genes, we found genes that were significantly downregulated in higher risk women relative to young average risk women showed a trend of downregulation in a subset of older average risk women. In contrast, genes that were upregulated in women with higher risk relative to young average risk women showed largely no distinguishable increase in the older women (Figure 6C**, S6B**).

Common age-dependent and susceptibility-associated DE genes showed significant protein-protein interaction (PPI) network enrichment (*stringdb* PPI enrichment p = 4.3×10-7), including a highly connected 60-gene network that involved 14 DE TFs (*stringdb* interaction score ≥ 0.4, medium confidence) (**Figure S6C**). This network featured seven gene communities (*igraph* optimal community structure) anchored by: (1) NTN1, and functionally enriched for biological processes related to cell differentiation, epithelial tube morphogenesis and axon guidance; (2) *SHC1* and *LRRK1*, and functionally enriched for ErbB and ephrin signaling; (3) *DAPK1*, TF *RARB*, TF *RXRA* and *RARRES3*, and includes TF *NR3C1,* and functionally enriched for signaling by nuclear receptors, steroid hormone receptor signaling and glandular epithelial cell differentiation; (4) *KRT14*, and includes TFs *GRHL2* and *ELF5*, and functionally enriched for cell junction assembly and epithelial cell differentiation; (5) RTP4 and TF *PLSCR1*, and functionally enriched for toll-like receptor signaling, inflammatory response and innate immunity; (6) TF *IRF1*, *B2M* and *TRIM29*, and functionally enriched for cytokine signaling, IFN-gamma response and innate immunity; (7) TFs *FOXP1* and *ZNF503,* and includes TFs *BCL11A, TSHZ3, TSHZ2, BATF* and *SATB1*, and functionally enriched for nucleic acid/DNA-binding, chromatin binding, chromosomal rearrangement and adaptive immunity (Figure 6D) (*stringdb* network functional enrichment FDR *p* < 0.05). Community 4 includes *KRT14* and *ELF5*, genes we have previously validated to be age-dependent. Functional enrichment in community 7 supports our new findings that suggest a role for chromatin organizing genes. In addition to genome-organizer *SATB1*, community 7 member genes include: *BCL11A* which is a subunit of the BAF (SWI/SNF) chromatin remodelling complex [Kadoch et al., 2013]; zinc finger homeobox gene *TSHZ3* which has been shown to interactive with a BAF remodeling complex subunit [Faralli et al., 2011]; and *BATF* which is considered a pioneer factor [Ciofani et al., 2012] – factors that can bind condense chromatin allowing for accessibility of condensed regions, and is involved in recruitment of chromatin-organizing complexes in a CTCF-dependent manner [Pham et al., 2019]. Community 7 also included *BTN3A1* and *BTN3A3* which are MHC-I associated genes that play a role in T-cell activation [Blazquez et al., 2018], which along with enrichment of innate and inflammatory response genes in communities 5 and 6, further suggest a role for the immune response in aging and cancer-susceptibility.

Genes that were DE either with age only (“aging-exclusive”) or with higher cancer risk only (“susceptibility-exclusive”) were themselves highly connected via PPI (*stringdb* PPI enrichment *p* = 1.4×10^-6^) to the common age-dependent and susceptibility-associated DE genes. A main 739 gene network included 156 common age-dependent and susceptibility-associated DE genes that had 93 direct interactions with 265 aging-exclusive genes, and 152 direct interactions with the 318 susceptibility-exclusive genes (*stringdb* interaction score ≥ 0.7, high confidence) (Figure 6E), and featured functional enrichment for biological processes related to signaling by receptor tyrosine kinases, cell and tube differentiation and morphogenesis, localization to apical and cell leading edges, and components of the basement membrane and collagen-containing extracellular matrix (*stringdb* network functional enrichment FDR *p* < 0.05).

Age-dependent dysregulation of specific interactions and networks in LEPs shared common routes with biologies that are clinically understood to have a higher likelihood of a lifetime cancer diagnosis. Furthermore, the changes in LEPs that occurred in older and in higher-risk individuals involved acquisition of MEP-like expression patterns, suggesting that perturbation of pathways that maintain epithelial lineage differentiation is a key step towards increased cancer susceptibility. Enrichment of genes involved in DNA and chromatin-binding, and chromatin-remodeling, as well as involvement of genome-organizer *SATB1,* further suggests dysregulation in chromatin and genome organization mediate this process.

### Age-dependent changes in methylation are priming events for increased susceptibility to breast cancer

We next examined if the transition towards increased susceptibility could be mediated by DNAm changes present in aged epithelia in two ways. The first involved binding of hypomethylated DMRs in aged LEPs by TFs that were dysregulated in higher risk women. The second involved exploring if DE susceptibility-associated genes themselves, composed of both the susceptibility-exclusive and the common age-dependent and susceptibility genes shown in Figure 6A, have underlying DNAm changes already present in aged LEPs.

We identified 17 TFs DE between LEPs of higher risk and younger average risk women (*adj. p* < 0.05) that had enriched binding motifs in 57% of all DMRs that lost methylation in older LEPs (*ELMER* FDR *p* < 0.05) (Figure 6F). Seven of these TFs, including four that were susceptibility-exclusive (*EPAS1, HIVEP2*, *RFX2* and *VDR*), were upregulated in higher risk LEPs and were enriched in 276 DMRs hypomethylated with age that overlapped gene promoter regions (**Figure S6D**). This suggested that direct binding of these susceptibility-associated TFs to CpG sites that had lost DNAm with age, and the resulting dysregulation of downstream targets, could be a mechanism through which aging leads to increased cancer susceptibility.

Next, we examined if DE susceptibility-associated genes were primed for dysregulation through age-dependent DM. In the subset of 435 susceptibility-associated genes in LEPs that had matching expression and DNAm data, 199 genes (46%) mapping to 600 CpG sites had corresponding DM in at least one CpG site with age (*adj. p < 0.05*) (Figure 6G). Of these 199 genes, 58% had DM at their PPRs, 64% had DM at multiple CpG sites, and 57% had more than 10% of their assayed probes showing DM with age. Of the 600 age-dependent DM sites associated with this susceptibility expression profile, 97% acquired MEP-like methylation patterns (Figure 6H). Moreover, 97% of these CpGs were mapped to lineage-specific genes (adj. *p* < 0.05), with 54% mapping to highly lineage-specific genes (adj. *p* < 0.001, fold change ≥ 2). Considering only CpG sites that lost methylation with age and that were plausibly primed for transcriptional activation, 72% mapped to MEP-specific genes specifically at CpG-rich PPRs and gene bodies (**Figure S6E**). In contrast, 60% of CpG sites that gained methylation with age mapped to LEP-specific genes at open seas located in PPRs and gene bodies (**Figure S6E**).

Furthermore, of the 199 susceptibility-associated genes above, 120 were susceptibility-exclusive and mapped to 317 CpG sites that had corresponding DM in at least one CpG site with age (*adj. p < 0.05*) (**Figure S6F**). Thus, age-dependent DM may precede the significant changes in expression seen in higher risk LEPs. Indeed, we detected correlation (81% inverse correlation) (cor. *adj. p < 0.05*) between DNAm and expression levels of 120 DM CpG sites and 56 genes that were not considered DE with age, but DE with cancer risk (**Figure S6G**). This was consistent with our hierarchical clustering results that showed a majority of older LEPs cluster with higher risk LEPs based on the trends of their expression of susceptibility-exclusive genes (Figure 6C). We highlighted the DNAm levels of few of these susceptibility-exclusive genes that showed age-dependent DM, such as: *TRIM2*, *BAIAP2* and *PTK6* that were down-regulated in LEPs of higher risk women; and *TBCD*, *SLIT3*, *ADCY9*, TF *SKI*, *EGFR, FAT1, PLAT*, TF *SP6* and *MYLK* that were up-regulated in LEPs of higher risk women (**Figure S6H**). These genes showed subtype-specific expression in breast cancers (**Figure S6I**), and most have known roles in cancer progression. Overexpression of oncogene *EGFR* is associated with poor differentiation and larger tumor sizes, and early recurrence in early breast cancers [Masuda et al., 2012, Gonzalez-Conchas et al., 2018]; *SKI* inhibits TGF-beta/Smad signaling and suppresses the activity of *YAP* and *TAZ* in the Hippo signaling pathway [Rashidan et al., 2015]; *FAT1* loss increases *CDK6* mediated proliferation [Li et al., 2018]; and *MYLK* promotes proliferation and invasion of breast cancer cells, and is required for switching from quiescent to proliferative states in breast cancer cells [Zhou et al., 2008, Barkan et al., 2008].

Our results suggested that susceptibility is tied to age-dependent acquisition of MEP-like methylation patterns at specific regions that lead to opening/priming of MEP-specific PPRs and to silencing of LEP-specific PPRs in higher risk LEPs. Thus, a fraction of DNAm changes directly leads to changes in gene expression with age that correlate with higher risk, such as promoter methylation. Other transcriptomic changes may require secondary events, such as age-dependent variable TF expression and binding to unmethylated regions, or binding of DNAm regulatory proteins and chromatin remodeling complexes, to culminate in expression changes that underlies susceptibility.

### DNA methylation changes in early stage breast cancers are already present in aged luminal epithelia

That LEPs showed significant DNAm changes with age, whereas MEPs exhibited few, raised the possibility that DNAm changes presage age-associated breast cancer risk in a lineage-specific manner. Furthermore, we observed more genes with DM compared to genes that were DE, which raised the possibility that DNAm served as priming events that preceded gene expression changes and required secondary events to ultimately alter expression.

To further investigate this idea of DNAm changes as priming events, we analyzed a group of methylation changes in our aging cohort that are known to arise in early stage breast tumors. Titus et al., reported 19 differentially methylated gene regions (DMGRs) across 11 genes (*AGRN, C1orf170, FAM41C, FLJ39609, HES4, ISG15, KLHL17, NOC2L, PLEKHN1, SAMD11, WASH5P*) that were consistently DM in low-stage tumors in four PAM50 intrinsic breast cancer subtypes: LumA, LumB, Her2 and Basal (n=523) compared to normal-adjacent breast samples (n=124) in The Cancer Genome Atlas (TCGA) [Titus et al., 2017a]. 113 of the 150 CpG sites in our dataset that comprise these 19 DMGRs clustered LEP and MEP samples by lineage, with 14 CpGs passing the high significance threshold for lineage-specificity, eight of which lost lineage fidelity with age (adj. *p* < 0.001, fold change ≥ 2) (**Figure S7A**). LEPs were clustered according to age with six of these CpG sites showing significant age-dependent DM (adj. *p* < 0.05) (**Figure S7B-C**).

We next considered all the 76,847 DMGRs across all PAM50 intrinsic subtypes relative to normal-adjacent breast samples, and the 225,345 CpG sites associated with them [Titus et al., 2017a] that represent ∼50% of the probes in the Infinium 450K platform. Of these, 175,101 were analyzed in our dataset, and we found 22,806 comprised highly lineage-specific CpG sites (adj. *p* < 0.001, fold change ≥ 2), which represented 63% of all lineage-specific sites identified (**Figure S7A**). Further, 13,212 CpG sites lost lineage-specific DM with age, which represented 61% of loss of lineage fidelity events (**Figure S7A**). 11,030 CpG sites were associated with significant age-dependent changes in LEPs (*adj. p* < 0.05), which represented 60% of all age-dependent DM sites in the lineage, and another 9 were associated with age-dependent changes in MEPs (*adj. p* < 0.05) (**Figure S7B**). The high degree of overlap between lineage-specific and age-dependent DM CpG sites and breast cancer subtype DMGRs suggested that etiologies of breast cancers were differentially influenced by epithelial lineages.

Because older women have higher incidences of ER+ luminal-subtype cancers, we examined the set of cancer-associated DMGRs that were specific to PAM50 LumA and LumB subtypes compared to normal-adjacent samples [Titus et al., 2017a]. PAM50 non-basal subtypes were significantly more enriched for DMGRs (∼2000-fold) (LumA n=70,681 DMGRs, LumB n=66,720 and Her2 n= 53,231) than the Basal (n=26) intrinsic subtypes. We examined methylation levels of 62,427 shared luminal-subtype-specific DMGRs, representing 90% of all identified early-stage cancer DMGRs (PAM50 luminal-subtypes LumA and LumB). We identified 203,330 CpG sites associated with these DMGRs, of which 157,690 were analyzed in our dataset. Of those, 21,314 DMGR-associated CpG sites were highly lineage-specific (58% of all lineage-specific sites) (*adj. p* < 0.001, fold change ≥ 2); 12,317 lost lineage-specific DM in the older cohort (57% of all sites that lost lineage fidelity with age) (**Figure S7A**). Furthermore, 10,259 CpG sites had significant age-dependent changes in DNAm in LEPs (55% of age-specific sites in LEP) (*adj. p* < 0.05), and 9 were DM with age in MEPs (*adj. p* < 0.05) (**Figure S7B**). Hierarchical clustering of samples based on the 10,263 age-dependent CpG sites associated with luminal-subtype DMGRs recapitulated the observed phenomenon whereby older LEPs cluster with MEPs (Figure 7A). Thus, methylation patterns dysregulated in luminal-subtype cancers were specifically dysregulated in LEPs, leading to loss of lineage fidelity and acquisition of more MEP-like methylation patterns. Taken together, a significant fraction of methylation changes previously ascribed to the earliest stages of breast cancer were already present in older epithelia. Indeed, they represented the majority of the observed changes in DNAm with age in the LEP lineage, which is consistent with the high incidence of luminal-type cancers in older women.

**Figure 7.**
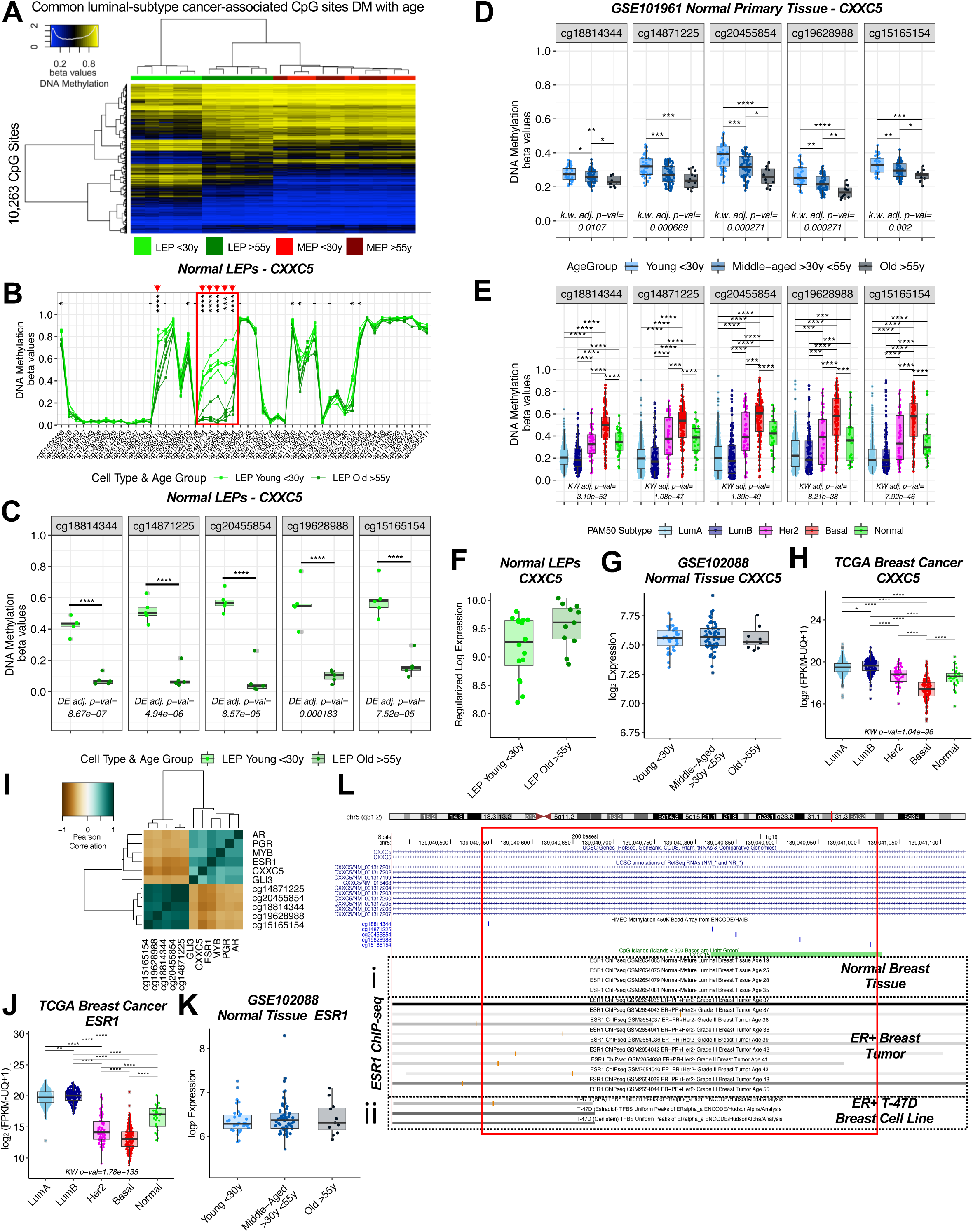
Aging-associated hypomethylation of TF-binding sites in *CXXC5* primes epithelia for *CXXC5* dysregulation in luminal-subtype cancers. (**A**) Hierarchical clustering of LEP and MEP samples based on age-dependent DM CpG sites (*limma adj. p* < 0.05) mapping to common DMGRs identified by Titus et al., associated with PAM50 LumA and LumB intrinsic-subtype breast cancers [Titus et al., 2017a]. DNAm beta values of DM CpG sites are shown as a heatmap. Clustering performed using Euclidean distances and Ward agglomerative method. (**B**) *CXXC5* DNAm beta values in LEPs from younger and older women. Age-dependent DM adj. *p*-value significance levels (*limma*) indicated (* < 0.05, ** < 0.01, *** < 0.001, **** < 0.0001). Red box highlights the *CXXC5* 5-CpG DMR (*DMRcate* FDR *p* < 0.05) composed of five contiguous DM *CXXC5* CpG sites: *cg18814344, cg14871225, cg20455854, cg19628988, cg1516515* located at the PPR. Red arrows indicate CpGs that are DM with age in LEPs with lfc > 1.5 (*adj. p* < 0.05). (**C-E**) Boxplots of DNAm beta values at the five contiguous *CXXC5* DM CpG sites in the age-dependent DMR (FDR *p* < 0.05) (**C**) in LEPs in younger <30Y and older >55y women; (**D**) in an independent normal primary breast tissue data set, GSE101961 [Song et al., 2017] across age groups: younger <30y, middle-aged >30y >55y and older >55y; and (**E**) in the TCGA breast cancer cohort across PAM50 intrinsic cancer subtypes: LumA, LumB, Her2, Basal and Normal. Age-dependent DM adj. *p*-value and significance levels in LEPs (*limma*); and KW test BH-adj. *p*-values across CpG sites between age groups in normal primary tissue and between cancer subtypes in TCGA, and post-hoc pair-wise Wilcoxon test adj. p-value significance levels are annotated (* < 0.05, ** < 0.01, *** < 0.001, **** < 0.0001). (**F-H**) Boxplots of gene expression values of *CXXC5* in LEPs from younger and older women; (**G**) in normal primary breast tissue from an independent data set, GSE102088 [Song et al., 2017] across age groups, and (**H**) in the TCGA breast cancer cohort across cancer subtypes. No significant age-dependent DE in LEPs and normal primary tissue (*limma*) detected. KW test *p*-value between cancer subtypes in TCGA and post-hoc pair-wise Wilcoxon test adj. *p*-value significance levels are annotated (* < 0.05, ** < 0.01, *** < 0.001, **** < 0.0001). (**I**) Heatmap of Pearson correlations between *CXXC5* gene expression, *CXXC5* DNAm at the five contiguous DM CpG sites, and expression of highly correlated TFs *AR*, *PGR*, *ESR1*, *MYB* and *GLI3* (Pearson abs(cor) > 0.5) with enriched motifs (*ELMER* FDR *p* < 0.05) at the *CXXC5* 5-CpG DMR in the TCGA cohort. Clustering performed using complete agglomerative method with 1-correlation as distance metric. (**J-K**) Boxplots of gene expression values of *ESR1* (**J**) in the TCGA breast cancer cohort across cancer subtypes; or (**K**) in normal primary breast tissue, GSE102088 [Song et al., 2017] across age groups. No significant age-dependent DE in normal primary tissue (*limma*) detected. KW test *p*-value between cancer subtypes in TCGA and post-hoc pair-wise Wilcoxon test adj. *p*-value significance levels are annotated (* < 0.05, ** < 0.01, *** < 0.001, **** < 0.0001). (**L**) Publicly available *ESR1* ChIP-seq peaks from (**i**) normal breast tissues and ER+ breast tumors, GSE99680 [Chi et al., 2019]; and (**ii**) T-47D ER+ breast cancer cell line [ENCODE Project Consortium, 2011, 2012] overlapping the *CXXC5* 5-CpG DMR (FDR *p* < 0.05) displayed on a zoomed region on the UCSC Genome Browser. See also **Figure S7**.

### DNAm regulatory proteins are primed for dysregulation via aging-associated DNAm changes

Because DNAm changes are more robust indicators of aging in LEPs and indicative of the genome-wide dysregulation of DNAm with age, we looked for significant age-dependent changes in expression or methylation levels of genes that encode for proteins associated with regulation of DNAm. The specific classes of proteins evaluated were: (1) DNA methyltransferases, (2) DNA demethylases ten-eleven-translocation enzymes, (3) zinc finger ZF-CxxC domain-containing proteins that bind unmethylated DNA [Long et al., 2013], and (4) proteins that bind methylated DNA including Methyl-CpG binding proteins (MBDs) and methyl-CpG binding zinc-finger Kaiso family members [Bogdanović et al., 2009]. Expression levels of genes in these classes did not significantly change with age, but DNAm levels of at least one CpG site was altered in 10 of 15 of genes in older LEPs (*adj. p* < 0.05) including DNA methyltransferase *DNMT3A* that is involved in *de novo* methylation [Okano and Li, 1998] and associated *DNMT3L* that stimulates activity of *DNMT3A* and *DNMT3B* [Suetake et al., 2004], DNA demethylase *TET1* [Pastor et al., 2013], ZF-CxxC family members *CXXC1, CXXC5, FBXL19, KDM2A, KDM2B,* and Kaiso family members *ZBTB4* and *ZBTB38* (**Figure S7D**). All 45 CpG sites in these 10 genes acquired MEP-like DNAm patterns in older LEPs (**Figure S7Ei-x and S7F**), which suggested methylation changes in those key DNAm-regulatory genes were non-random. 71% of those CpG sites showed decreased DNAm levels with age. Of specific interest to us were DNAm-regulatory genes with DM sites that represent a large fraction of all associated probes (>10%) (**Figure S7D, S7Ei-x**). *CXXC5*, *DNMT3A* and *KDM2B*, showed the largest magnitude changes in DM with age occurred at CpG islands or CpG shores (*adj. p* < 0.05, lfc > 1.5) (**Figure S7G**).

### Aging-associated hypomethylation of TF-binding sites in *CXXC5* primes epithelia for *CXXC5* dysregulation in luminal type cancers

As a case study, we focused on *CXXC5* DNAm in LEPs. Five contiguous *CXXC5* CpG sites in an age-dependent DMR (*DMRcate* FDR *p* < 0.05): *cg18814344, cg14871225, cg20455854, cg19628988,* and *cg15165154*, spanning 3,388 bp underwent hypomethylation events in older LEPs at a PPR annotated to a CpG island (Figure 7B red box, **7C**). Hypomethylation of this CpG island region in older LEPs was confirmed in our WGBS dataset [Senapati et al., 2020] (**Figure S7Hi**). This 5-CpG DMR was located in the first intron (**Figure S7Hi**), a region that has been shown to have inverse correlation between intron methylation and gene expression [Anastasiadi et al., 2018]. The contributions of the LEP lineage to age-dependent DM of *CXXC5* in bulk normal primary tissue were detectable in two independent datasets that included young <30y, middle-aged >30y <55Y and older >55y women (KW adj. *p* < 0.05-0.0001, lfc ≥ 1.5, red arrows): GSE101961 (m=121 subjects) [Song et al., 2017] and GSE88883 (m=100) [Johnson et al., 2017], and were most clearly seen at the five contiguous CpG sites (**Figure S7I, S7J**; red box). Post-hoc pair-wise comparisons (pwc) showed highly significant DM between younger <30Y and older >55y women in the 5-CpG DMR (Wilcoxon test *adj. p* < 0.05-0.0001) (Figure 7D**, S7K**). In contrast, the contributions of the LEP lineage to age-dependent DM in bulk normal primary tissues for DNMT3A, DNMT3L, TET1 and KDM2B were less distinguishable (KW adj. *p* < 0.05-0.0001, lfc ≥ 1.5, red arrows) (**Figure S7Li-iv**). Moreover, this 5-CpG DMR of CXXC5 stratified PAM50 breast cancer subtypes in women in The Cancer Genome Atlas (TCGA) cohort (n=781 samples, m=772 subjects with PAM50 annotation) (KW *adj. p* < 0.0001) (Figure 7E**, S7M**), consistent with its role as one of the PAM50-gene intrinsic subtype predictors [Parker et al., 2009]. This 5-CpG DMR was methylated in LumA and LumB subtypes at levels (mean beta=0.22 and 0.23, respectively) comparable to older LEPs and to MEPs; methylated in Normal-like and HER2 subtypes at levels (mean beta=0.36 and 0.37, respectively) comparable to normal bulk-tissue samples; and methylated in Basal subtypes at levels (mean beta=0.53) comparable to young LEPs (Figure 7E**, S7M**). Post-hoc comparisons were highly significant between all subtypes in the 5-CpG DMR (Wilcoxon *adj. p* < 0.001-0.0001) (Figure 7E). Other gene regions of *CXXC5* were also DM across PAM50 subtypes that were not DM with age in normal LEPs or normal tissue, including a region that was unmethylated in normal LEPs and normal tissue downstream of the 5-CpG DMR, (**Figure S7M**, red dotted box). Methylation levels of the *CXXC5* CpG sites in tumors were more variable than in bulk normal primary tissue, which possibly reflected varying degrees of *CXXC5* dysregulation across individuals with cancer.

DM events in *CXXC5* were significant, whereas we did not detect significant changes with age in expression of *CXXC5* in LEPs (Figure 7F) nor in bulk normal primary tissues (GSE102088 m=114) [Song et al., 2017] (Figure 7G). This led us to hypothesize that hypomethylation events required additional binding of TF at DM regions to lead to gene expression changes. Indeed, *CXXC5* was significantly DE in women across PAM50 breast cancer subtypes in TCGA (KW *adj. p* < 0.0001) (n=1,089 samples, m=1,078 subjects with PAM50 annotation) where it was highly expressed in LumA and LumB subtypes and downregulated in basal subtypes (Figure 7H), suggesting that TF-binding of the unmethylated PPR CpG island in the 5-CpG DMR may be the activating event to increase in *CXXC5* expression in luminal-subtype cancers.

Motif analysis of the PPR at the 5-CpG DMR identified 63 TF motifs that were enriched (*ELMER* FDR *p* < 0.05) in the region +/-250bp of the given CgG sites (**Figure S7N**). To identify candidate regulators of *CXXC5* expression, we performed a correlation analysis between expression levels of the selected TFs and *CXXC5,* and methylation levels at the *CXXC5* 5-CpG DMR within the TCGA cohort with matched expression and methylation data across PAM50 subtypes (m=770 subjects with PAM50 annotation) (**Figure S7O**). Five TFs were correlated with *CXXC5* expression and anti-correlated with *CXXC5* 5-CpG site methylation (Pearson abs(cor) > 0.5): *ESR1, GLI3, AR, MYB* and *PGR. ESR1* had the strongest correlation with *CXXC5* expression (cor=0.66) and the strongest anti-correlation to the *CXXC5* 5-CpG site methylation (cor=-0.64) (Figure 7I). *ESR1, GLI3, AR, MYB* and *PGR* were all significantly upregulated in luminal-subtypes (KW *adj. val* < 0.0001) (Figure 7J**, S7Pi-iv**). All five TFs were not DE with age in normal primary tissue, which was consistent with *CXXC5* not being upregulated with age in normal tissue despite aging-associated hypomethylation of *CXXC5* (Figure 7K, **S7Qi-iv**). We validated *ESR1* binding to this 5-CpG DMR of *CXXC5* via publicly available ChIP-seq data (GSE99680) that included 10 ER+ breast tumor samples (age range: 37-55y) and four ER+ mature luminal epithelia from normal breast (age range: 19-35y) [Chi et al., 2019]. In 9 out of 10 ER+ breast tumors, ESR1 ChIP-seq peaks overlapped at least one of the *CXXC5* five CpG sites, and in the majority of samples, ESR1 peaks spanned the entire region. In contrast, no ESR1 ChIP-seq peaks overlapped the same region in normal ER+ luminal epithelia (Figure 7Li, 7Hii). ENCODE *ESR1* ChIP-seq data for ER+ T-47D breast cancer cell line treated with estradiol, genestein – a phytoestrogen, and bisphenol A (BPA) – an estrogen mimicking compound, likewise showed overlap of ESR1 peaks with the first CpG site in the 5-CpG DMR of *CXXC5* [ENCODE Project Consortium, 2011, 2012] (Figure 7Lii, 7Hiii). These results suggested that *CXXC5* is epigenetically primed during aging, but large-scale dysregulation of its expression occurred during cancer progression concomitant with dysregulation of expressed TFs that bind the primed CpG sites. In luminal-subtype cancers, very high expression levels of *ESR1* may out-compete other binding proteins to become the dominant regulator of *CXXC5*.

## Discussion

Using lineage-specific analyses, we have shown that aging involves integration of stereotyped and variant responses that reshape the transcriptomic and epigenomic landscapes of the two main epithelial lineages of the breast, luminal epithelial cells (LEPs) and myoepthelial cells (MEPs). These changes lead to a loss in lineage fidelity with age where faithfulness of lineage-specific expression is diminished – encapsulating the genome-wide observed loss of tuned windows of expression of lineage-specific genes and a decrease in the magnitude of differential expression between the genes that define LEPs and MEPs. Through our approach, we dissected the contribution of each epithelial lineage and identified two models mediating this loss of lineage fidelity in breast epithelia with age. Either via directional changes, as measured by differential gene expression; or via an increase in variance, as measured by differential variability analysis. Aging-associated increases in expression variances occur in both epithelial lineages, whereas we showed that the overwhelming majority of directional gene expression and DNAm changes with age occurred in LEPs and not in MEPs. This is a striking finding when one considers the two lineages reside juxtaposed in tissue and that they arise from common bipotent progenitors. Moreover, the vast majority of directional changes in LEPs led to acquisition of expression and DNAm patterns that are MEP-like. We hypothesize that LEPs are the nexus of integration for variant responses that lead to stereotyped age-dependent outcomes – the likely result of a dynamic process of iterative feedback between LEPs and MEPs, and other cell types in the breast. Age-dependent changes in epithelia were associated with enrichment of biological processes dysregulated in breast cancers. We discovered a core subset of genes with similar expression and methylation profiles in older LEPs and higher risk LEPs that were highly connected through well-defined protein-protein interactions, which intricately linked a cancer susceptibility profile directly with aging. In particular, age-dependent DNAm changes that led to hypomethylation and increased accessibility of gene-regulatory regions may lead to “primed” DNAm states in older women. Considered in the context of increased biological variability across the older individuals, we hypothesize variable TF expression and binding of these primed hypomethylated DNAm states leads to the divergent dysregulation of these susceptibility-associated genes and provides a plausible mechanism for the observed differential susceptibility to cancer initiation in older individuals.

DNAm is a key factor regulating the metastability of the observed loss of lineage fidelity in expression. Lineage-specific gene expression was anti-correlated with methylation of corresponding CpG sites at promoter proximal and gene regulatory regions including CTCF binding sites. Furthermore, we found concordant loss of lineage-specific expression and loss of differential methylation at these CpG sites with age. While CTCF did not show age-dependent expression, we found a large fraction (60%) of CpG sites that overlapped CTCF binding sites that lost lineage-specific DM with age. This finding was striking because CTCF binding is thought to be more cell type invariant than most other DNA binding proteins [Kim et al., 2007]. Loss of lineage-specific DM at CTCF binding sites was associated with gained accessibility to CpGs in aged LEPs at sites that were unmethylated in MEPs of younger women; and loss of accessibility of CpGs in aged LEPs at sites that were unmethylated in LEPs of younger women. CTCF is involved in higher-order chromatin conformation and has insulator properties that demarcate regulatory regions from promoters and serve as chromatin boundary elements [Gaszner and Felsenfeld, 2006]. Loss of lineage-specific DM at CTCF binding sites with age may mediate the concerted genome-wide loss of lineage-specific expression.

Aging studies that examined gene expression and DNAm in human tissues have been largely restricted to analysis of bulk tissue, lacked cell-type specific resolution, and were focused on directional changes with age using DE and DM analyses. Bulk analyses make it impossible to separate impacts of aging that are driven by the intrinsic changes that occur molecularly within each lineage versus compositional changes that reflect shifts in cell type proportions. Lineage-specific analyses provide intermediate resolution between bulk RNA-seq and single-cell RNA-seq and allows for cost effective analysis of cell population-level responses and interactions. As such, lineage-resolution analyses also provide an avenue to validate computational deconvolution methods that have emerged to extract cell-type specific contributions in bulk tissue [Shen-Orr and Gaujoux, 2013; Titus et al., 2017b]. We provided evidence that tissue-level changes with age are driven not just by changing compositions of the breast [Garbe et al., 2012, Benz et al., 2008], but by intrinsic molecular changes in the underlying cell populations. Indeed, we found gene expression of genome organizer *SATB1* to be significantly down-regulated both in LEPs and in normal primary breast tissue with age, along with 87 other genes that showed significant DE in LEPs and nominally significant DE in primary tissue despite the difference in platforms (RNA-seq vs. microarray). In addition, we found 39 CpGs DM with age in LEPs were also reported by two independent studies to be DM in normal primary breast tissue with age [Song et al., 2017, Johnson et al., 2017]. These 39 CpGs robustly stratified breast tissue samples with age and should be considered as potential biomarkers of epithelia that are susceptible to cancer initiation. While detection of age-dependent changes in bulk tissue was limited, there was sufficient signal to identify potential biomarkers of aging breast tissue that was driven by the underlying molecular changes in the luminal lineage.

LEPs from older women lose lineage fidelity and acquire MEP-like expression patterns, but still maintain a significant fraction of their LEP-specific features. This is consistent with phenotypes we previously reported that showed LEPs acquiring expression of proteins that were otherwise expressed in young MEPs. As an example, we previously showed that LEPs gained expression of K14 and the YAP TF known to drive basal and EMT-related gene programs while maintaining LEP-specific expression of K19 [Garbe et al., 2012; Pelissier et al., 2014]. Similarly, we previously reported decreased expression of LEP-specific TF *ELF5* in older women [Miyano, Sayaman et al, 2017]. Such changes may reflect a loss luminal specification, and co-expression of luminal and basal (myoepithelial) markers indicative of an altered differentiation program. For instance, K19+/K14+ phenotype was associated with epithelial bipotent progenitors in the breast [Villadsen et al., 2007], and *ELF5* plays a role in regulating cell fate and progenitor function. Indeed, mammary glands of Elf5-null mice accumulate cell types with dual luminal/basal properties such as K8+/K14+ co-expression that are thought to represent the blocked differentiation of CD61+ luminal progenitors that increase in abundance with Elf5 deletion [Chakrabarti et al. 2011, Oakes et al., 2008]. We present further evidence of multiple examples of aging changes in LEPs that likely impinge upon the maintenance of the differentiated luminal state. Up-regulation in older LEPs of MEP-specific TF *ZNF503* that acts as an inhibitor of *GATA3*, which plays an essential role in maintenance of luminal epithelial differentiation [Shahi et al., 2017, Kouros-Mehr et al., 2006]. Down-regulation in older LEPs of TF *HES4*, a canonical target of Notch1 that plays an important role in normal breast epithelial differentiation and cancer development [Kontomanolis et al., 2018]. Down-regulation in older LEPs of LEP-specific TFs *GRHL2* and *SGSM2* that are associated with EMT in mammary epithelial cells and the loss of potent tumor-suppressor, E-cadherin, that plays a key role in the maintenance of differentiated epithelial states [Xiang et al., 2012, Lin et al., 2019, Berx and Van Roy, 2001]. Furthermore, we observed acquisition of MEP-like expression patterns in LEPs of women who are clinically considered to have increased risk of breast cancer. Those MEP-like changes were detected both in the subset of genes that were commonly DE between older and higher-risk women relative to younger average-risk women, and in the subset of genes that were DE in higher-risk women only. This suggests that susceptibility entails loss of proper specification of the luminal lineage, and that molecular changes in LEPs contributes to this loss. Some older women showed much larger shifts in expression that were equivalent to those in higher-risk women, underscoring how variability across aged individuals may lead to differential susceptibility. Prat and Perou proposed that, based on differentiation predictor scores and genomic characteristics, invasive breast cancer subtypes could be placed along a hierarchy of normal luminal epithelial differentiation with luminal-subtype cancers being closest to mature luminal cells [Prat and Perou, 2011]. In the context of these findings, we speculate that the aging changes in LEPs that we are reporting are signs of reprogramming and de-differentiation, or incomplete differentiation.

The lineage bias of age-dependent DE and DM suggests that epithelial lineages age via different mechanisms. The Horvath pan-tissue epigenetic clock was shown to be poorly calibrated in whole breast tissue [Horvath, 2013], and the incongruity may be due to lineage-specific differences and fractional changes of the lineages with age. Based on Horvath’s pan-tissue clock, our data shows that MEPs are shifted by as much as a decade “older” than matched LEPs. In the context of our earlier findings showing that older MEPs can impose aged phenotypes on younger LEPs [Miyano, Sayaman et al, 2017], the differential biological ages of the lineages may point to the ability of MEPs to propel LEPs through the aging process. MEPs are surrounded by an epithelial basement membrane (BM) that separates the epithelia from the stroma. MEPs showed significantly higher expression of *DNMT1*, which is primarily involved in maintenance of DNAm, and significantly lower expression of TET DNA demethylases compared to LEPs. We speculate that the patterns of expression of these epigenetic enzymes, together with the need for MEPs to be responsive to the dynamic and reciprocal signaling from the epithelial basement membrane and stromal microenvironments [Bissell and Aggeler, 1987], mitigates accumulation of DNAm changes in MEPs. Thus, accompanying gene expression changes would not be expected to occur in this lineage at the same rates as in LEPs. LEPs have shorter telomeres than matched MEPs, indicating that they undergo relatively more proliferation [Sputova et al., 2013, Meeker et al., 2004]. Older LEPs lose expression of *ZNF827*, a TF that promotes telomere elongation and homeostasis through recruitment of the nucleosome-remodeling and histone-deacetylation complex, and DNA repair proteins [Vilas et al., 2018], further highlighting instability in this lineage with age. Along with higher expression of TET proteins, and lower expression of *DNMT1* compared to MEPs, this could explain why DNAm changes and accompanying gene expression changes occur more frequently in the LEP lineage and why this lineage appears to be the integration point of age-dependent directional changes.

The striking phenotypic changes in LEPs are starkly juxtaposed to MEPs, which have so far revealed few obvious signs of changes with age. Nevertheless, heterochronous bilayers of MEPs and LEPs suggest that the chronological age of MEPs controls the biological age of LEPs, illustrating that MEPs do change with age and revealing the existence of a non-cell autonomous mechanism that integrates aging-imposed damage across the tissue [Miyano, Sayaman et al, 2017]. Here, we showed that changes in MEPs largely involved changes in gene expression and DNAm variances with age that were also observed in LEPs. Aging-associated increases in variances in both lineages drove a large fraction of the observed loss of lineage fidelity with age, comparable to the contribution of age-dependent directional changes. In our opinion, changes in variance is an underappreciated component of aging analyses. DE and DM analyses are ubiquitous statistical tools used in the analysis of expression and DNAm profiling studies, whereas only more recently have changes in variance been systematically analyzed [Xie et al., 2011, Slieker et al., 2016, de Jong, et al., 2019, Bashkeel, et al., 2019], along with the development of differential variance analytical tools [Phipson and Oshlack, 2014, Ran and Daye, 2017]. Indeed, for changes to be detected as significant in DE and DM analyses, the following assumptions must be met: (i) the biological phenomenon causes dysregulation that is directional – e.g., genes are either up- or down-regulated; and (ii) dysregulation occurs at the same time – e.g., in the same genes in the same pattern, across the majority of individuals in the group of interest – i.e., it is stereotypic. While some aging processes may be deterministic, like telomere shortening, other processes may be stochastic, born out of the accumulation of random physicochemical insults that manifest as an increase in noise in the system [Todhunter, Sayaman et al., 2018]. In the latter case, the signal itself is the noise in the system. Another way to view this type of dysregulation is by observing the deviation from a set range. A change in the dynamic range of expression, for instance, of regulatory genes that have very tuned or narrow windows of expression, can lead to dysregulation as expression deviates from the set range. This noise can lead to decoupling of tightly regulated networks. While both increase and decrease in dynamic range with age do occur, here we specifically focused on increases in variance and its effect on the loss of lineage fidelity with age. Accordingly, in single-cell studies, aged cells were shown to have increased transcriptional variability and loss of transcriptional coordination compared to younger cells of the same tissue [Kowalczyk et al., 2015, Martinez-Jimenez et al., 2017, Enge et al., 2017, Levy et al., 2020], suggesting that increase in cellular heterogeneity with age underlies the population-level increases in variances between the individuals we observed. As such, the molecular signals of aging cells may not be fully captured as stereotyped directional changes – rather a large fraction of age-associated changes will be reflected as increases in measured variance in the molecular signal across an aged cohort.

Regulatory factors like TFs that have very tightly tuned windows of expression in young healthy individuals and that see large increases in variance in older subjects, are good candidates for susceptibility factors that could be predictive of breast cancer risk. Several TFs that showed significant increases in variances in older women include luminal-specific TF *EHF*, canonical Notch target gene *HES4,* proto-oncogene *MYCL, GLI1*, a hedgehog pathway activator in mammary stem cells, and histone demethylase *KDM2B* in LEPs, and estrogen regulated TF *HES6* in MEPs [Kontomanolis et al., 2018, Bhateja et al., 2019, Hartman et al., 2009]. Moreover, the motifs for these variably expressed TFs were enriched in hypomethylated DMRs of aged LEPs overlapping gene promoters, suggesting that these variances may differentially regulate downstream targets. The increase in variance of *KDM2B*, which encodes a DNAm regulatory ZF-CxxC protein, in LEPs is of considerable interest because it is lost in a subset of older women. *KDM2B* binds unmethylated DNA and recruits polycomb repressor complex (PRC)-1 to CpG islands [Farcas et al., 2012], and in the process protects polycomb-bound promoters from *de novo* methylation [Boulard et al., 2015]. Meanwhile, loss of *KDM2B* results in *de novo* methylation of promoters that are typically co-occupied by both *KDM2B* and PRCs [Boulard et al., 2015]. Here, we demonstrated how an increase in molecular noise during aging may lead to sufficient variance in the transcriptomes and methylomes between aged individuals to engender differential susceptibility to development of breast cancers. Because cancer susceptibility indicates a state that could be more easily pushed towards cancer initiation, we can consider the variances between aged individuals to occupy multiple metastable states, some of which represent susceptible phenotypes that can be perturbed and pushed towards cancer.

Our analyses further identified potential ligand-receptor pairs and junctional proteins including tight junctions, desmosomes and gap junction components, that mediate dysregulated cell-cell and cell-microenvironment signaling within the epithelium. We provided experimental validation for the role of gap junction protein, connexin-30 (*GJB6*) in mediating the ability of MEPs to impose an aging phenotype on LEPs [Miyano, Sayaman et al., 2017]. It is unclear whether this occurs chemically through passage of ions or small molecules through gap junction channels, indirectly via gap junction-mediated structural proximity of LEPs and MEPs, or via signaling complexes with connexin-interacting proteins including cytoskeletal elements, tight and adherens junctions, and enzymes like kinases and phosphatases [Dbouk et al., 2009]. How this occurs will require further exploration.

GSEA identified age-dependent enrichment of gene sets in LEPs and MEPs that were commonly dysregulated in breast cancers including gene sets related to: inflammation and immunosenescence, processes synonymous with aging and cancer progression [Fulop et al., 2017]; cell-cycle related targets of E2F transcription factors, which are thought to play a role in regulating cellular senescence [Lanigan et al., 2011]; and targets of the oncogene *MYC.* Whereas our age-specific analyses did not identify oncogenes that were DE between young and old epithelia, gene set enrichment in LEP and MEP revealed another putative example of priming. *Myc* is amplified or overexpressed in ∼35% of breast cancers and exerts pleiotropic effects across the genome [Xu et al., 2010]. We did not detect changes in Myc expression or DNAm in either epithelial lineage with age. However, Myc targets were among the gene sets that were significantly enriched through DE or DV analysis. In the context of cancer progression, Myc is able to induce telomerase activity, which enables bypass of the replicative senescence barrier in mammary epithelial cells [Garbe et al., 2014]. While we do not have evidence for the direct involvement of Myc in this context, we speculate that secondary events such as demethylation at Myc binding sites at target genes could explain the enrichment of Myc relevant signatures. Furthermore, genes that were commonly DE in older women and women at higher-risk of breast cancer relative to young women were enriched for genes involved in chromatin and genome organization. This involves DE of genome-organizer *SATB1*, chromatin-remodeling complex-associated TFs, *BCL11A, TSHZ3* [Kadoch et al., 2013, Faralli et al., 2011]; and pioneer TF *BATF* which can bind condensed chromatin and allow for accessibility of these condensed regions, and is involved in recruitment of chromatin-organizing complexes in a CTCF-dependent manner [Ciofani et al., 2012, Pham et al., 2019]. Involvement of chromatin- and genome-organization explains how transcriptomic changes with age are modulated genome-wide in a lineage-specific manner.

A striking finding of Titus et al. was the ∼2000-fold preponderance of DMGRs in early stage LumA, LumB, and HER2 PAM50 intrinsic subtypes relative to Basal subtypes [Titus et al., 2017a]. In contrast, the number of DMGRs in late-stage cancers were comparable between intrinsic subtypes [Titus et al., 2017a], suggesting that DMGRs in luminal-subtype cancers are acquired early. We propose that this is in large part due to acquisition of age-dependent DM in LEPs. Indeed, CpGs mapping to DMGRs in early stage luminal-subtype cancers represent the majority of age-dependent DM CpGs that we found in LEPs. *That is, genome states that were associated with cancer arise before any detectable cancer exists.* In the context of previous transcriptome profiling that showed ER+ luminal-subtype cancers have similar profiles to those of mature luminal cells [Tharmapalan et al., 2019], we speculate that we have identified an epigenetic mechanism to explain why luminal-subtype cancers have higher incidence in older women. That is, genes for key epigenetic regulatory proteins are themselves epigenetically regulated with age, including hypomethylation of *CXXC5* at a PPR that is enriched for *ESR1* binding motifs. *CXXC5* is a ZF-CxxC family member that binds unmethylated CpGs [Bogdanović et al., 2009, Long et al., 2013], and is known to regulate several signal transduction pathways including TGF-beta, Wnt and ATM/p53 [Xiong et al., 2019]. The importance of *CXXC5* in determination of intrinsic breast cancer subtypes is already well-established as it is one of the 50-gene subtype predictors in PAM50 with prognostic significance [Parker et al., 2009]. Of the TFs predicted to bind *CXXC5* and that are upregulated in luminal-subtype cancers, *ESR1*, *PGR* and *AR* are nuclear receptors that play roles in tumor proliferation, apoptosis, and response to cancer therapy [Conzen et al., 2008]. The role of estradiol-estrogen receptor (E2-ER) in regulating *CXXC5* expression was shown via binding of ER-alpha to estrogen response elements (EREs) upstream of the *CXXC5* initial translation codon [Yaşar et al., 2016]. ESR1 ChIP-seq in normal ER+ luminal epithelia and breast cancer tissues, and in the ER+ breast cancer cell line T-47D showed that *ESR1* binds this *CXXC5* region only in cancers [Chi et al., 2019, ENCODE Project Consortium, 2011, 2012], which could explain why *CXXC5* expression is only modulated in tissues of women with cancer but not in normal primary tissue. Unmethylated *CXXC5* regions in older women may be bound by ZF-CxxC proteins such as *KDM2B* that protect it from methylation; and the variable and subtype-specific increase in *ESR1* during tumorigenesis might lead to competitive binding at this region. We thus propose a mechanism that regulates this binding via the concurrent demethylation of a CpG island in the PPR of *CXXC5* and upregulation of *ESR1* in luminal-subtype cancers – illustrating a case study of a DNAm priming event present in aged epithelia that creates an opportunity for cancer subtype-specific regulation. Indeed, *CXXC5* was shown to participate in E2-driven cellular proliferation through its modulatory effect on the expression of other genes also regulated by E2 [Ayaz et al., 2020]. *CXXC5* overexpression, which we establish to be anti-correlated with DNAm at specific CpGs, is associated with poor prognosis of ER+ breast cancers [Ayaz et al., 2020]. Because *CXXC5* binds unmethylated CpG dinucleotides, upregulation of *CXXC5* in luminal-subtype cancers could lead to increased positive (self-)regulation by TFs by preventing methylation of *CXXC5*-occupied promoter regions. Moreover, *CXXC5* is known to recruit Tet2, a DNA demethylase involved in 5-methylcytosine hydroxylation to 5-hydroxymethylcytosine [Scourzic, et al., 2015, Ma, et al., 2017]. The role of *CXXC5* in Tet2 recruitment in breast is unknown, but it provides a reasonable avenue for further study of feedback mechanisms that can result in widespread epigenetic dysregulation through conservation of unmethylated regions.

Our studies culminate in the exploration of how stereotyped and variant responses are integrated in breast tissue of older women and contribute to differential susceptibility to cancer initiation. That aging leads to more robust DNAm changes than what is found in gene expression raises important questions about the impact of these epigenetic changes. We propose that age-dependent DNAm changes act both through direct canonical regulation of gene expression, and as priming events that require secondary events to occur. Similarly, DNAm changes at *CTCF* binding sites may require additional epigenetic changes to affect chromatin looping. We propose that these age-dependent DNAm priming events mediate susceptibility to age-associated cancers, and the variable occurrence of secondary events across individuals further dictates the risk of individuals to develop cancer. If our priming hypothesis is correct, it simultaneously explains: (i) why hypomethylation of PPRs of MEP-specific genes in aged LEPs leads to robust stereotyped epigenetic changes with age, but leads to varying transcriptomic outcomes that are dependent on TF binding; and (ii) how variable expression of TFs across aged individuals can lead to differential cancer susceptibility.

### Limitations

Our study of luminal and myoepithelial cells is conducted in a finite, pre-stasis HMEC culture system that recapitulates most, but not all, characteristics of normal primary epithelia and may thus miss relevant aspects of normal breast biology. Our data in primary organoids are too limited to perform the rigorous statistical analysis we employed in HMEC, and we thus refer to organoid data to validate observed trends. We further mitigate this limitation by excluding genes with highly discordant dynamic ranges or with loss of directional lineage-specific expression between primary organoids and HMEC. Future expansion of our collection of HMEC samples would allow for increased statistical power for our DE and DV analyses. To address these limitations, we compared our findings to those found in large-scale studies of bulk normal primary breast tissue (n > 100 for expression and n > 200 for DNAm), which validated some of the age-dependent changes we identified in the luminal lineage, suggesting that our HMEC model does in fact capture relevant age-dependent changes, and that LEPs are likely drivers of these age-dependent changes observed in bulk tissue.

## Acknowledgements

We would like to acknowledge the contributions of Dr. James Garbe to the HMEC bank; our current and previous research associates Jennifer Lopez, Jessica Bloom and Jonathan Lee for their technical support; our patient advocates, Susan Samson and Sandy Preto; and the City of Hope Bioinformatics Core led by Dr. Xiwei Wu, specifically Dr. Min-Hsuan Chen who generated the RNA-sequencing raw count data, and Dr. Jinhui Wang who prepared the RNA-sequencing library and performed the sequencing on the Illumina HiSeq2500 platform. The results shown and referenced here are based in part upon data generated by the TCGA Research Network: https://www.cancer.gov/tcga, the ENCODE Consortium: https://www.encodeproject.org/, and the following ENCODE production laboratories: Bernstein Laboratory, Broad Institute, Stamatoyannopoulous Laboratory, University of Washington, and Myers Laboratory, Hudson Alpha Institute for Biotechnology.

The investigators are grateful for support from the National Cancer Institute (NCI) Cancer Metabolism Training Program Postdoctoral Fellowship T32CA221709 (R.W.S). From the National Institutes of Health R01AG040081, U01CA244109, R33AG059206, R01EB024989, R01CA237602; the Department of Defense/Army Breast Cancer Era of Hope Scholar Award BC141351 and Expansion Award BC181737, Conrad N. Hilton Foundation, Yvonne Craig-Aldrich Fund for Cancer Research, and City of Hope Center for Cancer and Aging (M.A.L). Research reported in this publication included work performed in the Integrative Genomics and Bioinformatics, and Analytical Cytometry Cores supported by the National Cancer Institute of the National Institutes of Health under grant number P30CA033572. The content is solely the responsibility of the authors and does not necessarily represent the official views of the National Institutes of Health. The funders had no role in study design, data collection and analysis, decision to publish, or preparation of the manuscript.

## Author Contributions

Conceptualization, R.W.S., M.M., M.A.L.; Methodology, R.W.S., M.M.; Software Programming, R.W.S.; Validation, M.M., R.W.S., P.S.; Formal Analysis, R.W.S.; Investigation, R.W.S., M.M., M.R.S., S.S., P.S., M.E.T., A.Z.; Resources, M.A.L., M.R.S., M.M. D.E.S.; Data Curation, M.R.S., M.M., R.W.S., M.E.T., S.S.; Writing – Original Draft Preparation, R.W.S., M.A.L.; Writing – Review & Editing Preparation, R.W.S., M.A.L., M.M., D.E.S., M.R.S., P.S, S.L.N., A.Z., M.E.T., V.S, S.S.; Visualization, R.W.S.; M.M., P.S.; Supervision, M.A.L., D.E.S.; Project Administration, M.A.L.; Funding Acquisition, M.A.L.

## STAR Methods

### RESOURCE AVAILABILITY

#### Lead Contact

Further information and requests for resources and reagents should be directed to and will be fulfilled by the Lead Contact, Mark A. LaBarge.

#### Materials Availability

Human mammary epithelial cells (HMECs) derived from subjects included in this study are available upon request.

Forward and reverse primer sequences generated in this study are indicated below: GJB6 forward and reverse primers:

5’-CTACAGGCACGAAACCACTCG-3’, 5’ACCCCTCTATCCGAACCTTCT-3’

ELF5 forward and reverse primers:

5’-TAGGGAACAAGGAATTTTTCGGG-3’, 5’-GTACACTAACCTTCGGTCAACC-3’

TBP forward and reverse primers:

5’-GAGCTGTGATGTGAAGTTTCC-3’, 5’-TCTGGGTTTGATCATTCTGTAG-3’

RPS18 forward and reverse primers: 5’-GGGCGGCGGAAAATAG-3’, 5’-CGCCCTCTTGGTGAGGT-3’

Sequences for shGJB6 and shCtrl were ggatacttgctccattcatac and gcttcgcgccgtagtctta, respectively. shCtrl (CSHCTR001LVRU6GP) and shGJB6 Lenti-virus vector (HSH06069132LVRU6GP) were purchased from GeneCopoeia.

#### Data and Code Availability

The datasets generated during this study including RNA-sequencing count data are available at: https://figshare.com/s/769dd9402cd84e19c516 (RNA-seq fastq files will be made available in GEO upon acceptance/publication), and Illumina 450K array beta values are available at: https://figshare.com/s/3dc20a6ea4893ac7f6d3 (Illumina 450K IDAT files will be made available in GEO upon acceptance/publication). The code generated for analyses in this study will be deposited on github (upon acceptance/publication).

### EXPERIMENTAL MODEL AND SUBJECT DETAILS

We examined genome-wide transcription in primary LEPs and MEPs from 43 women across a wide spectrum of ages and cancer risk profiles, and matched DNA methylomes in a subset of these individuals (**Table S1, S2**). We examined two age cohorts: younger <30y women considered to be premenopausal (age range 16-29y, n=32 LEP and MEP samples, m=11 subjects) and older >55y women considered to be postmenopausal (age range 56-72y, n=22 LEP and MEP samples, m=8 subjects) who were considered to have average risk for breast cancer. Breast tissue samples from these subjects were collected from reduction mammoplasties (RM, m=19). We also examined women considered to be at higher risk to develop breast cancer based on accepted genetic or clinical risk factors (age range 25-72y, n=48 LEP and MEP samples, m=24 subjects). These women are either known carriers of high risk germline mutations (*BRCA1* m=8, *BRCA2* m=3, *PALB2* m=1, or *PALB2/APC* m=1), and/or have personal history (m=16) and/or known family history (m=6) of breast cancer (m=11). Breast tissue samples from these subjects were collected from prophylactic mastectomies (PM, m=8) and normal contralateral-to-tumor (CLTT, m=12) or normal peripheral-to-tumor (PTT, m=4) tissue samples.

Prophylactic mastectomy and contralateral to tumor breast tissues were collected at City of Hope. The Institutional Review Boards (IRB) has been approved at City of Hope (Duarte, CA). Breast organoids from reduction mammoplasty, contralateral-to and peripheral-to-tumor breast tissues were prepared at Lawrence Berkeley National Laboratory (Berkeley, CA) with approved IRB for sample distribution and collection from specific locations.

Luminal epithelial (LEPs) and myoepithelial cells (MEPs) FACS-enriched from finite lifespan, non-immortalized human mammary epithelial cells (HMECs) serve as our experimental model system. HMECs are derived from organoids isolated from collected breast tissue samples and grown to 4^th^ passage [LaBarge et al., 2013, Garbe et al., 2009].

### METHOD DETAILS

All computational analyses were conducted using using R (3.5.0) [R Core Team, 2018] (https://www.R-project.org/) and Bioconductor (3.7) [Huber et al., 2015] (https://www.bioconductor.org/) unless otherwise noted.

#### Breast tissue collection and HMEC culture

Primary HMECs were established and maintained according to previously reported protocol using M87A medium containing cholera toxin and oxytocin at 0.5 ng/ml and 0.1nM, respectively [LaBarge et al., 2013, Garbe et al., 2009]. For experiments, 4^th^ passage HMECs were cultured for 4-6 days (depending on strain) to sub-confluence prior to FACS-sorting. Cell cultures were fed every 2 days upto and including the day before FACS-sorting. HMEC strains used in this study were listed in **Table S1** for RNA-seq and **Table S2** for DNAm.

#### Disassociation of uncultured cells from organoids

For dissociation of uncultured cells from organoids, organoids were digested with 0.5% trypsin/EDTA for 10min at 37C with agitation. After trypsin treatment, organoids were disrupted by vigorous shaking for 30sec. Cells were then passed through a 40um cell strainer (BD Falcon).

#### Flow cytometry

LEP and MEP enrichment was performed across multiple studies (M. Miyano, S. Shalabi, M.E. Todhunter). Enrichment was conducted by FACS (BD FACSVantage SE, FACSAriaIII, FACS AriaSORP or Bio-Rad S3 Cell Sorter) or EasySEP Cell Separation (Stem Cell Technologies) using well-established LEP-specific (CD227 or CD133) and MEP-specific (CD271 or CD10) cell-surface markers. Protocols were validated to sort similar populations regardless of antibody combination and enrichment methodology.

Briefly, breast epithelial cells were stained following standard flow cytometry protocol. Primary HMEC strains for RNA-seq were stained with anti-human CD227-FITC (BD Biosciences, clone HMPV) or anti-human CD133-PE (BioLegend, clone7), and anti-human CD271-APC (BioLegend, clone ME20.4) and were separated by FACS or EasySEP. Primary HMEC strains for Infinium 450K array stained with anti-human CD227-FITC (BD Biosciences, clone HMPV) and anti-human CD10-PE (BioLegnend, clone HI10a) and were separated by FACSVantage SE. Uncultured cells dissociated from organoids for RNA-seq were stained with anti-human CD133-PE (BioLegend, clone7) and anti-human CD271-APC (BioLegend, clone ME20.4) and separated by FACS AriaIII.

#### Cell co-cultures

In co-culture study [Miyano, Sayaman et al., 2017], FACS-enriched MEPs from 4^th^ passage HMEC were re-plated on 6-well plates and cultured until the cells were confluent. The cells were treated with mitomycin C (Santa Cruz Biotechnology) at 10μg/ml for 2.5h. In co-culture with shGJB6 study, FACS-enriched shGJB6 transduced MEPs were plated on 6-well plates and cultured until the cells were confluent. FACS-enriched 4^th^ passage LEPs from young <30y women were seeded directly on the mitomycin C-treated or shRNA transduced MEP layer. LEPs from co-cultures were separated by FACS after 10 days for gene expression qPCR analysis. Co-cultured LEPs were stained with anti-human CD133-PE (BioLegend, clone7) and anti-human CD271-APC (BioLegend, clone ME20.4) and were separated using BD FACS ARIAIII.

For Gap junction inhibition assay, cells were cultured with indicated concentration of 18-alpha-Glycryrhetinic acid (Sigma, G8503) for 7days. LEP from co-culture was separated using BD FACSVantage with anti-CD227-FITC (BD Biosciences, 559774, clone HMPV) and anti-CD10-PE (Biolegend, 312204, clone HI10a).

#### RNA isolation and qPCR

Total RNAs were isolated from FACS-enriched LEPs and MEPs with Quick-RNA Microprepkit (Zymo Research). For RNA-seq, isolated RNAs were submitted to Integrative Genomic Core at City of Hope (IGC at COH) for library preparation and sequencing. For qPCR, cDNAs were synthesized with iScirpt reverse transcriptase (BioRad) according to the manufacturer’s manual. Quantitative gene expression analysis was performed by CFX384 real-time PCR (BioRad) with Universal SYBRGreensupermix (BioRad). Data were normalized to RPS18 or TBP by relative standard curve method.

#### RNA-seq sequencing library preparation and sequencing with Illumina Hiseq2500

RNA sequencing libraries were prepared with Kapa RNA mRNA HyperPrep kit (Kapa Biosystems, Cat KR1352) according to the manufacturer’s protocol. Briefly, 100 ng of total RNA from each sample was used for polyA RNA enrichment. The enriched mRNA underwent fragmentation and first strand cDNA synthesis. The combined 2^nd^ cDNA synthesis with dUTP and A-tailing reaction generated the resulting ds cDNA with dAMP to the 3’ ends. The barcoded adaptors were ligated to the ds cDNA fragments. A 10-cycle of PCR was performed to produce the final sequencing library. The libraries were validated with the Agilent Bioanalyzer DNA High Sensitivity Kit and quantified with Qubit. Sequencing was performed on Illumina HiSeq 2500 with the single read mode of 51cycle. Real-time analysis (*RTA v2.2.38*) software was used to process the image analysis.

#### Sequence alignment and gene counts

RNA-Seq reads were trimmed to remove sequencing adapters using *Trimmomatic* [Bolger et al., 2014]. The processed reads were mapped back to the human genome (hg19) using *TOPHAT2* software [Kim et al., 2013]. *HTSeq* [Anders and Huber, 2010] and RSeQC [Wang et al., 2012] software were applied to generate the count matrices and strand information, respectively with default parameters.

#### RNA-seq Pre-processing

RNA-sequencing data pre-processing was conducted in the entirety of the LaBarge sequencing collection across organoids and 4^th^ passage HMECs (n=120 LEP and MEP samples, m=48 individuals) including samples not included in the study. The experimental design group was defined by the combination of the culture condition (organoid, 4^th^ passage), cell type (LEP, MEP) and age/risk status (normal risk RM young <30y, normal risk RM old >55y, and PM/CLTT/PTT without or with germline mutation) of the samples. Raw counts for 34,623 transcripts from RNA-sequencing of FACS-sorted LEPs and MEPs were normalized and regularized log (rlog) transformed in *DESeq2* (*v1.20.0*) [Love et al., 2014] after removal of 30,196 transcripts with zero values across all samples. For initial QA, transformation was run blind to the design matrix (blind=TRUE). Sample QA revealed one outlier sample (subject ID 160, MEP from organoid) that was removed. Rlog transformation was repeated after outlier sample removal without blinding the transformation to the design matrix (∼RNA-seq Batch + Design Group) (blind=FALSE). Rlog were batch-adjusted using *ComBat* (*sva v.3.35.2*) [Johnson et al., 2007, Leek et al., 2020] function with the experimental design group as covariate in the model matrix (∼Design Group). *ComBat* batch-adjusted rlog values used for quality control and assessment of transcripts and filtered as discussed below. Filtered *ComBat* batch-adjusted rlog values were subsetted for samples of interest and used for visualization (*ggplot v.2_3.3.3, gplots v.3.0.3::heatmap.2*) [Wickham, 2016, Warnes et al., 2020] and downstream analysis of expression values. Subject-level data were calculated as the mean value of the batch-adjusted rlog values for subjects with multiple samples.

#### RNA-seq Filtering

During quality control assessment, 5 entries (no feature, ambiguous, too low quality, not aligned, aligned not unique) were filtered out. Count data were imported into *edgeR* (*v.3.22.5*) [Robinson et al., 2010] as a *DGEList* object. Transcripts with low counts were determined using the *edgeR::filterbyExpr* function with experimental design group and batch as covariates in the design matrix (∼0 + Design Group + RNA-seq Batch); 14,647 transcripts with low counts were removed. Next, genes with highly discordant expression levels between organoids and 4^th^ passage HMECs were considered. Mean subject-level batch-adjusted rlog values were calculated for each lineage and age group. Linear regression was performed on the gene expression value means from cells isolated from organoid vs. 4th passage culture in each of the RM LEP <30y, LEP >55y, MEP <30y, MEP >55y subsets, and transcripts with absolute value of model residuals ≥ 6 (∼4*sd) in either of the 4 subsets were considered outliers; 423 transcripts were flagged for exclusion in the DGEList object. Finally, we considered genes that did not maintain consistent lineage-specific expression between organoids and 4th passage. Normalization factors were calculated for the filtered data using *edgeR::calcNormFactors* function using TMM method. DE analysis in *limma voom* (*v.3.36.5*) [Ritchie et al., 2015, Law et al., 2014] was performed between LEPs and MEPs from younger <30 women in both organoids and 4^th^ passage cells (see Quantification and Statistical Analysis). To more fully capture deviations, we used a less stringent cut-off, and lineage-specific genes were defined to be those with DE *adj. p* < 0.1 between LEPs and MEPs in young <30 women in either organoid and 4^th^ passage culture. 2,220 genes that did not maintain consistent lineage-specific expression between organoids and 4^th^ passage cells were subsequently excluded. 17,328 genes were used for all downstream analyses of 4^th^ passage and organoid data.

#### Batch-adjustment of RNA-seq Data

RNA-seq count data were pooled from five experiments and three studies (M. Miyano, S. Shalabi, M.E. Todhunter) conducted at different times. Each experiment was defined to be an RNA-seq batch. Batch effects were found during QC in hierarchical clustering (*hclust*) and principal component analysis (PCA) (*prcomp*) of LEP and MEP samples based on normalized rlog expression values.

For visualization and down-stream analysis of normalized expression data, rlog values were corrected for batch effects using *ComBat* – an empirical Bayes approach for adjusting data for batch effects that is robust to outliers in small sample sizes [Johnson et al., 2007]. The experimental design group was defined by the combination of culture condition (organoid, p4), cell type (LEP, MEP) and age group/risk group (average risk RM young <30y, average risk RM old >55y, or higher risk PM/CLTT/PTT with or without germline mutation) and was included in the model matrix (∼Design Group). *ComBat* batch-adjustment was applied in the *sva* (*v3.35.2*) package [Leek et al., 2020]. *ComBat* batch-adjusted data were then filtered to remove the set of genes described above. Filtered, batch-adjusted data were checked with PCA, and linear regression analysis to confirm removal of PC association with RNA-seq batch. Hierarchical clustering was also used pre- and post-*ComBat* treatment for visualization of batch effects and the clustering of bridge samples.

For DE analysis of RNA-seq count data in *limma voom* (*v3.36.5*) [Ritchie et al., 2015, Law et al., 2014], RNA-seq batch was included in the linear model along with the above design group (∼0 + Design Group + Batch). For DV analysis of RNA-seq count data in *MDSeq* (*v.1.0.5*) [Ran and Daye, 2017], since we were not able to model multiple samples from the same subject, *ComBatSeq* batch-adjustment (*sva_devel*) [Zhang et al., 2020] of the count data was first performed using the above design group in the model matrix. *ComBatSeq* batch-adjusted count data were then normalized using TMM method (*MDSeq::normalize.counts*). Subject-level data were generated as the mean value of *ComBatSeq* batch-adjusted normalized data for subjects with multiple samples and used in the DV analysis.

#### Annotation of RNA-seq Data

RNA-seq transcript Ensembl IDs were mapped to corresponding gene symbols, Entrez IDs and Uniprot IDs using *EnsDb.Hsapiens.v86* (*v2.99.0*) databse [Rainer, 2017].

#### DNA Methylation Pre-processing

Infinium 450K array pre-processing was done in the *ChAMP* (*v.2.10.2*) package [Tian et al., 2017, Morris et al., 2014]. IDAT files were imported (*ChAMP::champ.import*) from LEP and MEP samples across the two age groups (<30y, n=10 samples, m=4 individuals; >55y, n=10, m=5). Of the 485,557 probes included in the array, 65 with probe IDs beginning with ‘rs’ were automatically excluded; 1,355 probes with detection p-values > 0.1 and 6,336 probes with beadcount <3 in at least 5% of samples were filtered out (*ChAMP::champ.filter*). BMIQ normalization was performed on DNAm beta values (*ChAMP::champ.norm*). DNAm m-values were calculated from beta values (*lumi v.2.32.0::beta2m*) [Du et al., 2008]. Non-batch-adjusted subject-level data was calculated as the mean value of the beta or m-values for subjects with multiple samples.

#### Batch-adjustment of DNA methylation data

For visualization and down-stream analysis of DNAm data, normalized data was first corrected for batch effects using *ComBat* as applied in the *sva* (*v.3.35.2*) package [Johnson et al., 2007, Leek et al., 2020]. Infinium 450K batches were defined by the chip IDs. The experimental design group was defined by the combination of the cell type (LEP, MEP) and age group (younger <30Y and older >55y) and was included in the model matrix (∼Design Group). To avoid beta values from exceeding the biologically understood range of 0 to 1, m-values were batch adjusted using *ComBat*; these *ComBat* batch-adjusted m-values were then converted back to beta values (*lumi v.2.32.0::m2beta*) [Du et al., 2008]. Batch-adjusted data were checked with PCA, and linear regression analysis to confirm removal of PC association with chip batch. Hierarchical clustering was also used pre- and post-*ComBat* treatment for visualization of batch effects and the clustering of bridge samples. *ComBat* batch-adjusted subject-level data was calculated as the mean value of the batch-adjusted beta or m-values for subjects with multiple samples. *ComBat* batch-adjusted beta values were used for visualization (*ggplot v.2_3.3.3, gplots v.3.0.3::heatmap.2*) [Wickham, 2016, Warnes et al., 2020].

For DM analysis of DNAm m-values in *limma* (*v3.36.5*) [Ritchie et al., 2015], chip batch was included in the linear model along with the above design group (∼0 + Design Group + Batch). For DMR analysis of DNAm m-values in *DMRcate* (*v.1.16.0*) [Peters et al., 2015], since we were not able to model multiple samples from the same subject, *ComBat* batch-adjusted m-values were used in the analysis.

#### Infinium 450K array probe exclusion

Additionally, 132,922 probes previously reported to be polymorphic or to have cross-reactivity [Chen et al., 2013, Nordlund et al., 2013; Zhou et al., 2017] and 416 probes located in the Y chromosome were further flagged for removal. A total of 347,280 probes were used for all downstream analyses.

#### Annotation of DNA methylation data

Illumina Infinium HumanMethylation450 v1.2 BeadChip probes were annotated using the Illumina Infinium 450K manifest file (HumanMethylation450_15017482_v1-2.csv; https://support.illumina.com/downloads/humanmethylation450_15017482_v1-2_product_files.html). For simplicity, probes annotated with multiple UCSC reference gene names were assigned to the first gene symbol. Probes with multiple assignments to UCSC reference gene groups where assigned with priority to the following gene regions: TSS1500, TSS200, 5’UTR, 1st exon, gene body, and lastly 3’UTR; those with no annotation were assigned to the Intergenic region. CpG island groups were defined by the relation to UCSC GpG islands: N-shelf, N-shore, CpG island, S-shore, S-shelf; those with no annotation were assigned to open sea regions. Enhancer element region and DNaseI hypersensitivity sites were defined based on the Infinium 450K manifest Enhancer and DHS annotation. Additional annotation of CpG islands were derived from Price et al., HIL Classes: high-density CpG island (HC), intermediate-density CpG island (IC) and non-island (LC), ICshore (regions of intermediate-density CpG island shore that border HCs) [Price et al., 2013].

#### Whole-genome bisulfite sequencing data

Pre-processed WGBS data were provided by P. Senapati and D.E. Schones. WGBS data were generated for LEPs from two younge <30Y and two older >55y women for our other manuscript [Senapati, et al., 2020]. WGBS bigwig files were imported in the *UCSC Genome Browser* (*GRCh37/hg19*) and mapped to the *CXXC5* gene region.

#### Normal Primary Breast Tissue Data

Normalized microarray log_2_ expression data, pheno data and feature data from normal primary breast tissues from 114 women (GSE102088) [Song et al., 2017] were downloaded from the Gene Expression Omnibus (GEO) database (https://www.ncbi.nlm.nih.gov/geo/) using the *GEOquery* (*v.2.48.0*) package [Davis and Meltzer, 2007]. The experimental design group was defined by age groups (young <30y, middle aged >30y <55y, and old >55y).

Illumina 450K array IDAT files from primary breast tissue were downloaded from two independent GEO datasets with 121 samples (GSE101961) [Song et al., 2017] and 100 samples (GSE88883) [Johnson et al., 2017] respectively. Pheno data and feature data were imported using *GEOquery* (*v.2.48.0*) [Davis and Meltzer, 2007]. Similarly, the experimental design group was defined by age groups (young <30y, middle aged >30y <55y, and old >55y). As with LEP and MEP data, detection *p*-value filtering and BMIQ normalization were performed in each dataset. Infinium 450K batches were defined by the chip IDs. DNAm beta values were ComBat batch-adjusted (sva v.3.35.2) [Johnson et al., 2007, Leek et al., 2020] using age groups and BMI groups in the model matrix (∼Age Group + BMI Group), with BMI groups were defined as: underweight <18.5, normal ≥18.5 <25, overweight ≥25 <30 and obese ≥30. The experimental design group was defined by age groups (young <30y, middle aged >30y <55y, and old >55y).

#### The Cancer Genome Atlas Data

Normalized and pre-processed TCGA RNA-seq FPKM-UQ expression values and Infinium 450K DNAm beta values, as well as clinical data, from TCGA were downloaded using *TCGAbiolinks* (*v.2.8.4*) [Colaprico et al., 2015] package. RNA-seq expression data were imported using the following parameters: project = “TCGA-BRCA”; data.category = “Transcriptome Profiling”; data.type = “Gene Expression Quantification”; workflow.type = “HTSeq - FPKM-UQ”; legacy = FALSE. TCGA expression Ensembl IDs were mapped to gene symbols using *EnsDb.Hsapiens.v86*. Infinium 450K DNAm beta values were imported using the following parameters: project = “TCGA-BRCA”; data.category = “DNA methylation”; platform = “Illumina Human Methylation 450”; legacy = TRUE. DNAm beta values for specific probes were not available in the TCGA dataset and were assigned NA values. Samples from men and those lacking annotation for PAM50 breast cancer intrinsic subtype were removed. The experimental design group was defined by PAM50 subtypes (LumA, LumB, Her2, Basal, Normal).

#### ESR1 ChIP-Seq Data

Publicly available *ESR1* ChIP-Seq data (GSE99680) from 10 ER+ breast tumor samples (age range: 37-55y) and five ER+ mature luminal epithelia from normal breast (age range: 19-35y) [Chi et al., 2019] were downloaded from the GEO database; however data from one of the five normal tissues were not correctly formatted and was excluded. The available broad peaks from normal tissue and narrow peaks from tumor tissue were imported into the UCSC Genome Browser (GRCh37/hg19) for visualization.

ENCODE TFBS Uniform Peaks [ENCODE Project Consortium, 2011, 2012] of *ESR1* from ChIP-Seq data from the ER+ T-47D cell line after treatment by estradiol, genestein and BPA (Myers Laboratory, Hudson Alpha Institute for Biotechnology, UCSC Accession: wgEncodeEH001577; wgEncodeEH001556; wgEncodeEH002299) were accessed and visualized directly from the UCSC Genome Browser (GRCh37/hg19). *ESR1* binding sites were then mapped to the *CXXC5* 5-CpG DMR of interest.

#### CTCF Peak Binding Sites in Infinium 450K array

ENCODE TFBS Uniform Peaks [ENCODE Project Consortium, 2011, 2012] of *CTCF* from ChIP-Seq data from the HMEC (Bernstein Laboratory, Broad Institute, Stamatoyannopoulous Laboratory, University of Washington, UCSC Accession: wgEncodeEH000075, wgEncodeEH000419) were downloaded. Uniform narrow peaks from the two datasets were imported and transformed into genomic ranges using the *GenomicRanges* (*v.1.32.7*) package [Lawrence et al., 2013]. Correspondingly, Infinium 450K probe chromosomal and base pair position (Infinium 450K manifest CHR and MAPINFO) were also imported and transformed into genomic ranges, and merged by overlap (*GenomicRanges::mergeByOverlaps*) with both Broad peaks (14,630 CTCF CpGs) and UW peaks (22,301 CpGs). CTCF peak binding sites in the Infinium 450K array were defined as the union of these overlapped regions from Broad and UW (36,931 CpGs).

#### Functional Annotation of Genes of Interest

Functional roles of genes highlighted in the manuscript were first explored in The Human Gene Database (*GeneCards v5.0*, https://www.genecards.org/) and subsequent literature search. Transcription factors were identified based on [Lambert et al., 2018], and oncogenes and tumor suppressor genes were identified based on [Davoli et al., 2013].

### QUANTIFICATION AND STATISTICAL ANALYSIS

All quantification and statistical analyses were conducted using using R (3.5.0) [R Core Team, 2018] (https://www.R-project.org/) and Bioconductor (3.7) [Huber et al., 2015] (https://www.bioconductor.org/) unless otherwise noted.

#### Differential Expression Analysis

For lineage-specific and age-dependent DE analysis, RNA-seq *edgeR* (*v.3.22.5*) [Robinson et al., 2010] filtered and normalized *DGEList* object (see *DGEList* object in RNA-seq Pre-Processing) were subsetted for 4^th^ passage LEP and MEP RM samples (<30y, n=32 LEP and MEP samples, m=11 subjects; >55y, n=22, m=8). DE analysis was performed with linear modeling in *limma* (*v.3.36.5*) [Ritchie et al., 2015, Law et al., 2014] using the *voom* implementation. The experimental design group was defined by the combination of cell type and age group. Batch was modeled into the design matrix (∼0 + Design Group + Batch). The four contrast terms included comparisons of MEP vs. LEP lineages in younger <30y and MEP vs. LEP in older >55y groups for lineage-specific DE, and older vs. younger cells in LEP and older vs. younger cells in MEP lineages for age-dependent DE. Sample replicates, as well as the paired nature of MEP/LEP samples were accounted for by calculating the correlation between measurements using the *limma::duplicateCorrelation* function [Smyth et al., 2005] and blocking for subject ID. Because the calculated correlations changed the voom weights slightly, *voom* was re-run for a second time. Correlations were then re-calculated using the new *voom* weights. Linear modeling (*limma::lmFit*) was performed on the *voom* transformed data, with blocking for subject ID. Empirical Bayes moderation (*limma::eBayes*) of computed statistics was then applied.

Number of differentially expressed genes (*limma::topTable*) were reported at different Benjamini-Hochberg (BH) adjusted p-value significance levels (*p* < 0.05, <0.01, <0.001) in each of the contrast terms. Highly lineage-specific DE genes were defined as those with DE adj. *p* < 0.001 and fold-change ≥ 2 in contrasts of LEPs and MEPs of younger women. MEP-specific and LEP-specific genes were indicated by (+) lfc and (-) lfc respectively. Lineage-specific DE (adj. *p* < 0.001 and fold-change ≥ 2) was then annotated as lost, gained or maintained in older women. Age-dependent genes were defined to be those with DE adj. *p* < 0.05 in contrasts between older and younger cells within each lineage; genes upregulated in older cells and younger cells were indicated by (+) lfc and (-) lfc respectively.

The same methodology was applied to LEP and MEP samples from organoids but due to limited sample sizes (<30y LEP m=4, MEP m=3; >55y LEP n=3, MEP n=1) only one contrast term, MEP vs. LEP lineages in younger <30y women, was evaluated. Highly lineage-specific DE genes were defined as those with DE adj. *p* < 0.001 and fold-change ≥ 2 in contrasts of LEPs and MEPs of younger women and were used during QC to compare maintenance of lineage-specific genes between organoids and 4th passage cells in younger women. Age-independent DE analysis of cell type-specificity of gene expression was also modeled (∼0 + Cell Type + Batch), comparing all LEPs and all MEPs in both organoids and 4^th^ passage cells. This allowed for an increase in power in the DE analysis of organoids (LEP m=7, MEP m=4) and for direct comparisons of cell type-specific expression of *DNMT1* and *TET2* between organoids and 4^th^ passage cells in all samples regardless of age.

The same methodology was also applied for susceptibility-associated DE analysis of LEP samples. RNA-seq *edgeR* (*v.3.22.5*) filtered and normalized *DGEList* object were subsetted for 4^th^ passage LEP and MEP samples derived from RM from younger women who were considered average risk (n=32 LEP and MEP samples, m=11 subjects) and samples derived from PM/CLTT/PTT from women with high risk germline mutations and/or personal/family history of breast cancer who were considered higher risk (n=48, m=24). The experimental design group was defined by the combination of cell type and risk group. Batch was similarly modeled into the design matrix (∼0 + Design Group + Batch). The two contrast terms included comparisons of higher risk vs. average risk cells in LEPs and higher risk vs. average risk cells in MEPs. Only the former was reported in the results. Susceptibility-associated genes were defined to be those with DE adj. *p* < 0.05 in contrasts between higher risk cells and young average risk cells within each lineage; genes upregulated in higher risk cells and young average risk cells were indicated by (+) lfc and (-) lfc respectively.

DE analysis was also performed in *limma* (*v.3.36.5*) on a publicly available microarray dataset from normal primary breast tissue, GSE102088 [Song et al., 2017]. Normalized log_2_ expression data were downloaded from the Gene Expression Omnibus (GEO) using *GEOquery* (*v.2.48.0*) [Davis and Meltzer, 2007]. Gene-level data were calculated as the mean expression across all probes mapped to a gene. The experimental design group was defined by age group: younger <30y (m=35), middle-aged >30y <55y (m=68), and older >55y (m=11), in the design matrix (∼0 + Design Group). The three contrast terms included pair-wise comparisons between the three age groups. Linear modeling and empirical Bayes moderation of computed statistics with trend parameter = TRUE were applied. Significant age-dependent DE in normal primary tissue was defined as genes with DE adj. *p* < 0.05, while nominally significance level was defined as genes with DE unadj. *p* < 0.05.

#### Differential Variability Analysis

For DV analysis, RNA-seq count data were batch-adjusted using *ComBatSeq* (*sva_devel*) [Zhang et al., 2020] and bach-adjusted values were normalized using TMM method (*MDSeq v.1.0.5::normalize.counts*); the mean value *ComBatSeq* batch-adjusted normalized expression data was then taken for subjects with multiple samples. DV analysis was performed using *MDSeq* (*v.1.0.5*) [Ran and Daye, 2017] on subject-level *ComBatSeq* batch-adjusted normalized count data. The experimental design group was defined by the combination of cell type and age group in the design matrix (∼Design Group). Outlier removal, which removes outlier samples influential upon the effects of covariates, was performed prior to DV analysis with the following parameters: minimum valid sample size threshold was set to 5 samples and significance level cutoff of outlier selections was set to 0.05. The two contrast terms included comparisons of older vs. younger cells in LEP and older vs. younger cells in MEP lineages for age-dependent DV. *MDSeq* was run with default parameters. Testing results were extracted (*MDSeq::extract.ZIMD*) using an inequality test with the following parameters: lfc threshold for inequality testing was set at 0 and with BH *p*-value adjustment method.

Number of differentially variable genes were defined at different BH-adjusted *p*-value significance levels (*p* < 0.05, <0.01, <0.001) in each of the contrast terms. Age-dependent DV genes were defined to be those with DV adj. *p* < 0.05 in contrasts between older and younger cells within each lineage; genes with increases in variances (higher lfc dispersion) in older cells and younger cells were indicated by (+) lfc and (-) lfc in dispersion respectively.

#### Differential Methylation analysis

For age-dependent DM analysis, non-batch-adjusted, filtered, normalized beta values from 4^th^ passage LEP and MEP samples (<30y, n=10 LEP and MEP samples, m=4 individuals; >55y, n=10, m=5) were first converted to m-values (*lumi v.2.32.0::beta2m*) [Du et al., 2008] as recommended for probe-level DM analysis [Du et al., 2010]. DM analysis was performed using *limma* (*v.3.36.5*) [Ritchie et al., 2015] which applies an empirical Bayes approach proposed to provide more stable inference when number of arrays is small [Smyth, 2004]. The experimental design group was defined by the combination of cell type and age group. Batch was modeled into the design matrix (∼0 + Design Group + Batch). The four contrast terms included comparisons of MEP vs. LEP lineages in younger <30y and MEP vs. LEP in older >55y groups for lineage-specific DM, and older vs. younger cells in LEP and older vs. younger cells in MEP lineages for age-dependent DM. Array weights were also computed (*limma::arrayWeights*) and included in the model. Sample replicates, as well as the paired nature of MEP/LEP samples were accounted for by calculating the correlation between measurements. This was done by blocking for subject ID in the *limma::duplicateCorrelation* function [Smyth et al., 2005] with array weights as additional input. Linear modeling (*limma::lmFit*) was performed with blocking for subject ID and computed array weights. Empirical Bayes moderation (*limma::eBayes*) of computed statistics was applied, with trend parameter = TRUE.

Number of differentially methylated CpG sites (*limma::topTable*) were reported at different BH-adjusted p-value significance levels (*p* < 0.05, <0.01, <0.001) in each of the contrast terms. Highly lineage-specific DM CpG sites were defined as those with DM adj. *p* < 0.001 and fold-change ≥ 2 in contrasts of LEPs and MEPs of younger women. MEP-specific and LEP-specific CpG sites were indicated by (+) lfc and (-) lfc respectively. Lineage-specific DM (adj. *p* < 0.001 and fold-change ≥ 2) was then annotated as lost, gained or maintained in older women. Age-dependent DM CpG sites were defined to be probes with DM adj. *p* < 0.05 in contrasts between older and younger cells within each lineage; CpG sites with higher DNAm in older cells and younger cells were indicated by (+) lfc and (-) lfc respectively.

#### Differential Methylated Region analysis

For DMR analysis, DNAm normalized m-values were batch-adjusted using *ComBat* (*sva v.3.35.2*) [Johnson et al., 2007, Leek et al., 2020] and the mean value was taken for subjects with multiple samples. DMR analysis was performed using *DMRcate* (*v.1.16.0*) [Peters et al., 2015] on subject-level *ComBat* batch adjusted m-values. The experimental design group was defined by the combination of cell type and age group in the design matrix (∼0 + Design Group). The four contrast terms included comparisons of MEP vs. LEP lineages in younger <30y and MEP vs. LEP in older >55y groups for lineage-specific DM, and older vs. younger cells in LEP and older vs. younger cells in MEP lineages for age-dependent DM. To annotate input CpGs (*DMRcate*::*cpg.annotate*), the FDR *p*-value cutoff that defines the significance of each individual CpG site were set at different BH-adjusted *p*-values (*p* < 0.05, <0.01, <0.001) in each of the contrast terms. DMRs were then identified at each DM significance level of input CpGs (*DMRcate*::*dmrcate*). *DMRcate* computes a kernel estimate against a null comparison, and was implemented using the following parameters: Gaussian kernel bandwidth, lambda = 1000 nucleotides, scaling factor bandwidth C=2, minimum number of consecutive CpGs constituting a DMR, min.cpgs=2. The *p*-value cutoff to determine DMRs was automatically determined by the number of significant CpGs returned by *limma* (*v.3.36.5*), and *p*-values were adjusted using the BH method. DMRs were then annotated with overlapping promoters using the hg19 genome (*DMRcate::extractRanges*).

Number of DMRs were reported at different BH-adjusted *p*-values (*p* < 0.05, <0.01, <0.001). Highly lineage-specific DMRs were defined as regions with DMR adj. *p* < 0.001 in contrasts of LEPs and MEPs within each age group; MEP-specific and LEP-specific DMRs were indicated by (+) betafc and (-) betafc respectively. Age-dependent DMRs were defined to be regions with DMR adj. *p* < 0.05 in contrasts between older and younger cells within each lineage; CpG sites with higher DNAm in older cells and younger cells were indicated by (+) betafc and (-) betafc respectively.

#### Non-parametric Kruskal-Wallis and Post-hoc Wilcoxon Test

*Limma*-based DE analysis was not performed on publicly available gene expression datasets from breast cancer tissue from the TCGA cohort. Analysis was limited to genes of interest, and non-parametric Kruskal-Wallis (KW) test (*rstatix v.0.5.0::kruskal_test*) [Kassambara, 2020a], a one-way ANOVA on ranks, was used to determine differences in log_2_(FPKM-UQ+1) values between PAM50 subtypes. Post-hoc pair-wise comparison between groups was performed using non-parametric Wilcoxon test (*rstatix v.0.5.0::wilcox_test*) [Kassambara, 2020a]. Wilcoxon BH-adj. *p*-values were annotated (*ggpubr v.0.2.5*) [Kassambara, 2020b] at different significance levels (*p* < 0.05 (*), < 0.01 (**), < 0.001 (***), < 0.0001 (****).

*Limma*-based DM analysis was also not performed on publicly available DNAm datasets from normal primary breast tissue, GSE101961 and GSE88883 [Song et al., 2017 PMID: 29383109, Johnson et al., 2017 PMID: 28693600], or breast cancer tissue from the TCGA cohort. Analysis was limited to CpG sites mapping to genes of interest, and KW test (*rstatix::kruskal_test*) was used to determine differences in DNAm beta values between groups – e.g. age groups in normal primary tissue or PAM50 subtypes in TCGA. KW *p*-values were corrected for multiple testing (BH) across all CpG sites within a gene. Post-hoc pair-wise comparison between groups was performed using Wilcoxon test (*rstatix::wilcox_test*). Wilcoxon BH-adj. *p*-values were likewise annotated (*ggpubr v.0.2.5*) [Kassambara, 2020b] at different significance levels (*p* < 0.05 (*), < 0.01 (**), < 0.001 (***), < 0.0001 (****)).

#### Lepage Test on Location and Scale

Two-sample Lepage test (*nonpar v.0.1-2*) [Pepler, 2017] is a joint non-parametric test of equality for location (central tendency) and scale (variability). Lepage test was performed on the *ComBat* batch-adjusted normalized rlog expression values of genes encoding for junctional proteins between younger and older cells in each lineage. Significant age-dependent modulation of genes for cell-surface junctional proteins in LEPs and MEPs were defined at *p* < 0.05.

#### Lineage-specific Ligand-Receptor Pair Interactions

LRPs [Ramilowski et al., 2015] gene symbols were checked against *EnsDb.Hsapiens.v86* (*v2.99.0*) database [Rainer, 2017] gene symbols and then mapped to Ensembl IDs. Both ligand and receptor median expressions in LEPs and MEPs from younger and older women were calculated from *ComBat* batch-adjusted normalized rlog values. LRPs were defined to be either LEP-specific or MEP-specific based on their lineage-specific DE (adj. *p* < 0.001 and fold-change ≥ 2) in the younger cohort. LRP interactions were considered to be lost in the older cohort when lineage-specific DE of the ligand and/or its cognate receptor was lost in the older cells (not DE at adj. *p* < 0.001, fold change > 2). Lineage-specific LRP interactions between ligands and cognate receptors in the younger cohort and lineage-specific LRP interactions lost in the older cohort were visualized in an interactome map (*migest v.1.8.1, circlize v.0.4.8*) [Abel, 2013, Gu et al., 2014]. Ligands were connected by chord diagrams from the cell type expressing it (cell type-L-gene symbol) to the cell type expressing its cognate receptor (cell type-R-gene symbol).

#### CTCF Peak Binding Sites Overlapping DM CpG sites

Genomic ranges of *ENCODE* uniform *CTCF* peak binding sites (UCSC Accession: wgEncodeEH001577, wgEncodeEH001556) were mapped to genomic positions of Infinium 450K probes (*GenomicRanges v.1.32.7*) [Lawrence et al., 2013]. Lineage-specific DM CTCF sites represented DM CpGs (adj. *p* < 0.001, fold change > 2) overlapped CTCF peak binding sites and were defined to either have LEP-specific or MEP-specific DNAm in younger and older cells.

#### Gene Set Enrichment Analysis

Fast gene set enrichment analysis (*fgsea v.1.6.0*) [Korotkevich et al., 2019] was used to identify age-dependent enrichment of MSigDB (*v7.2*, http://www.gsea-msigdb.org/gsea/msigdb/index.jsp) hallmark gene sets in LEPs or MEPs. MSigDB gene symbols were mapped to Ensembl IDs using *org.Hs.eg.db* (*v.3.6.0*) [Carlson, 2019] and mapped to the RNA-seq datasets. Fast GSEA was performed on either ordered DE t-statistics (*limma*) or DV test statistic for dispersion (*MDSeq*) using 1000 permutations. Minimal and maximum gene set sizes were set to 15 and 500 respectively. Enriched gene sets were defined as those with enrichment BH adj. *p* < 0.05.

#### Transcription Factor Motif Enrichment

TF motif enrichment analysis was performed on each of the age-dependent DMRs in LEPs (adj. *p* < 0.05) that were either hypomethylated in older or younger women. Enriched motifs within +/-250bp of the given CpG sites in the DMR were identified using *ELMER* (*v.2.4.4*) [Silva et al., 2019] which uses the 771 human binding models from the Homo sapiens Comprehensive Model Collection (*HOCOMOCO v11*) database [Kulakovskiy et al., 2017]. *ELMER* enrichment was run using the *Probes.motif.hg19.450K* dataset which contains motif occurrences within probe regions using the following parameters: minimum incidence of the motif in the DMR probes was set to 1 and FDR *p*-value cut off at 0.05. *ELMER* enriched motifs (FDR *p* < 0.05) were then mapped to the *HOCOMOCO* database to extract TF gene symbols and consensus sequences. For each enriched motif, the number of associated age-dependent DMRs (adj. *p* < 0.05) hypomethylated in younger and older cells were quantified. Enriched TFs and consequence sequences associated with ≥ 50 hypomethylated DMRs in younger and older cells were reported. TFs with age-dependent DE (adj. *p* < 0.05) and DV (adj. *p* < 0.05) and TFs with susceptibility-associated DE (adj. *p* < 0.05) in LEPs were mapped to age-dependent DMRs, and overlapping gene promoters (*DMRcate::extractRanges*) were annotated and visualized (*circlize*).

#### Protein-protein Interactions and Network Analysis

PPI analysis was performed using the *STRING* database (*stringdb v.11*, https://string-db.org/). PPI analysis of the 175 common aging-dependent and susceptibility-associated DE gene set in LEPs showed a PPI enrichment *p* = 4.3×10^-7^ with 156 nodes and 93 edges (expected 53), and average node degree of 1.19 and average local clustering coefficient of 0.333. Network was visualized in *igraph* (v.1.2.5) [Csárdi et al., 2006], and only the largest fully connected main network of 60 genes was plotted. Community detection was performed on this main network using *igraph::cluster_optimal* algorithm [Brandes et al., 2008], which yielded 7 communities. Each community was analyzed in *stringdb* for functional network enrichment. All enriched Gene Ontology (GO) biological processes, cellular compartments, molecular functions, KEGG and Reactome pathways, Pfam/InterPro/SMART protein domains, *stringdb* local network cluster and annotated keywords (FDR *p* < 0.05) were downloaded, and representative terms were reported and plotted.

PPI analysis on the union of 825 DE genes in LEPs from aging-exclusive, susceptibility-exclusive and common aging-dependent and susceptibility-associated gene sets showed a PPI enrichment *p* = 1.4×10^-6^ with 739 nodes and 820 edges (expected 693), and average node degree of 2.22 and average local clustering coefficient of 0.347. Functional network enrichment from *stringdb* (FDR *p* < 0.05) were downloaded, and representative terms reported. Network was visualized in *igraph*, and only the largest fully connected main network of 289 genes was plotted, with the number of interactions between the three gene sets quantified and annotated.

#### Correlation of Gene Expression and DNA methylation in 4th passage LEPs and MEPs

Subject-level expression and DNAm values were calculated for subjects with multiple samples by taking the mean of *ComBat* batch-adjusted values. Spearman correlations of batch-adjusted normalized rlog gene expression values and batch-adjusted DNAm m-values (*stats::cor.test)* [R Core Team, 2018] were calculated for the subset of samples with matched expression and DNam data with correlation coefficients and *p*-values for each gene-CpG pair. Correlation *p*-values were adjusted for multiple testing (BH). Significant correlation of gene expression and DNAm defined at BH-adj. *p* < 0.05.

#### Correlation of Gene Expression and DNA methylation in TCGA

Correlation analysis were limited to samples from women and those with PAM50 intrinsic subtype assignment. Subject-level log_2_(FPKM-UQ+1) expression and DNAm beta values were calculated by taking the mean values for subjects with multiple samples. Gene expression values for *CXXC5* and TFs enriched in the *CXXC5* 5-CpG DMR were extracted; DNAm beta values for the five contiguous CpGs of interest *CXXC5* were also extracted. Pair-wise Pearson correlations were calculated across expression and DNAm values. TFs with positive correlation to *CXXC5* expression (cor coefficient ≥ 0.5) and/or negative correlation to *CXXC5* DNAm of at least one CpG (cor coefficient ≤ −0.5) were selected for further analysis.

#### Linear regression on expression values between organoids and 4th passage cells

Mean subject-level batch-adjusted rlog values were calculated for each lineage and age group in organoids and 4^th^ passage cells. Linear regression was performed on the gene expression value means from cells isolated from organoid vs. 4^th^ passage culture in each of the RM LEP <30y, LEP >55y, MEP <30y, MEP >55y subsets. Regression lines and 95% confidence intervals were plotted along with the y-intercept and slope of the line. Linear regression *R^2^* and *p*-value were reported. Residuals were calculated from the linear model (*stats::lm*) [R Core Team, 2018]. Distribution of residuals were plotted, and mean value and sd of the residuals were calculated. Transcripts with absolute value of model residuals ≥ 6 (∼4*sd) in either of the 4 subsets were considered outliers.

#### Kolmogorov-Smirnov Test on lineage-specific DE or DM log_2_ fold changes

Non-parametric two-sample Kolmogorov-Smirnov (KS) test (*stats::ks.test*) [R Core Team, 2018] on the equality of distributions performed to compare distributions of lineage-specific DE or DM log_2_ fold changes in younger and in older cells. Significance defined at *p* < 0.05.

#### T-test on the differences of DE or DM log_2_ fold changes between age groups

One-sided t-test (*stats::t.test*) [R Core Team, 2018] performed on the distribution of pair-wise differences in lineage-specific DE or DM log_2_ fold changes between age groups (lfc in young - lfc in old) to test if the mean of all values is different from 0. Significance defined at *p* < 0.05, nominal significance defined at *p* ≤ 0.1.

#### T-test on qPCR values between experimental treatments

Two-sided t-test (*stats::t.test*) [R Core Team, 2018] performed on normalized expression of *ELF5* between young LEP/young MEP (Y/Y) co-cultures vs. young LEP/old MEP (Y/O) co-cultures. One-tailed paired t-test performed on normalized expression of *GJB6* in MEPs after treatment with shCntrl or shGJB6 with alternative hypothesis *GJB6* expression is less in the shGJB6 treated cells. One-tailed paired t-test performed on normalized expression of *ELF5* in Y/O co-culture after treatment of MEPs with shCntrl or shGJB6 with alternative hypothesis *ELF5* expression is greater in the shGJB6 treated cells. Significance defined at *p* < 0.05, nominal significance defined at *p* ≤ 0.1.

#### Heatmap Clustering

Heatmap hierarchical clustering of LEP and MEPs samples was performed based on *ComBat* batch-adjusted normalized rlog values for gene expression and *ComBat* batch-adjusted beta values for DNAm. Heatmap clustering (*gplots v.3.0.3::heatmap.2*) [Warnes et al., 2020] were performed using *hclust* Ward’s clustering criterion (*ward.D2*) agglomerative method with Euclidean distances as distance metric. Heatmap clustering (*gplots::heatmap.2*) of TCGA Pearson correlations of gene expression and DNAm data were performed using *hclust* complete-linkage (*complete*) agglomerative method with 1-correlation as distance metric.

#### Hierarchical Clustering with Uncertainty Assessment

*Pvclust* [Suzuki and Shimodaira, 2006] assesses uncertainty in hierarchical clustering analysis and was used in the unsupervised hierarchical clustering of LEP and MEP samples based on the respective *ComBat* batch-adjusted normalized rlog expression values or *ComBat* batch-adjusted DNAm beta values. *Pvclust* was also used in semi-supervised hierarchical clustering of normal primary tissue samples (GSE101961 and GSE88883) [Song et al., 2017, Johnson et al., 2017] based on the *ComBat* batch-adjusted DNAm beta values of the 39 CpG probes identified to have common age-dependent DM in LEPs and in both independent normal primary tissue datasets. Dendrograms were plotted using the *dendextend* (*v.v1.13.4*) [Galili, 2015] package. Approximately unbiased (AU) p-values and bootstrap probability (BP) were annotated. Clusters with AU p-values ≥ 95 were highlighted.

#### Principal Component Analysis

PCA was performed (*stats*::*prcomp*, *factoextra v.1.0.7*) [R Core Team, 2018, Kassambara and Mundt, 2020] on *ComBat* batch-adjusted normalized rlog expression data from RNA-seq, with values scaled and centered. Gene set used for PCA was subsetted to genes with very low and low variance quantile categories in younger cells that had very high and high variance categories in older cells. LEPs and MEPs from younger and older women are shown on a principal PC map of PC1 vs. PC2, with 95% confidence ellipse (*factoextra*) drawn around individuals from each age group.

