## Supplemental Information for "Epigenetic changes with age primes mammary luminal epithelia for cancer initiation"

A

B

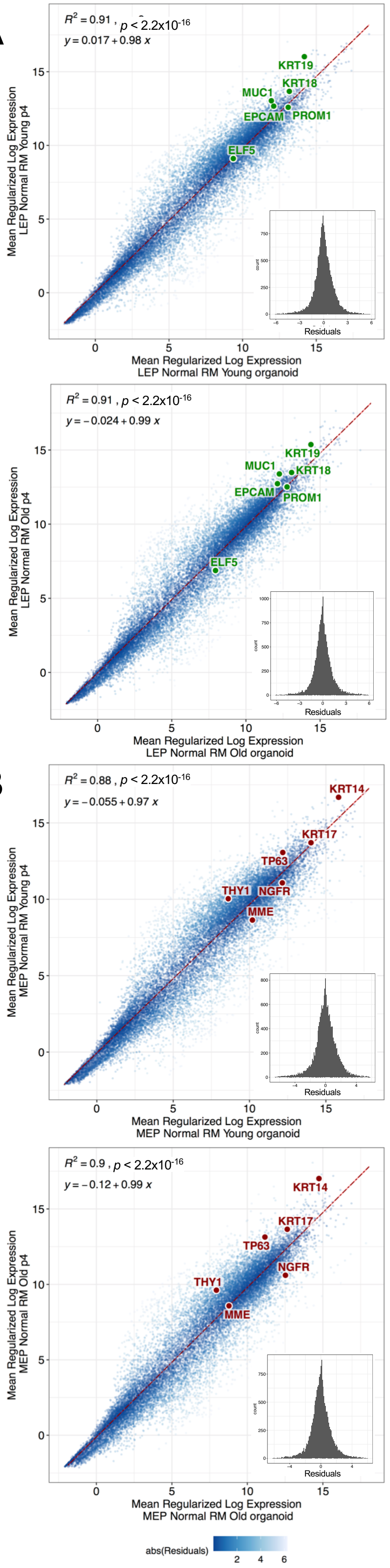


C

**Number of Genes MEP vs LEP in Young ≤30y**

| | # Differentially Expressed | ≥ 2-fold change | ≥ 4-fold change | ≥ 8-fold change |
|---|---|---|---|---|
| adj. p-val < 0.05 | 11,265 | 4,255 | 2,025 | 1,089 |
| adj. p-val < 0.01 | 10,096 | 4,216 | 2,025 | 1,089 |
| adj. p-val < 0.001 | 8,860 | 4,040 | 2,024 | 1,089 |

**Number of Genes MEP vs LEP in Old ≥55y**

| | # Differentially Expressed | ≥ 2-fold change | ≥ 4-fold change | ≥ 8-fold change |
|---|---|---|---|---|
| adj. p-val < 0.05 | 9,620 | 3,857 | 1,827 | 941 |
| adj. p-val < 0.01 | 8,028 | 3,645 | 1,824 | 941 |
| adj. p-val < 0.001 | 6,577 | 3,345 | 1,812 | 940 |

D

E

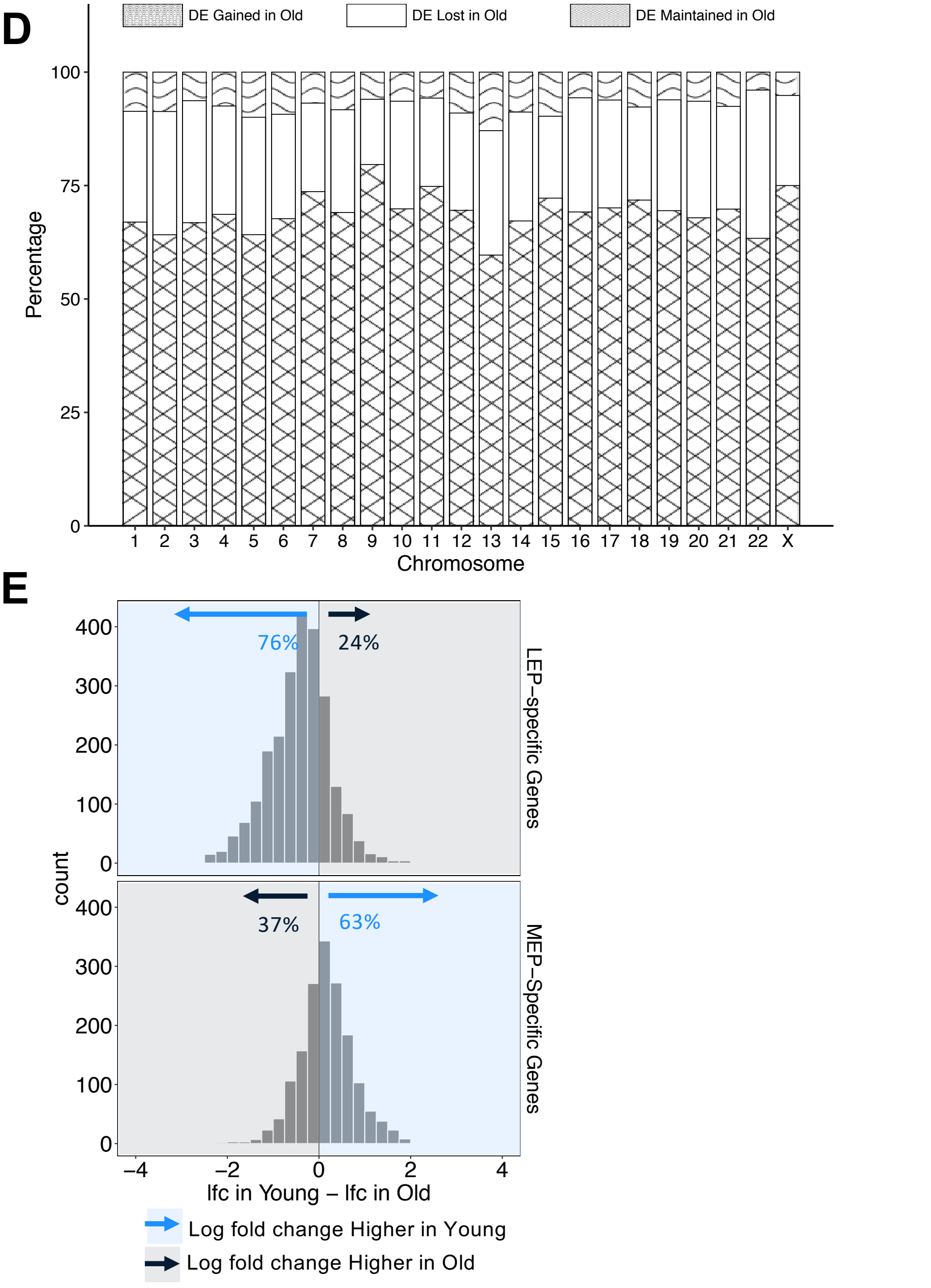

**F**

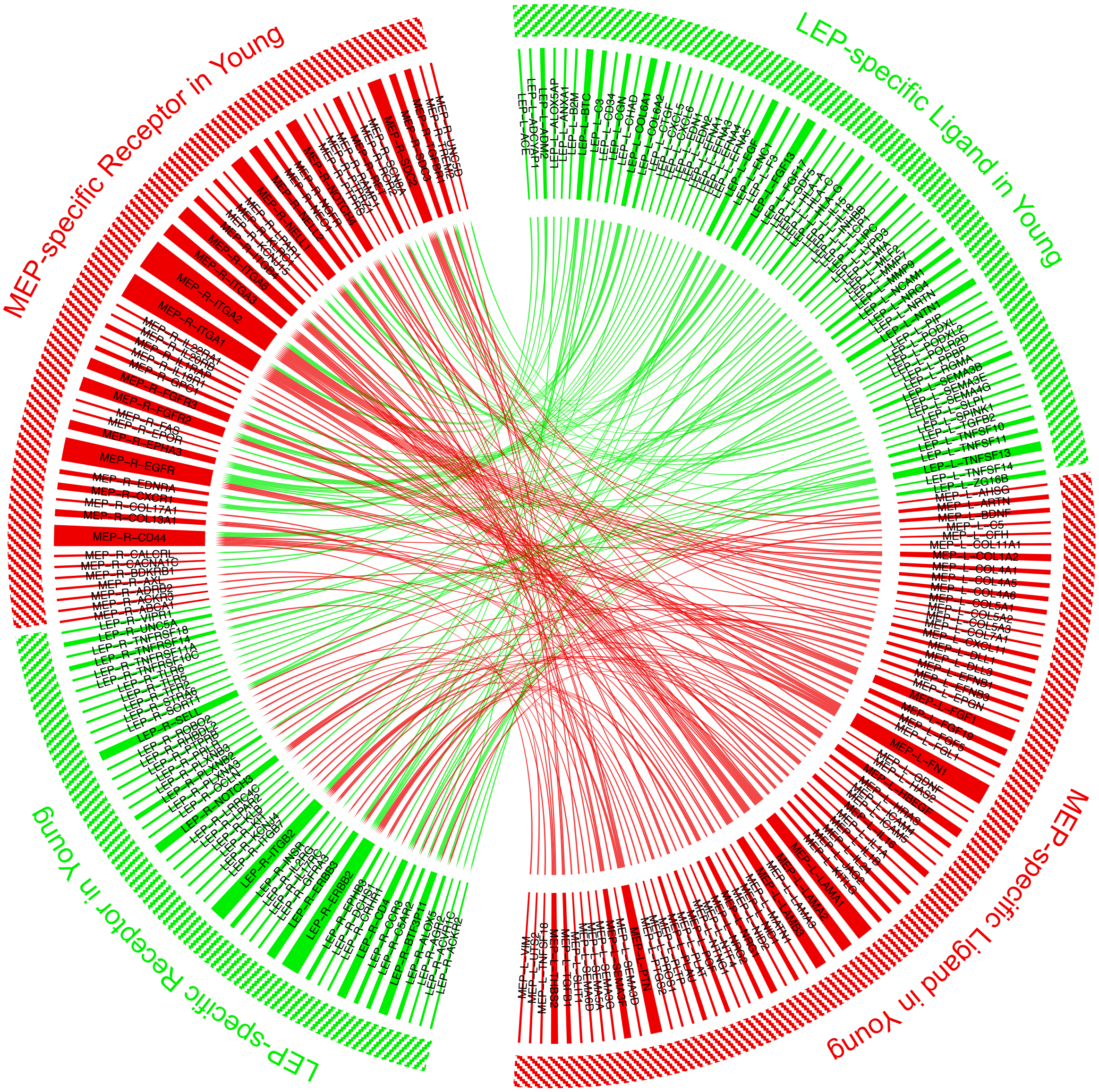


**G**

#### Ligand-associated KEGG Pathways

**LEP**

| KEGG ID | KEGG Pathway | FDR |
|---|---|---|
| hsa04060 | Cytokine-cytokine receptor interaction | 3.16E-10 |
| hsa05323 | Rheumatoid arthritis | 8.08E-07 |
| hsa04360 | Axon guidance | 3.66E-06 |
| hsa04151 | PI3K-Akt signaling pathway | 4.74E-05 |
| hsa05168 | Herpes simplex infection | 0.0006 |
| hsa04010 | MAPK signaling pathway | 0.00081 |
| hsa04015 | Rap1 signaling pathway | 0.00081 |
| hsa04612 | Antigen processing and presentation | 0.00098 |
| hsa05165 | Human papillomavirus infection | 0.00098 |
| hsa04014 | Ras signaling pathway | 0.001 |
| hsa04514 | Cell adhesion molecules (CAMs) | 0.001 |

**MEP**

| KEGG ID | KEGG Pathway | FDR |
|---|---|---|
| hsa04151 | PI3K-Akt signaling pathway | 4.02E-13 |
| hsa05200 | Pathways in cancer | 6.93E-12 |
| hsa05146 | Amoebiasis | 8.61E-11 |
| hsa04512 | ECM-receptor interaction | 6.21E-10 |
| hsa04974 | Protein digestion and absorption | 1.19E-09 |
| hsa04510 | Focal adhesion | 2.18E-09 |
| hsa04360 | Axon guidance | 1.04E-08 |
| hsa05222 | Small cell lung cancer | 2.69E-08 |
| hsa04933 | AGE-RAGE signaling pathway in diabetic complications | 3.81E-08 |
| hsa04010 | MAPK signaling pathway | 7.23E-08 |
| hsa05165 | Human papillomavirus infection | 1.44E-07 |
| hsa05224 | Breast cancer | 9.83E-06 |
| hsa04014 | Ras signaling pathway | 1.31E-05 |
| hsa04926 | Relaxin signaling pathway | 6.39E-05 |
| hsa04610 | Complement and coagulation cascades | 8.15E-05 |
| hsa01522 | Endocrine resistance | 0.00019 |
| hsa04060 | Cytokine-cytokine receptor interaction | 0.00028 |
| hsa05145 | Toxoplasmosis | 0.00032 |
| hsa04015 | Rap1 signaling pathway | 0.00053 |
| hsa05140 | Leishmaniasis | 0.00079 |
| hsa05218 | Melanoma | 0.00083 |
| hsa05226 | Gastric cancer | 0.001 |

#### Receptor-associated KEGG Pathways

**LEP**

| KEGG ID | KEGG Pathway | FDR |
|---|---|---|
| hsa04060 | Cytokine-cytokine receptor interaction | 8.65E-07 |
| hsa04360 | Axon guidance | 7.87E-06 |
| hsa04151 | PI3K-Akt signaling pathway | 3.08E-05 |
| hsa04514 | Cell adhesion molecules (CAMs) | 3.08E-05 |

**MEP**

| KEGG ID | KEGG Pathway | FDR |
|---|---|---|
| hsa05200 | Pathways in cancer | 1.72E-09 |
| hsa04060 | Cytokine-cytokine receptor interaction | 4.01E-08 |
| hsa04151 | PI3K-Akt signaling pathway | 4.01E-08 |
| hsa04810 | Regulation of actin cytoskeleton | 5.51E-08 |
| hsa05412 | Arrhythmogenic right ventricular cardiomyopathy (ARVC) | 7.67E-07 |
| hsa04512 | ECM-receptor interaction | 1.24E-06 |
| hsa05410 | Hypertrophic cardiomyopathy (HCM) | 1.24E-06 |
| hsa05414 | Dilated cardiomyopathy (DCM) | 1.48E-06 |
| hsa04640 | Hematopoietic cell lineage | 1.90E-06 |
| hsa05165 | Human papillomavirus infection | 1.17E-05 |
| hsa04010 | MAPK signaling pathway | 7.73E-05 |
| hsa04510 | Focal adhesion | 8.48E-05 |
| hsa05205 | Proteoglycans in cancer | 8.48E-05 |
| hsa05230 | Central carbon metabolism in cancer | 0.0002 |
| hsa01521 | EGFR tyrosine kinase inhibitor resistance | 0.00037 |
| hsa04020 | Calcium signaling pathway | 0.0006 |
| hsa04015 | Rap1 signaling pathway | 0.001 |

**H**

### Ligand-associated KEGG Pathways

LEP

| KEGG ID | KEGG Pathway | FDR | Genes |
|---|---|---|---|
| hsa05168 | Herpes simplex infection | 1.08E-05 | C3,HLA-A,HLA-C,HLA-G,IL15,TNFSF14 |
| hsa04514 | Cell adhesion molecules (CAMs) | 4.39E-05 | CD34,HLA-A,HLA-C,HLA-G,NCAM1 |
| hsa04612 | Antigen processing and presentation | 5.09E-05 | B2M,HLA-A,HLA-C,HLA-G |
| hsa04060 | Cytokine-cytokine receptor interaction | 0.0003 | GDF5,IL15,TNFSF11,TNFSF13,TNFSF14 |
| hsa04940 | Type I diabetes mellitus | 0.0003 | HLA-A,HLA-C,HLA-G |
| hsa05330 | Allograft rejection | 0.0003 | HLA-A,HLA-C,HLA-G |
| hsa05332 | Graft-versus-host disease | 0.0003 | HLA-A,HLA-C,HLA-G |
| hsa04145 | Phagosome | 0.00038 | C3,HLA-A,HLA-C,HLA-G |
| hsa05320 | Autoimmune thyroid disease | 0.00038 | HLA-A,HLA-C,HLA-G |
| hsa05416 | Viral myocarditis | 0.00045 | HLA-A,HLA-C,HLA-G |
| hsa05167 | Kaposi's sarcoma-associated herpesvirus infection | 0.00067 | C3,HLA-A,HLA-C,HLA-G |
| hsa05203 | Viral carcinogenesis | 0.00067 | C3,HLA-A,HLA-C,HLA-G |
| hsa05323 | Rheumatoid arthritis | 0.0011 | IL15,TNFSF11,TNFSF13 |
| hsa05166 | HTLV-I infection | 0.0017 | HLA-A,HLA-C,HLA-G,IL15 |
| hsa04650 | Natural killer cell mediated cytotoxicity | 0.0029 | HLA-A,HLA-C,HLA-G |
| hsa04218 | Cellular senescence | 0.0052 | HLA-A,HLA-C,HLA-G |
| hsa04672 | Intestinal immune network for IgA production | 0.0066 | IL15,TNFSF13 |
| hsa05169 | Epstein-Barr virus infection | 0.0084 | HLA-A,HLA-C,HLA-G |

MEP

| KEGG ID | KEGG Pathway | FDR | Genes |
|---|---|---|---|
| hsa05224 | Breast cancer | 1.71E-06 | DLL1,DLL3,FGF1,FGF19,HRAS,JAG2 |
| hsa01522 | Endocrine resistance | 4.51E-06 | DLL1,DLL3,HBEGF,HRAS,JAG2 |
| hsa04330 | Notch signaling pathway | 0.00072 | DLL1,DLL3,JAG2 |
| hsa05200 | Pathways in cancer | 0.00072 | DLL1,DLL3,FGF1,FGF19,HRAS,JAG2 |
| hsa04151 | PI3K-Akt signaling pathway | 0.00088 | COL1A2,FGF1,FGF19,HRAS,THBS2 |
| hsa05218 | Melanoma | 0.0014 | FGF1,FGF19,HRAS |
| hsa04658 | Th1 and Th2 cell differentiation | 0.0021 | DLL1,DLL3,JAG2 |
| hsa05226 | Gastric cancer | 0.008 | FGF1,FGF19,HRAS |

### Receptor-associated KEGG Pathways

LEP

| KEGG ID | KEGG Pathway | FDR | Genes |
|---|---|---|---|
| hsa04514 | Cell adhesion molecules (CAMs) | 0.0005 | CD4,ITGB2,LRRC4C,SELL |
| hsa04520 | Adherens junction | 0.0014 | ERBB2,INSR,PTPRB |
| hsa04060 | Cytokine-cytokine receptor interaction | 0.0019 | CCR3,IL2RG,TNFRSF11A,TNFRSF14 |
| hsa04360 | Axon guidance | 0.0092 | LRRC4C,PLXNB2,UNC5A |
| hsa05340 | Primary immunodeficiency | 0.0092 | CD4,IL2RG |

MEP

| KEGG ID | KEGG Pathway | FDR | Genes |
|---|---|---|---|
| hsa04010 | MAPK signaling pathway | 0.0012 | CACNA1C,EGFR,FAS,FGFR2,FGFR3 |
| hsa04270 | Vascular smooth muscle contraction | 0.0012 | CACNA1C,CALCRL,EDNRA,RAMP1 |
| hsa05200 | Pathways in cancer | 0.0012 | EDNRA,EGFR,FAS,FGFR2,FGFR3,NOTCH4 |
| hsa05206 | MicroRNAs in cancer | 0.0012 | CD44,EGFR,FGFR3,NOTCH4 |
| hsa04810 | Regulation of actin cytoskeleton | 0.0013 | EGFR,FGFR2,FGFR3,ITGA1 |
| hsa05205 | Proteoglycans in cancer | 0.0013 | CD44,EGFR,FAS,SDC2 |
| hsa05230 | Central carbon metabolism in cancer | 0.0013 | EGFR,FGFR2,FGFR3 |
| hsa01521 | EGFR tyrosine kinase inhibitor resistance | 0.0014 | EGFR,FGFR2,FGFR3 |
| hsa05165 | Human papillomavirus infection | 0.0053 | EGFR,FAS,ITGA1,NOTCH4 |
| hsa04514 | Cell adhesion molecules (CAMs) | 0.0058 | NEO1,SDC2,SDC3 |
| hsa04151 | PI3K-Akt signaling pathway | 0.0062 | EGFR,FGFR2,FGFR3,ITGA1 |
| hsa05332 | Graft-versus-host disease | 0.0074 | FAS,KLRC1 |
| hsa04020 | Calcium signaling pathway | 0.0087 | CACNA1C,EDNRA,EGFR |
| hsa05219 | Bladder cancer | 0.0087 | EGFR,FGFR3 |

**Figure S1. Genome-wide loss of lineage-specific expression with age. Related to Figure 1.** (**A-B**) Pair-wise comparison of *DESeq2* normalized regularized log (rlog) gene expression means between primary organoids and fourth passage (p4) in FACS-enriched (**A**) luminal epithelial (LEP) or (**B**) myoepithelial (MEP) cells isolated from finite-lifespan human mammary epithelial cells (HMEC) derived from reduction mammoplasties of younger (top) or older (bottom) women. Linear regression line (dotted red) and standard error (red) shown with regression $R^2$, coefficient *p*-value, slope and y-intercept annotated; distribution of residuals shown in the inset. Established lineage-specific markers are shown as reference: LEP-specific markers *KRT19, KRT18, MUC1 (CD227), EPCAM, PROM1,* and *ELF5* (annotated in green), and MEP-specific markers *KRT14, KRT17, MME (CD10), NGFR (CD271), THY1,* and *TP63* (annotated in red). (**C**) Table listing the number of differentially expressed (DE) genes between LEPs and MEPs in young women (top) and in older women (bottom) at different BH adj. *p*-value thresholds (< 0.05, 0.01, 0.001) and fold change cut-offs (2-, 4-, 8-fold change) (*limma*). (**D**) Bar chart showing the fractional distribution of DE lineage-specific genes gained (waves), lost (blank), or maintained (crosshatch) with age (*limma adj. p* < 0.001, fold change ≥ 2) based on their chromosomal position. (**E**) Histogram of pairwise differences in lfc in expression between LEPs and MEPs in younger women vs. older women for all DE LEP-specific (top panel) and MEP-specific (bottom panel) genes identified in young women. The percent of genes with higher lfc in younger women (light blue) or higher lfc in older women (dark blue) are indicated. (**F**) Interactome map of the ligand-receptor pairs (LRPs) [Ramilowski et al., 2015] in young women based on lineage-specific DE of ligands and their cognate receptors in LEPs (green) or MEPs (red) (*adj. p* < 0.001, fold change ≥ 2). LRPs are connected by chord diagrams from the cell type expressing the ligand (cell type-L-gene symbol) to the cell type expressing the cognate receptor (cell type-R-gene symbol). Table shows the number and percent of lineage-specific interactions. (**G**) Network functional enrichment identifies top KEGG pathways (*stringdb* FDR *p* < 0.001) associated with lineage-specific DE of ligands and/or cognate receptors in LEPs and MEPs in young women. (**H**) Network functional enrichment identifies top KEGG pathways (*stringdb* FDR *p* < 0.001) associated with loss of lineage-specific DE of ligands and/or cognate receptors in LEPs and MEPs in older women. Loss of LEP LRPs occurs through loss of lineage-specific DE of genes associated with (1) cell-cell and cell-ECM interactions, including CAMs, AGMs and adherens junctions (AJ) through dysregulation of (i) LEP ligands: CAM *CD34* and neural cell adhesion molecule *NCAM1*; (ii) LEP receptors: CAM/AGM *LRRC4C* and CAM selectin *SELL*; and (iii) cognate ligands that bind LEP receptors: prolactin induced protein *PIP* – which bind CAM *CD4*, matrix metallopeptidase *MMP9* – which bind CAM integrin *ITGB2*, neuregulin *NRG4* – which bind AJ receptor tyrosine kinase *ERBB2*, *AHSG* and *HRAS* – which bind AJ insulin receptor *INSR*, pleiotrophin *PTN* – which bind AJ protein tyrosine phosphatase receptor *PTPRB* and AGM plexin *PLXNB2*, and netrin *NTN1* – which bind AGM *UNC5A*; and (2) cytokine, immune and infection-related pathways through dysregulation of (i) LEP ligands: cytokine interleukin *IL15* – which binds cytokine interleukin receptor *IL2RG*, cytokine tumor necrosis factor superfamily *TNFSF11* – which binds cytokine receptors *BTF3P11* and TNF receptor superfamily *TNFRSF11A*, and antigen processing and presentation-associated MHC-I beta chain molecule *B2M*; (ii) LEP receptors: cytokine C-C motif chemokine receptor *CCR3,* and *TNFRSF14* – which binds cytokines*TNFSF13* and *TNFSF14*; and (iii) cognate receptors to LEP ligands: *ROR2* – which bind cytokine ligand growth differentiation factor *GDF5*, *C5AR2* – which bind LEP-ligand complement *C3*, and *NOTCH4* which bind human leukocyte antigen *HLA-C*. Further, *HLA-A* and *HLA-G* are indirectly associated through loss of lineage-specific expression of other ligands binding the *HLA* cognate receptors. Loss of MEP LRPs occurs through loss of lineage-specific DE of genes associated with (1) pathways in cancer and (2) pathways involved with MAPK, EGFR, NOTCH and PI3K-AKT signaling through dysregulation of (i) MEP receptors: cell surface death receptor *FAS, CD44,* and *NOTCH4*; and (ii) MEP ligands: *HRAS*, and fibroblast growth factors *FGF1* and *FGF19* – which bind the epidermal growth factor receptor *EGFR* and fibroblast growth receptors *FGFR2* and *FGFR3* that are associated with multiple cancer pathways; as well as genes associated with (3) MEP contractility through dysregulation of cognate ligands that bind MEP contractility and/or calcium-signaling associated receptors: adrenomedullin *ADM2* – which bind *CALCRL* and *RAMP1*, endothelin *EDN2* – which bind endothelin receptor *EDNRA*, and *NCAM1* – which bind *CACNA1C*. Further, *EGFR, FGFR2* and *FGFR3*-specific signaling in MEP are indirectly associated through loss of lineage-specific expression of cognate ligands *NRG4,* fibrogen-like *FGL1* and serine peptidase inhibitor *SPINK1* – which bind *EGFR*, *FGF17* – which bind *FGFR2* and *FGFR3,* and *NCAM1* – which bind *FGFR2*.

**A**

Number of CpG Sites MEP vs LEP in Young ≤30y

| | # Differentially Methylated | ≥ 2-fold change | ≥ 4-fold change | ≥ 8-fold change |
|---|---|---|---|---|
| adj. p-val < 0.05 | 77,978 | 46,389 | 17,520 | 7,375 |
| adj. p-val < 0.01 | 56,802 | 43,538 | 17,482 | 7,374 |
| adj. p-val < 0.001 | 39,742 | 36,488 | 17,212 | 7,365 |

Number of CpG Sites MEP vs LEP in Old ≥55y

| | # Differentially Methylated | ≥ 2-fold change | ≥ 4-fold change | ≥ 8-fold change |
|---|---|---|---|---|
| adj. p-val < 0.05 | 35,737 | 23,784 | 8,708 | 3,464 |
| adj. p-val < 0.01 | 24,258 | 20,721 | 8,653 | 3,462 |
| adj. p-val < 0.001 | 16,374 | 15,866 | 8,399 | 3,449 |

**B**

DMRs MEP vs LEP in Young ≤30y

| | # Differentially Methylated Regions |
|---|---|
| fdr < 0.05 | 9,783 |
| fdr < 0.01 | 6,880 |
| fdr < 0.001 | 4,570 |

DMRs MEP vs LEP in Old ≥55y

| | # Differentially Methylated Regions |
|---|---|
| fdr < 0.05 | 5,366 |
| fdr < 0.01 | 3,603 |
| fdr < 0.001 | 2,396 |

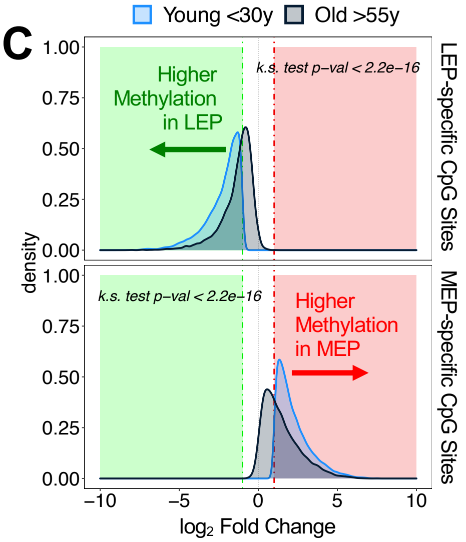


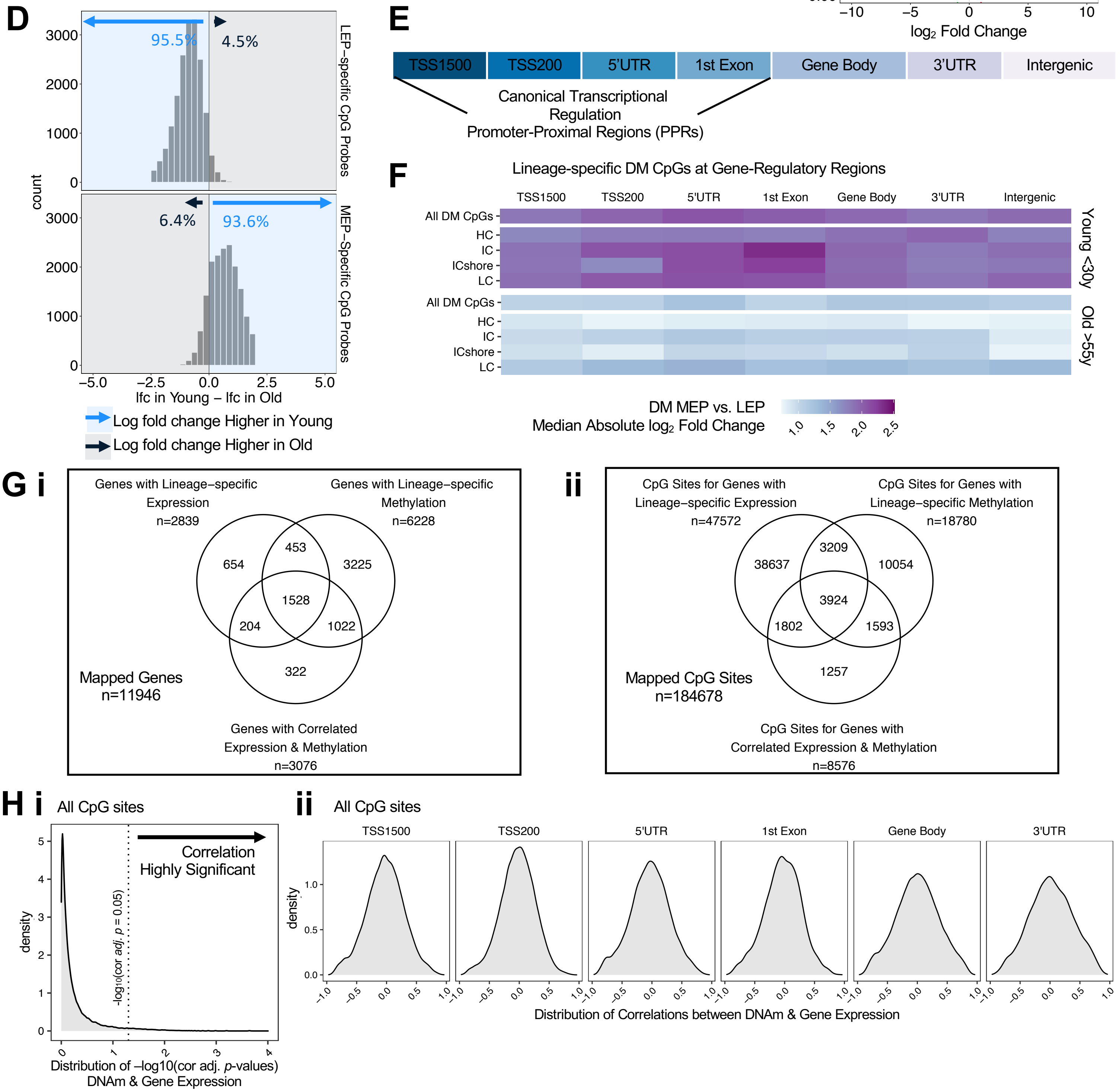

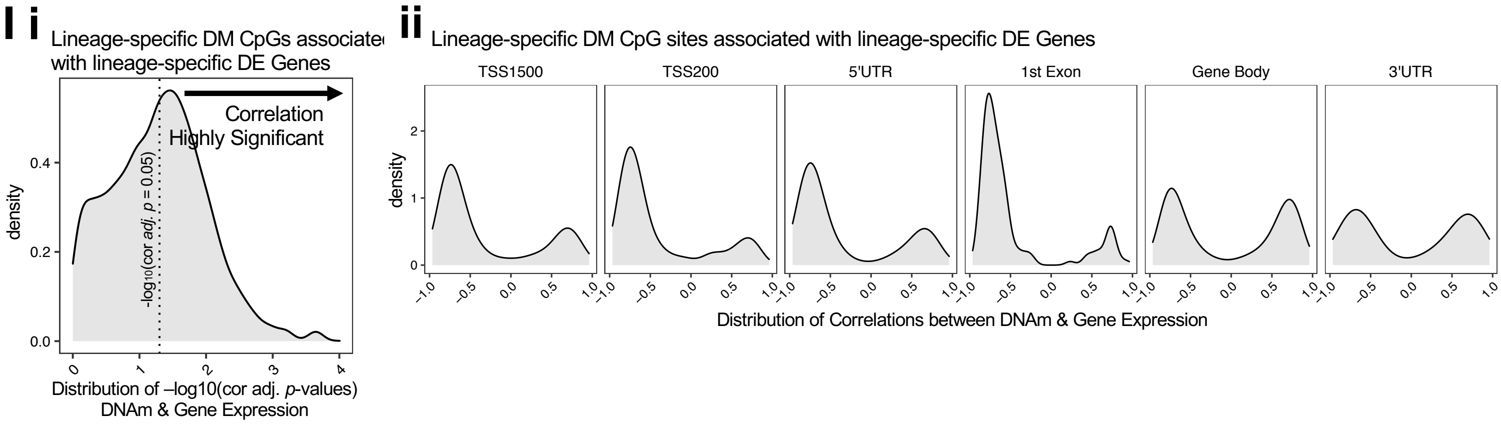


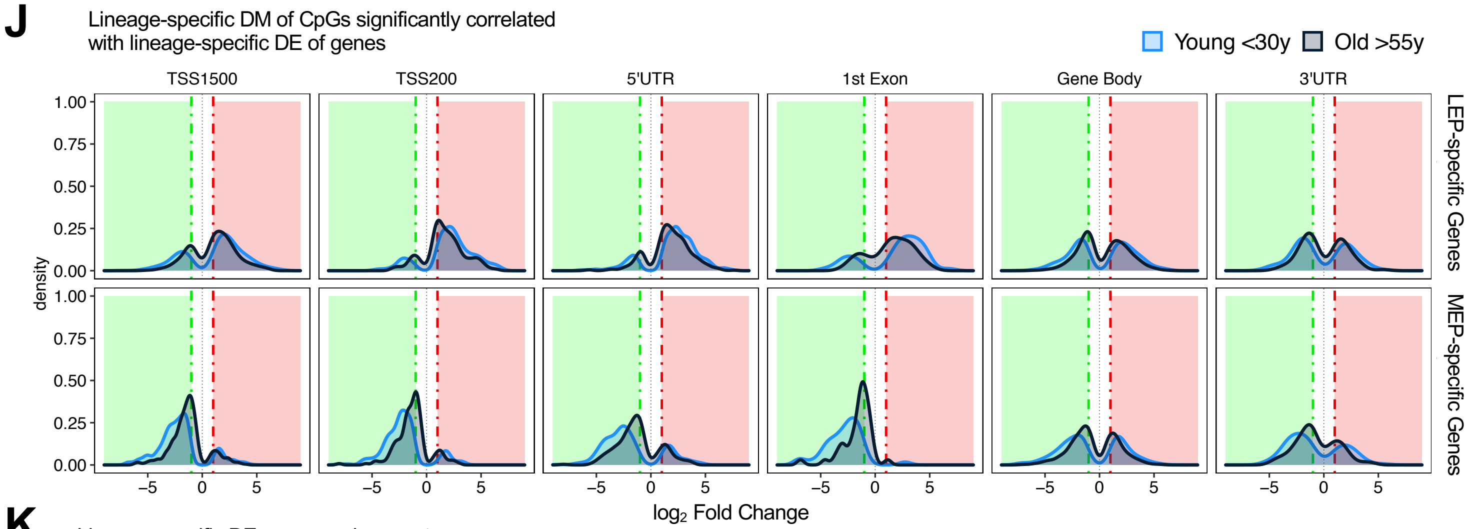


**K**

Lineage-specific DE genes and nearest lineage-specific DM CTCF CpG sites in Old >55y

| Lineage-Specific DE Gene Status | Neareast Lineage-specific DM CTCF CpG Site Status | Count |
|---|---|---|
| DE Lost in Old | DM Lost in Old | 632 |
| DE Lost in Old | DM Maintained in Old | 390 |
| DE Maintained in Old | DM Lost in Old | 1883 |
| DE Maintained in Old | DM Maintained in Old | 1135 |

**Figure S2. Loss of lineage fidelity with age is recapitulated in genome-wide DNA methylation. Related to Figure 2.** (**A**) Table listing the number of differentially methylated (DM) CpG sites between LEPs and MEPs in young women (top) and in older women (bottom) at different BH adj. *p*-values thresholds (< 0.05, 0.01, 0.001) and fold change cut-offs (2-, 4-, 8-fold change) (*limma*). (**B**) Table listing the number of differentially methylated regions (DMRs) between LEPs and MEPs in younger women (top) and in older women (bottom) at different FDR *p*-value thresholds (< 0.05, 0.01, 0.001) (*DMRcate*). (**C**) Distributions of lfc in DNA methylation (DNAm) between LEPs and MEPs in younger (light blue) and older subjects (blue gray) for either DM LEP-specific (top panel) or MEP-specific CpG sites (bottom panel) (*limma adj. p* < 0.001, fold change ≥ 2). LEP-specific CpG sites are shown with (-) lfc values and MEP-specific CpG sites with (+) lfc values relative to each other; KS test *p*-values for equality of distributions of lfc between younger and older women are annotated. (**D**) Histogram of pair-wise differences in lfc in DNAm between LEPs and MEPs in younger women vs. older women for all DM LEP-specific (top panel) and MEP-specific (bottom panel) CpG sites. (**E**) Infinium 450K annotation of CpG gene regions: transcription start site TSS1500, TSS200, 5'UTR, 1$^{st}$ exon, gene body, 3'UTR, and intergenic regions. (**F**) Heatmap of the median value of DM absolute lfc between LEPs and MEPs in young (top panel) and older (bottom panel) subjects across gene regions: TSS1500, TSS200, 5'UTR, 1$^{st}$ exon, gene body, and 3'UTR for all CpG sites and for specific annotated regulatory features denoting CpG islands HIL classes: high-density CpG island (HC), intermediate-density CpG island (IC) and non-island (LC), ICshore (regions of intermediate-density CpG island shore that border HCs) [Price et al., 2013]. (**G**) Venn diagram of (**i**) genes with lineage-specific DE (*adj. p* < 0.001, fold change ≥ 2), genes mapping CpG sites with lineage-specific DM (*adj. p* < 0.001, fold change ≥ 2), and genes with expression correlated to DNAm of at least one CpG site (Spearman cor *adj. p* < 0.05); and (**ii**) CpG sites mapping to genes with lineage-specific DE (*adj. p* < 0.001, fold change ≥ 2), CpG sites with lineage-specific DM (*adj. p* < 0.001, fold change ≥ 2), and CpG sites with DNAm correlated to expression of mapped genes (cor *adj. p* < 0.05). (**H**) Density distribution of (**i**) Spearman correlation *p*-values and (**ii**) Spearman correlation coefficients for all gene-CpG pairs across gene regions. (**I**) Density distribution of (**i**) Spearman correlation *p*-values and (**ii**) Spearman correlation coefficients for lineage-specific DE gene-DM CpG pairs (DE and DM *adj. p* < 0.001, fold change ≥ 2) across gene regions. (**J**) Distribution of lfc in DNAm between LEPs and MEPs in younger (light blue) and older subjects (blue gray) at DM CpG sites that are significantly correlated (cor *adj. p* < 0.05) with expression of either DE LEP-specific (top panel) or MEP-specific genes (bottom panel) across gene regions. CpG sites with higher DNAm in LEPs are shown with (-) lfc values and CpG sites with higher DNAm in MEPs with (+) lfc values relative to each other. (**K**) Summary table of lineage-specific DE genes (*adj. p* < 0.001, fold change ≥ 2) and the nearest lineage-specific DM CpGs (*adj. p* < 0.001, fold change ≥ 2) overlapping CTCF peak binding sites [ENCODE Project Consortium, 2011, 2012] in younger epithelial cells and their DE and DM status in older cells.

**A**

Number of Genes Young ≤30y vs Old ≥55y in LEP

| | # Differentially Expressed | ≥ 2-fold change | ≥ 4-fold change | ≥ 8-fold change |
|---|---|---|---|---|
| adj. p-val < 0.05 | 471 | 172 | 39 | 1 |
| adj. p-val < 0.01 | 166 | 82 | 19 | 1 |
| adj. p-val < 0.001 | 48 | 34 | 8 | 1 |

Number of Genes Young ≤30y vs Old ≥55y in MEP

| | # Differentially Expressed | ≥ 2-fold change | ≥ 4-fold change | ≥ 8-fold change |
|---|---|---|---|---|
| adj. p-val < 0.05 | 29 | 18 | 10 | 2 |
| adj. p-val < 0.01 | 2 | 1 | 0 | 0 |
| adj. p-val < 0.001 | 2 | 1 | 0 | 0 |

**B**

Number of CpG Sites Young ≤30y vs Old ≥55y in LEP

| | # Differentially Methylated | ≥ 2-fold change | ≥ 4-fold change | ≥ 8-fold change |
|---|---|---|---|---|
| adj. p-val < 0.05 | 18,522 | 14,343 | 2,543 | 317 |
| adj. p-val < 0.01 | 9,195 | 8,548 | 2,229 | 307 |
| adj. p-val < 0.001 | 3,178 | 3,161 | 1,490 | 261 |

Number of CpG Sites Young ≤30y vs Old ≥55y in MEP

| | # Differentially Methylated | ≥ 2-fold change | ≥ 4-fold change | ≥ 8-fold change |
|---|---|---|---|---|
| adj. p-val < 0.05 | 20 | 20 | 10 | 3 |
| adj. p-val < 0.01 | 3 | 3 | 3 | 2 |
| adj. p-val < 0.001 | 0 | 0 | 0 | 0 |

**C**

DMRs Young ≤30y vs Old ≥55y in LEP

| | # Differentially Methylated Regions |
|---|---|
| fdr < 0.05 | 3,141 |
| fdr < 0.01 | 1,542 |
| fdr < 0.001 | 436 |

DMRs Young ≤30y vs Old ≥55y in MEP

| | # Differentially Methylated Regions |
|---|---|
| fdr < 0.05 | 1 |
| fdr < 0.01 | 1 |
| fdr < 0.001 | 0 |

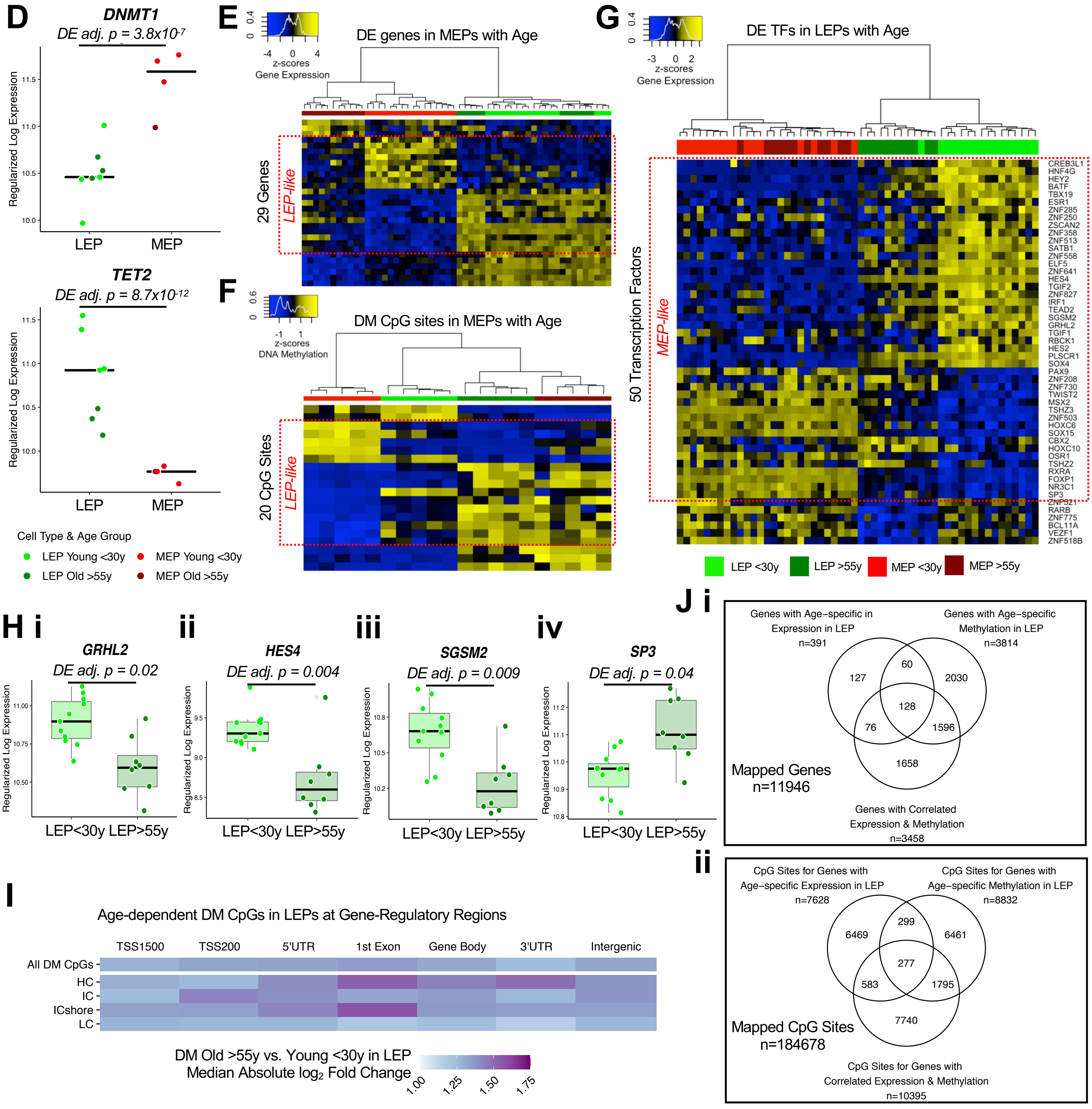

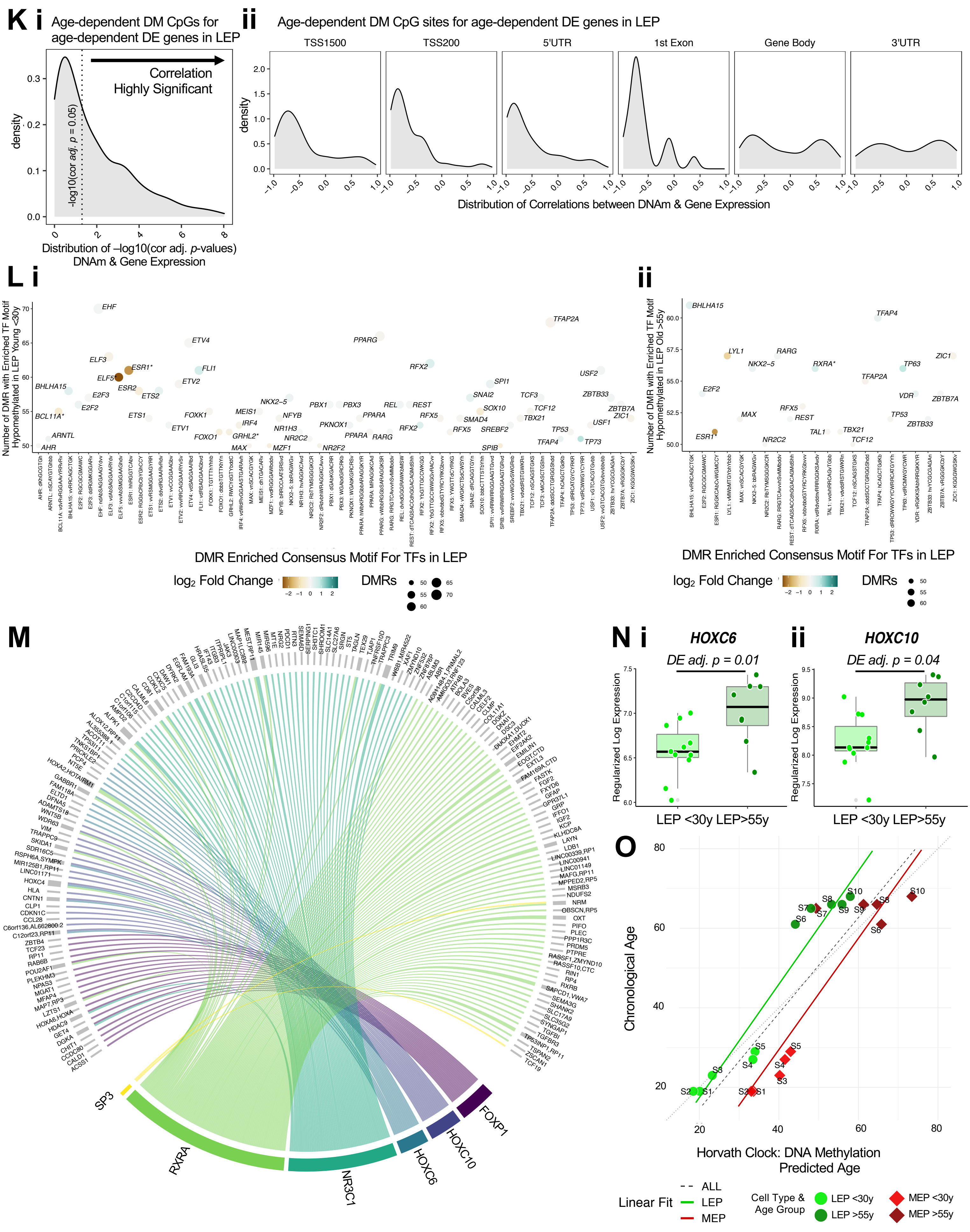
K i Age-dependent DM CpGs for age-dependent DE genes in LEP
Correlation Highly Significant
density
-log10(cor adj. p = 0.05)
Distribution of –log10(cor adj. p-values) DNAm & Gene Expression
ii Age-dependent DM CpG sites for age-dependent DE genes in LEP
TSS1500
TSS200
5'UTR
1st Exon
Gene Body
3'UTR
density
Distribution of Correlations between DNAm & Gene Expression
L i
Number of DMR with Enriched TF Motif Hypomethylated in LEP Young <30y
DMR Enriched Consensus Motif For TFs in LEP
log2 Fold Change
DMRs
ii
Number of DMR with Enriched TF Motif Hypomethylated in LEP Old >55y
DMR Enriched Consensus Motif For TFs in LEP
log2 Fold Change
DMRs
M
SP3
RXRA
NR3C1
HOXC6
HOXC10
FOXP1
N i HOXC6
DE adj. p = 0.01
Regularized Log Expression
LEP <30y LEP>55y
ii HOXC10
DE adj. p = 0.04
Regularized Log Expression
LEP <30y LEP>55y
O
Chronological Age
Horvath Clock: DNA Methylation Predicted Age
Linear Fit
ALL
LEP
MEP
Cell Type & Age Group
LEP <30y
LEP >55y
MEP <30y
MEP >55y

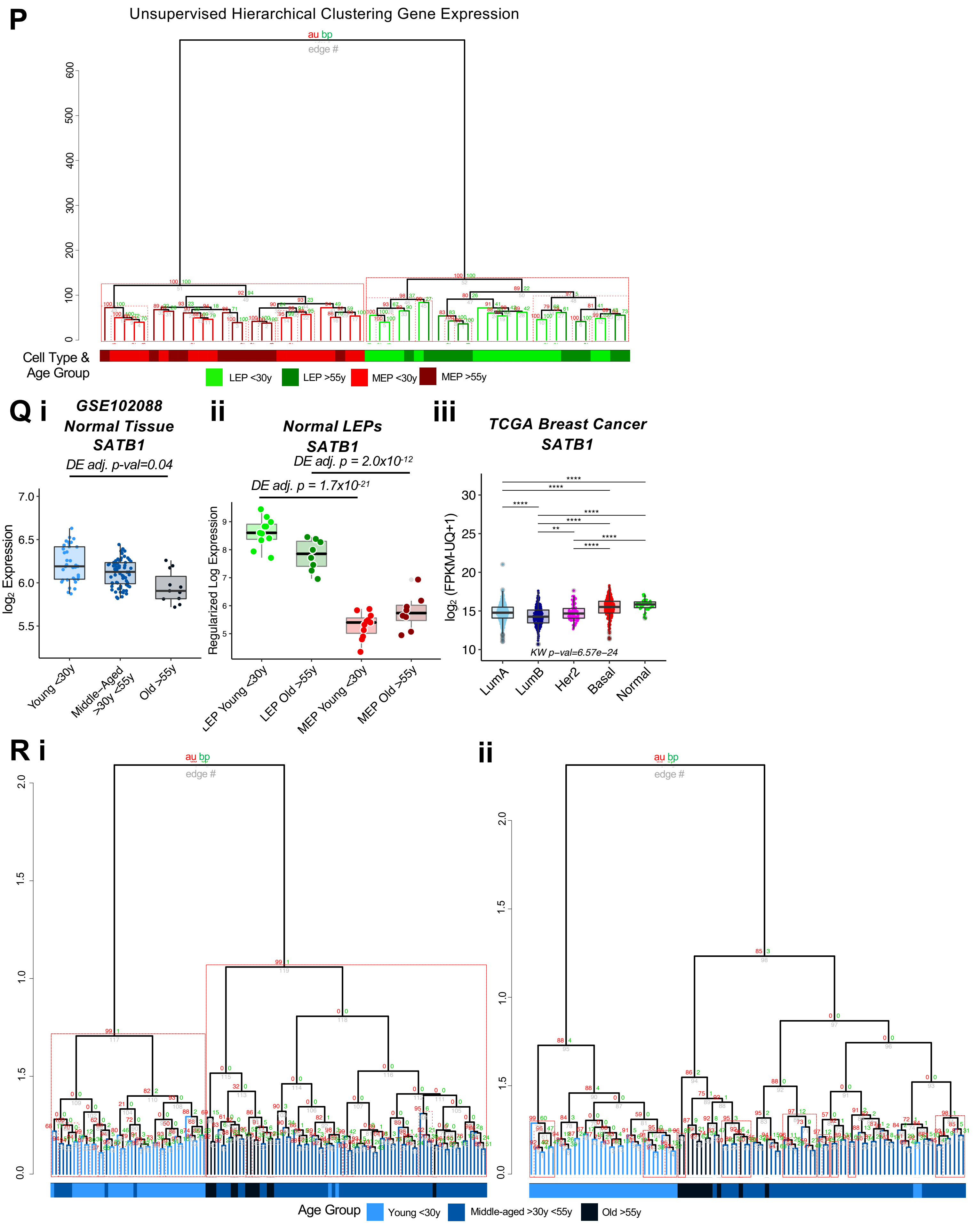
P
Unsupervised Hierarchical Clustering Gene Expression
au bp
edge #
Cell Type &
Age Group
LEP <30y
LEP >55y
MEP <30y
MEP >55y
Q i
GSE102088
Normal Tissue
SATB1
DE adj. p-val=0.04
log2 Expression
Young <30y
Middle-Aged >30y <55y
Old >55y
ii
Normal LEPs
SATB1
DE adj. p = 2.0x10-12
DE adj. p = 1.7x10-21
Regularized Log Expression
LEP Young <30y
LEP Old >55y
MEP Young <30y
MEP Old >55y
iii
TCGA Breast Cancer
SATB1
log2 (FPKM-UQ+1)
KW p-val=6.57e-24
LumA
LumB
Her2
Basal
Normal
R i
au bp
edge #
ii
au bp
edge #
Age Group
Young <30y
Middle-aged >30y <55y
Old >55y

**Figure S3. The luminal lineage is a hotspot for age-dependent directional changes. Related to Figure 3.** (**A-B**) Table listing the (**A**) number of DE genes and (**B**) number of DM CpG sites between younger and older LEPs (top) or MEPs (bottom) at different BH adj. *p*-values thresholds (< 0.05, 0.01, 0.001) and fold change cut-offs (2-, 4-, 8-fold change) (*limma*). (**C**) Table listing the number of DMRs between younger and older LEPs (top) or MEPs (bottom) at different FDR *p*-value thresholds (< 0.05, 0.01, 0.001) (*DMRcate*). (**D**) Dotplots of gene expression rlog values of (**i**) *DNMT1* and (**ii**) *TET2* in LEPs and MEPs from primary organoids. Median values indicated by horizontal bar. MEP vs. LEP DE adj. *p*-values (*limma*) annotated. (**E-G**) Hierarchical clustering of all samples based on (**E**) age-dependent DE genes and (**F**) age-dependent DM CpG sites in MEPs (DE and DM *adj. p* < 0.05), and (**G**) age-dependent DE transcription factors (TFs) in LEPs (*adj. p* < 0.05). Scaled rlog values of DE genes and scaled beta values of DM CpG sites are shown as a heatmap. Clustering performed using Euclidean distances and Ward agglomerative method. (**H**) Boxplots of gene expression rlog values of (**i**) *GRHL2*, (**ii**) *HES4*, (**iii**) *SGSM2* and (**iv**) *SP3* in younger and older LEPs. Age-dependent DE adj. *p*-values (*limma*) are indicated. (**I**) Heatmap of the median value of DM absolute lfc between younger and older LEPs across gene regions: TSS1500, TSS200, 5'UTR, 1$^{st}$ exon, gene body, and 3'UTR for all CpG sites and for specific annotated regulatory features denoting CpG islands HIL classes: high-density CpG island (HC), intermediate-density CpG island (IC) and non-island (LC), ICshore (regions of intermediate-density CpG island shore that border HCs) [Price et al., 2013]. (**J**) Venn diagram of (**i**) genes with age-dependent DE in LEPs (*adj. p* < 0.05), genes mapping CpG sites with age-dependent DM in LEPs (DM *adj. p* < 0.05), and genes with expression correlated to DNAm of at least one CpG site (Spearman cor *adj. p* < 0.05); and (**ii**) CpG sites mapping to genes with age-dependent DE in LEPs (*adj. p* < 0.05), CpG sites with age-dependent DM in LEPs (*adj. p* < 0.05), and CpG sites with DNAm correlated to expression of mapped genes (cor *adj. p* < 0.05). (**K**) Density distribution of (**i**) Spearman correlations *p*-values and (**ii**) Spearman correlation coefficients for age-dependent DE gene-DM CpG pairs in LEPs (DE and DM adj. *p* < 0.05) across gene regions. (**L**) Top TFs with enriched motifs (*ELMER* FDR *p* < 0.05) in ≥ 50 DMRs in LEPs that are differentially hypomethylated in (i) younger or (ii) older women (DMRcate FDR *p* < 0.05). Point sizes reflect the number of DMRs with motif enrichment and are color coded based on lfc of the corresponding TFs. TFs DE with age are annotated with an (*). (**M**) Differentially upregulated TFs (*limma adj. p* < 0.05) in aged LEPs and their corresponding enrichment (*ELMER* FDR *p* < 0.05) in gene promoter regions overlapping DMRs hypomethylated (*DMRcate* FDR *p* < 0.05) in LEPs of older women. (**N**) Boxplots of gene expression rlog values of (**i**) *HOXC6* and (**ii**) *HOXC10* in LEPs of younger and older LEPs women. Age-dependent DE adj. *p*-values are indicated (*limma*). (**O**) Predicted age of LEPs and MEPs based on DNA methylation levels of 353 CpG probes defined in the Horvath epigenetic clock [Horvath, 2013], plotted against chronological age. Regression line for LEPs (green) and MEPs (red) are shown. (**P**) Unsupervised hierarchical clustering of LEPs and MEPs from younger and older women based on rlog expression values of all gene transcripts. Clusters with AU ≥ 95 (*pvclust*) are highlighted (dashed red, with the largest supported cluster in solid red). Clustering performed using Euclidean distances and Ward agglomerative method. (**Q**) Boxplots of *SATB1* gene expression: (**i**) $\log_2$ values in normal primary breast tissue from an independent data set, GSE102088 [Song et al., 2017]; (**ii**) rlog values in LEPs and MEPs of young and older women; and (**iii**) $\log_2$ FPKM values in the TCGA breast cancer cohort. Age-dependent DE adj. *p-values* in normal primary tissue and lineage-specific DE adj. *p*-values in LEPs (*limma*) are indicated. Kruskal-Wallis (KW) test *p*-value between cancer subtypes in TCGA and post-hoc pair-wise Wilcoxon test BH-adj. *p*-value significance levels are annotated (* < 0.05, ** < 0.01, *** < 0.001, **** < 0.0001). (**R**) Semi-supervised hierarchical clustering of normal primary tissue samples from younger <30y, middle-aged >30y <55y and older >55y women from two independent datasets, (i) GSE101961 and (ii) GSE88883 [Song et al., 2017, Johnson et al., 2017] based on the DNAm beta values of the 39 CpGs identified to be DM in LEPs and primary tissue with age. Clusters with AU ≥ 95 (*pvclust*) are highlighted (dashed red, with the largest supported cluster in solid red).

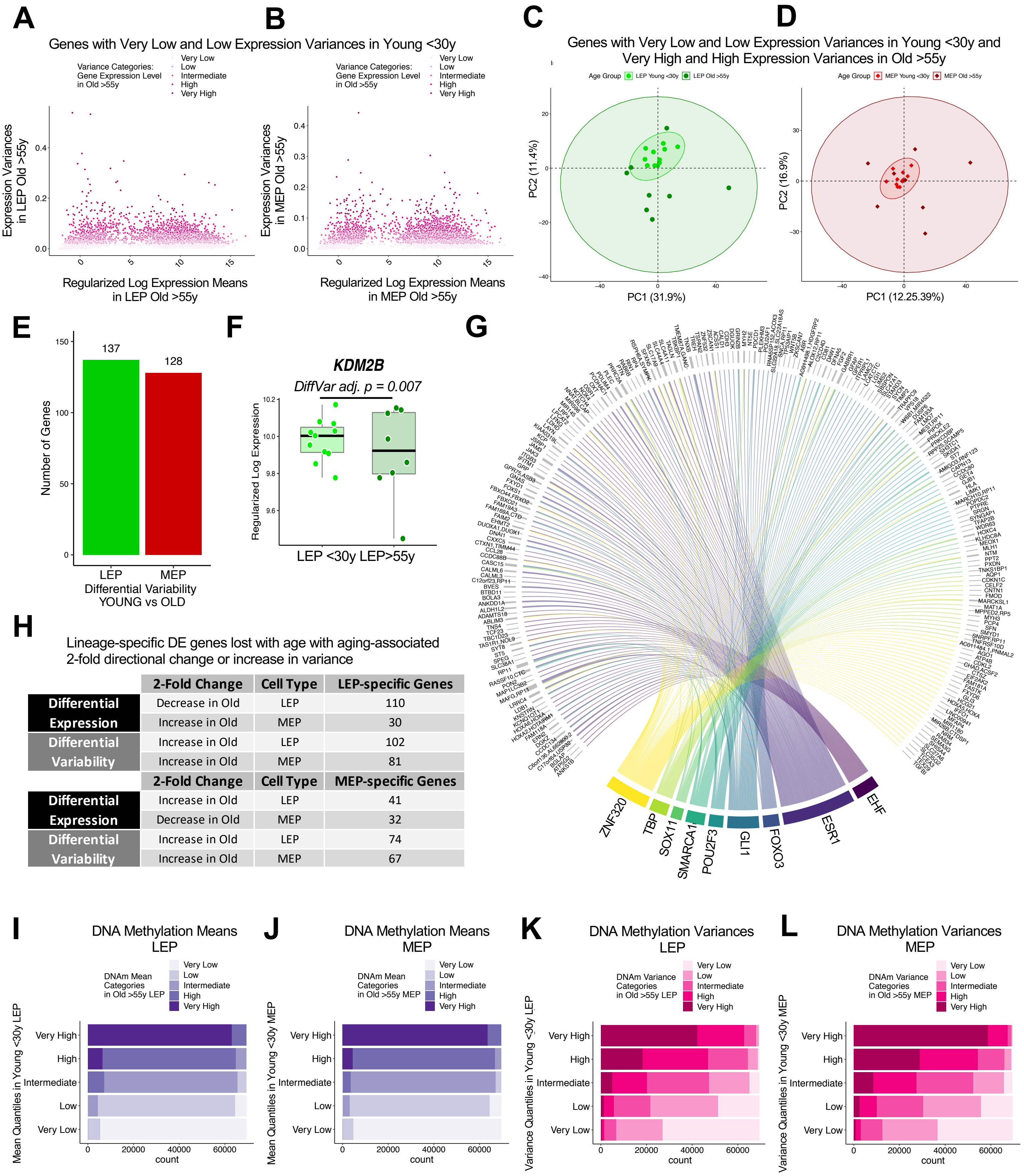


| | 2-Fold Change | Cell Type | LEP-specific Genes |
|---|---|---|---|
| Differential Expression | Decrease in Old | LEP | 110 |
| | Increase in Old | MEP | 30 |
| Differential Variability | Increase in Old | LEP | 102 |
| | Increase in Old | MEP | 81 |
| | **2-Fold Change** | **Cell Type** | **MEP-specific Genes** |
| Differential Expression | Increase in Old | LEP | 41 |
| | Decrease in Old | MEP | 32 |
| Differential Variability | Increase in Old | LEP | 74 |
| | Increase in Old | MEP | 67 |

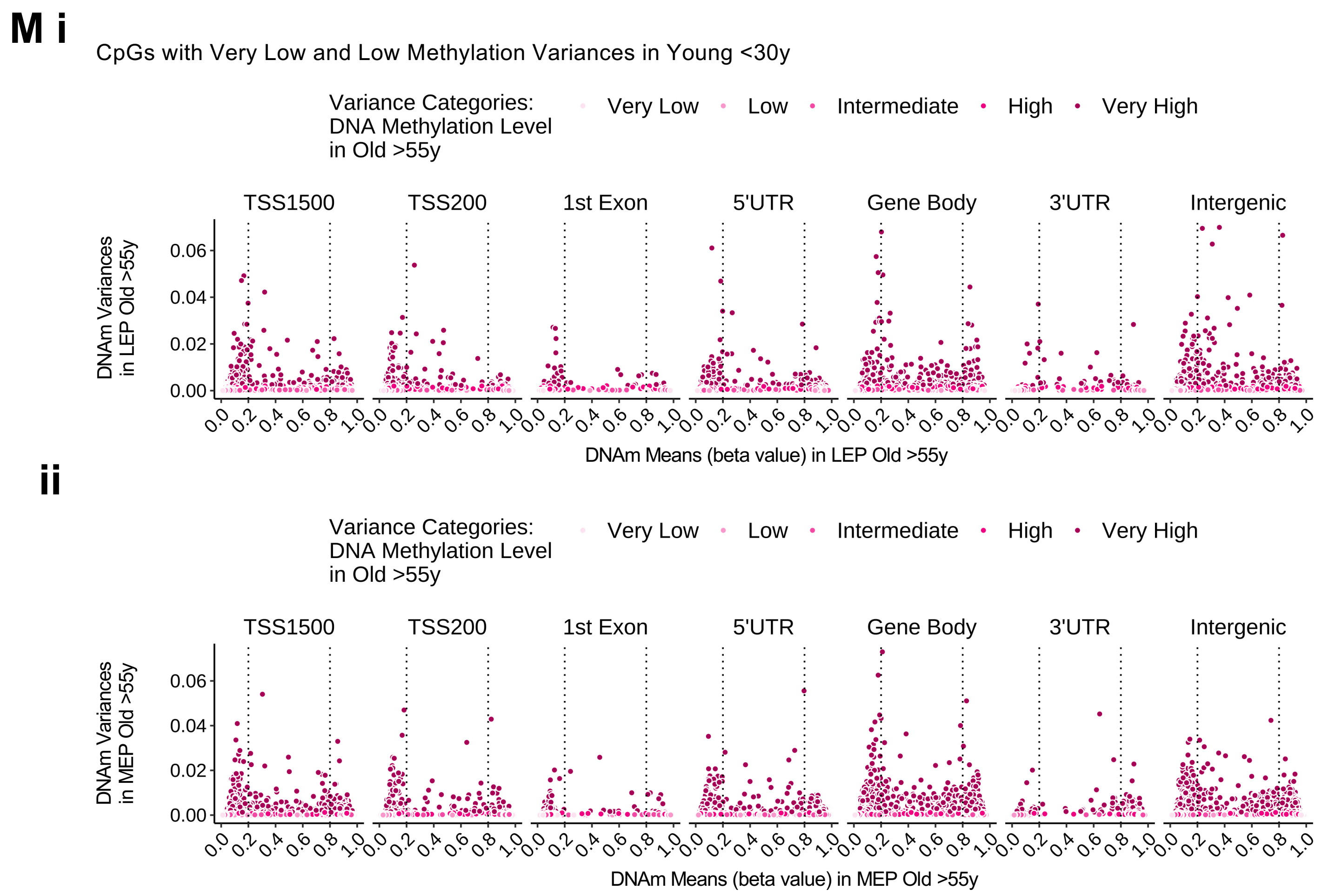
M i
CpGs with Very Low and Low Methylation Variances in Young <30y
Variance Categories: DNA Methylation Level in Old >55y
Very Low
Low
Intermediate
High
Very High
TSS1500
TSS200
1st Exon
5'UTR
Gene Body
3'UTR
Intergenic
DNAm Variances in LEP Old >55y
0.00
0.02
0.04
0.06
0.0
0.2
0.4
0.6
0.8
1.0
DNAm Means (beta value) in LEP Old >55y
ii
DNAm Variances in MEP Old >55y
DNAm Means (beta value) in MEP Old >55y

**Figure S4. Aging-associated increase in variance contributes to loss of lineage fidelity. Related to Figure 4.** (**A, B**) Scatterplot of gene expression means vs. variances of (**A**) LEPs and (**B**) MEPs from older women for the subset of very low and low variance genes identified in younger cells. Genes are color coded based on their variance categories in older cells: very low, low, intermediate, high, very high variance. (**C-D**) Principal Component Analysis (PCA) based on the subset of genes with very low to low variances identified in younger cells that have very high to high variances in older cells. (**C**) LEPs and (**D**) MEPs from younger and older women are shown on a principal PC map of PC1 vs. PC2, with 95% confidence ellipse drawn around individuals from each age group. (**E**) Number of age-dependent differentially variable (DV) genes between younger and older LEPs or MEPs (*MDSeq adj.* $p < 0.05$). (**F**) Boxplot of gene expression rlog values of *KDM2B* in LEPs of younger and older LEPs. Age-dependent DV adj. *p*-values (*MDSeq*) are indicated. (**G**) DV expressed TFs (*adj.* $p < 0.05$) in LEPs and their corresponding enrichment (*ELMER* FDR $p < 0.05$) in gene promoter regions overlapping DMRs hypomethylated (*DMRcate* FDR $p < 0.05$) in LEPs of older women. (**H**) Table showing the number of lineage-specific genes lost with age (DE *adj.* $p < 0.001$, fold change $\geq 2$) that have at least an age-dependent 2-fold directional change (*limma*) or aging-associated 2-fold increase in variance (*MDSeq*) in the older cohort. (**I-L**) DNAm (**I, J**) means and (**K, L**) variances of LEPs and MEPs from younger women are categorized into quantile levels: very low, low, intermediate, high, very high. Corresponding categories of DNAm means and variances in older cells, as defined by threshold values in younger cells, are fractionally represented in color. (**M**) Scatterplot of DNAm means vs. variances of (**i**) LEPs and (**ii**) MEPs from older women across gene regions for the subset of very low and low variance CpG sites identified in younger cells. CpG sites are color coded based on their variance categories in older cells: very low, low, intermediate, high, very high variance. Unmethylated (beta values < 0.2) and methylated (beta values > 0.8) regions are indicated by dashed lines.

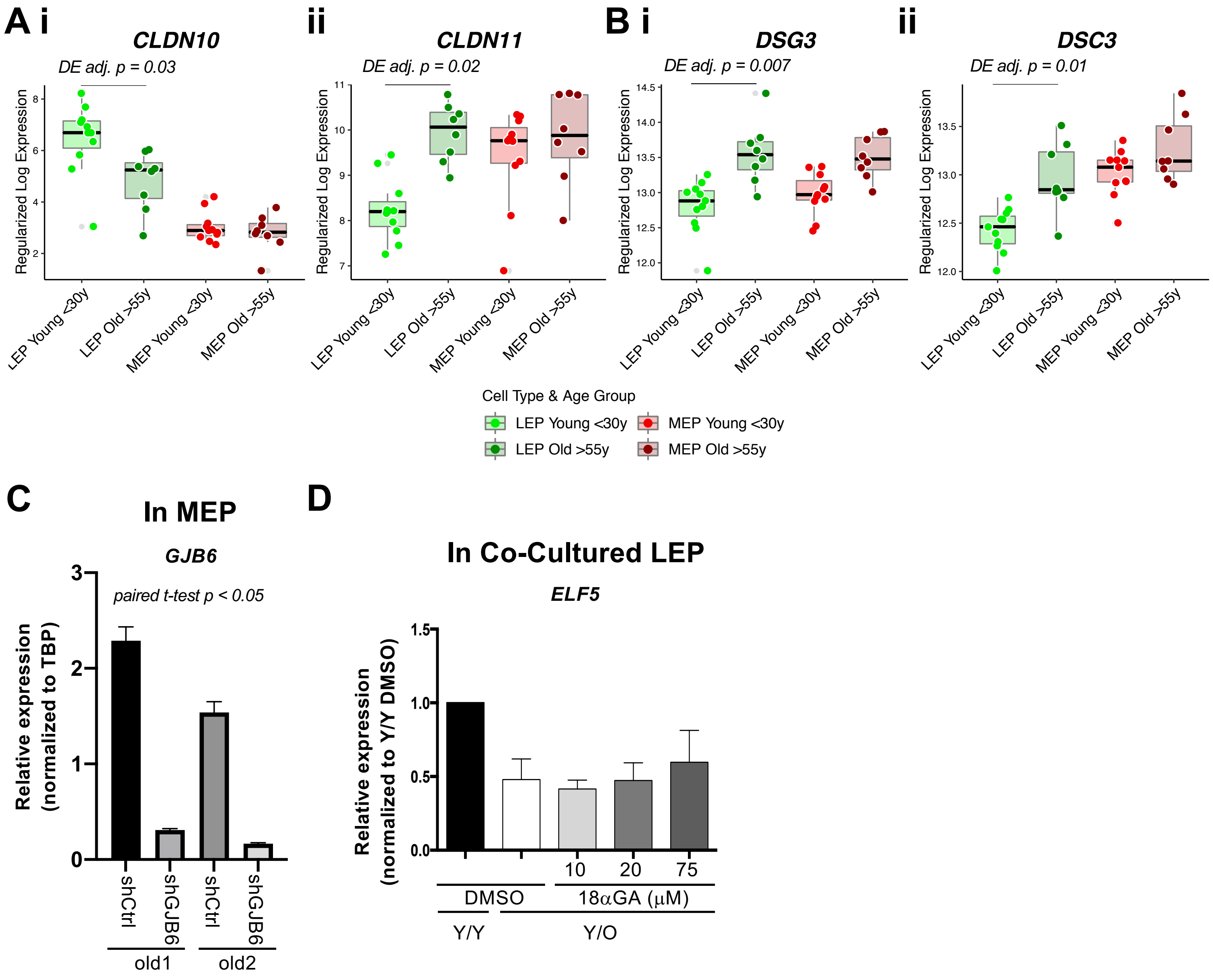
A i
CLDN10
DE adj. p = 0.03
ii
CLDN11
DE adj. p = 0.02
B i
DSG3
DE adj. p = 0.007
ii
DSC3
DE adj. p = 0.01
Regularized Log Expression
LEP Young <30y
LEP Old >55y
MEP Young <30y
MEP Old >55y
Cell Type & Age Group
C
In MEP
GJB6
paired t-test p < 0.05
Relative expression (normalized to TBP)
shCtrl
shGJB6
old1
old2
D
In Co-Cultured LEP
ELF5
Relative expression (normalized to Y/Y DMSO)
10
20
75
DMSO
18αGA (μM)
Y/Y
Y/O

**Figure S5. Hallmark pathways associated with cancer are dysregulated with age in luminal and myoepithelial lineages. Related to Figure 5.** (**A**) Boxplots of gene expression rlog values of (**i**) CLND10 and (**ii**) CLDN11 in LEPs and MEPs from younger and older women. Age-dependent DE adj. *p*-values (*limma*) are indicated. (**B**) Boxplots of gene expression rlog values of (**i**) DSG3 and (**ii**) DSC3 in LEPs and MEPs from younger and older women. Age-dependent DE adj. *p*-values (*limma*) are indicated. (**C**) Relative expression of *GJB6* in either shControl or shGJB6 older MEPs in two individuals. One-tailed paired t-test p-value indicated. (**D**) Relative expression of *ELF5* in younger LEPs co-cultured with older MEPs treated with increasing concentrations of 18aGA compared to Y/Y and Y/O treated with DMSO.

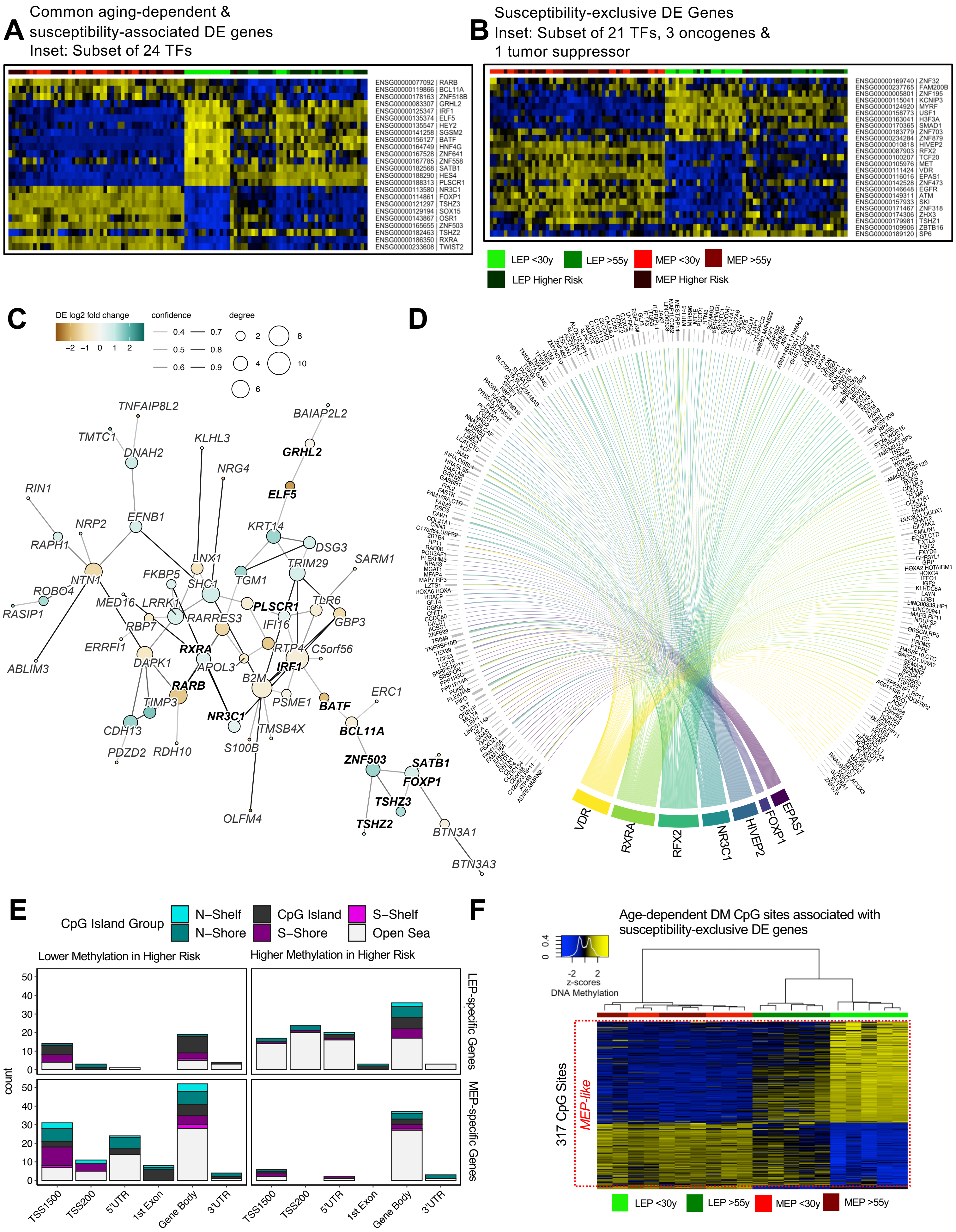
A
Common aging-dependent &
susceptibility-associated DE genes
Inset: Subset of 24 TFs
ENSG00000077092 | RARB
ENSG00000119866 | BCL11A
ENSG00000178163 | ZNF518B
ENSG00000083307 | GRHL2
ENSG00000125347 | IRF1
ENSG00000135374 | ELF5
ENSG00000135547 | HEY2
ENSG00000141258 | SGSM2
ENSG00000156127 | BATF
ENSG00000164749 | HNF4G
ENSG00000167528 | ZNF641
ENSG00000167785 | ZNF558
ENSG00000182568 | SATB1
ENSG00000188290 | HES4
ENSG00000188313 | PLSCR1
ENSG00000114861 | FOXP1
ENSG00000113580 | NR3C1
ENSG00000121297 | TSHZ3
ENSG00000129194 | SOX15
ENSG00000143867 | OSR1
ENSG00000165655 | ZNF503
ENSG00000182463 | TSHZ2
ENSG00000186350 | RXRA
ENSG00000233608 | TWIST2
B
Susceptibility-exclusive DE Genes
Inset: Subset of 21 TFs, 3 oncogenes &
1 tumor suppressor
ENSG00000169740 | ZNF32
ENSG00000237765 | FAM200B
ENSG00000005801 | ZNF195
ENSG00000115041 | KCNIP3
ENSG00000124920 | MYRF
ENSG00000163041 | H3F3A
ENSG00000170365 | SMAD1
ENSG00000183779 | ZNF703
ENSG00000158773 | USF1
ENSG00000234284 | ZNF879
ENSG00000010818 | HIVEP2
ENSG00000087903 | RFX2
ENSG00000100207 | TCF20
ENSG00000111424 | VDR
ENSG00000116016 | EPAS1
ENSG00000142528 | ZNF473
ENSG00000105976 | MET
ENSG00000146648 | EGFR
ENSG00000149311 | ATM
ENSG00000157933 | SKI
ENSG00000171467 | ZNF318
ENSG00000174306 | ZHX3
ENSG00000179981 | TSHZ1
ENSG00000109906 | ZBTB16
ENSG00000189120 | SP6
LEP <30y
LEP >55y
MEP <30y
MEP >55y
LEP Higher Risk
MEP Higher Risk
C
DE log2 fold change
−2 −1 0 1 2
confidence
0.4 0.5 0.6 0.7 0.8 0.9
degree
2 4 6 8 10
TNFAIP8L2
BAIAP2L2
TMTC1
DNAH2
KLHL3
GRHL2
NRG4
ELF5
RIN1
NRP2
EFNB1
KRT14
DSG3
RAPH1
LNX1
SARM1
TRIM29
FKBP5
TGM1
NTN1
SHC1
ROBO4
MED16
LRRK1
PLSCR1
TLR6
RASIP1
RARRES3
GBP3
RBP7
IFI16
ERRFI1
RXRA
RTP4
C5orf56
ABLIM3
DAPK1
APOL3
IRF1
B2M
RARB
ERC1
TIMP3
PSME1
BATF
NR3C1
CDH13
TMSB4X
BCL11A
PDZD2
RDH10
S100B
ZNF503
SATB1
FOXP1
TSHZ3
OLFM4
TSHZ2
BTN3A1
BTN3A3
D
VDR
RXRA
RFX2
NR3C1
HIVEP2
FOXP1
EPAS1
E
CpG Island Group
N−Shelf
N−Shore
CpG Island
S−Shore
S−Shelf
Open Sea
Lower Methylation in Higher Risk
Higher Methylation in Higher Risk
LEP-specific Genes
MEP-specific Genes
count
0 10 20 30 40 50
TSS1500
TSS200
5'UTR
1st Exon
Gene Body
3'UTR
F
Age-dependent DM CpG sites associated with
susceptibility-exclusive DE genes
0 0.4
-2 2
z-scores
DNA Methylation
317 CpG Sites
MEP-like
LEP <30y
LEP >55y
MEP <30y
MEP >55y

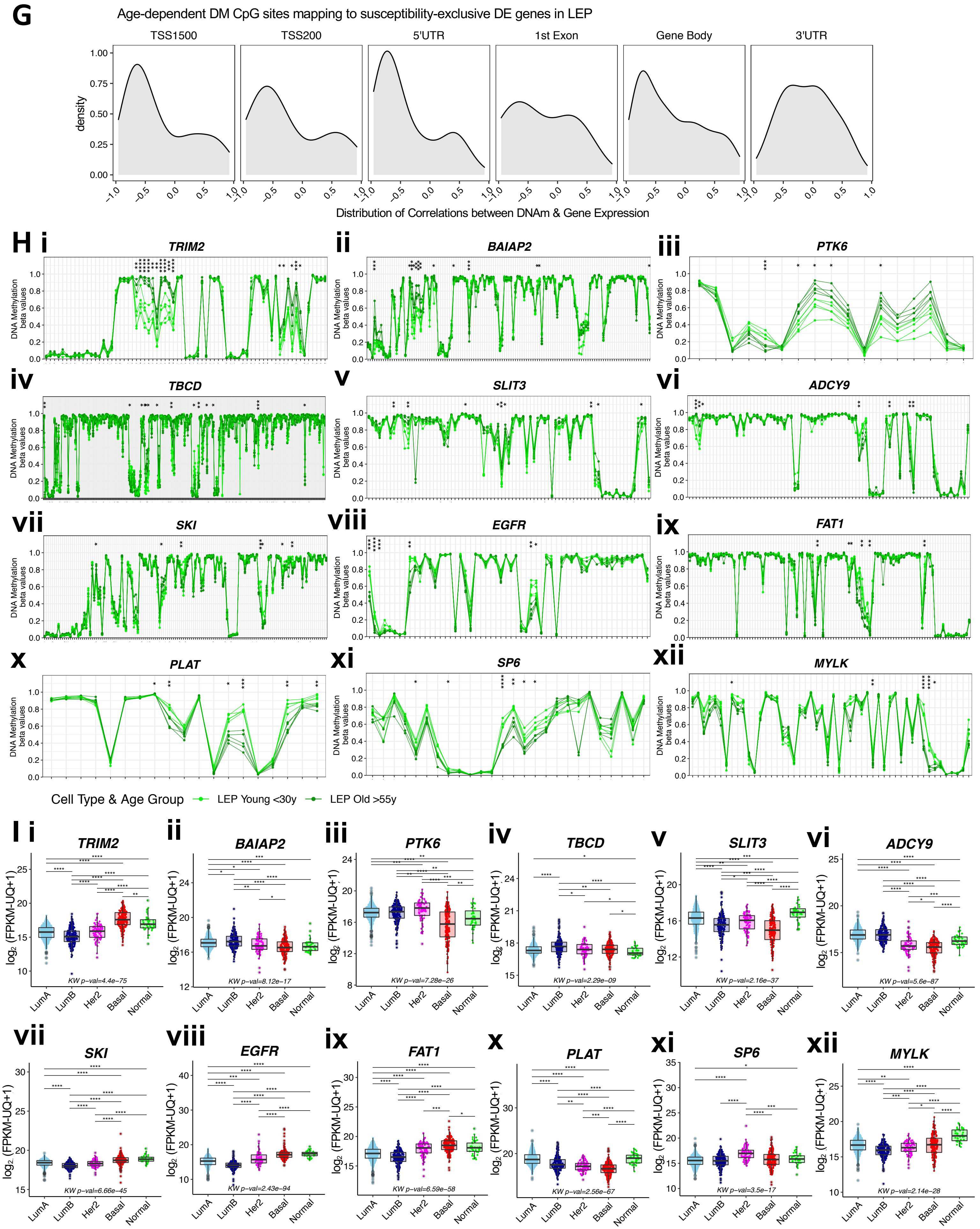
G
Age-dependent DM CpG sites mapping to susceptibility-exclusive DE genes in LEP
TSS1500
TSS200
5'UTR
1st Exon
Gene Body
3'UTR
density
Distribution of Correlations between DNAm & Gene Expression
H i
TRIM2
ii
BAIAP2
iii
PTK6
iv
TBCD
v
SLIT3
vi
ADCY9
vii
SKI
viii
EGFR
ix
FAT1
x
PLAT
xi
SP6
xii
MYLK
DNA Methylation beta values
Cell Type & Age Group
LEP Young <30y
LEP Old >55y
I i
TRIM2
KW p-val=4.4e-75
ii
BAIAP2
KW p-val=8.12e-17
iii
PTK6
KW p-val=7.28e-26
iv
TBCD
KW p-val=2.29e-09
v
SLIT3
KW p-val=2.16e-37
vi
ADCY9
KW p-val=5.6e-87
vii
SKI
KW p-val=6.66e-45
viii
EGFR
KW p-val=2.43e-94
ix
FAT1
KW p-val=6.59e-58
x
PLAT
KW p-val=2.56e-67
xi
SP6
KW p-val=3.5e-17
xii
MYLK
KW p-val=2.14e-28
log2 (FPKM-UQ+1)
LumA
LumB
Her2
Basal
Normal

**Figure S6. Age-dependent changes in methylation are priming events for increased susceptibility to breast cancer. Related to Figure 6.** (**A-B**) Heatmap of scaled rlog values for (**A**) common age-dependent and susceptibility-associated DE TFs (*limma adj. p* < 0.05) and (**B**) susceptibility-exclusive DE TFs, oncogenes and tumor suppressor genes (*limma adj. p* < 0.05) across all samples from younger AR, older AR and HR women. Samples in (**A**) and (**B**) are clustered based on **Figure 6B** and **6C** respectively. Scaled rlog values of DE genes are shown as a heatmap. (**C**) Protein-protein interaction (PPI) network (*stringdb* interaction score ≥ 0.4, medium confidence) of common age-dependent and susceptibility-associated DE genes (adj. *p* < 0.05), with TFs annotated in bold. Node sizes reflect gene connectivity, and node colors indicate the magnitude and direction of susceptibility-associated DE lfc (*limma*). (**D**) Differentially upregulated TFs (*adj. p* < 0.05) in LEPs of HR women and their corresponding enrichment (*ELMER* FDR *p* < 0.05) in gene promoter regions overlapping DMRs hypomethylated (*DMRcate* FDR *p* < 0.05) in LEPs of older women. (**E**) Number of age-dependent DM CpG sites mapping to susceptibility-exclusive DE genes and the proportion of annotated CpG island groups across different gene regions. (**F**) Hierarchical clustering of all samples from younger and older AR women based on age-dependent DM CpG sites mapping to susceptibility-exclusive DE genes (DE and DM *adj. p* < 0.05). Scaled beta values of DM CpG sites are shown as a heatmap. Clustering performed using Euclidean distances and Ward agglomerative method. (**G**) Density distribution of Spearman correlation coefficients for gene-CpG pairs associated with age-dependent DM CpG sites mapping to susceptibility-exclusive DE genes (DE and DM *adj. p* < 0.05) across gene regions. (**H**) DNA-m beta values of younger and older LEPs at CpG sites mapping to (**i**) *TRIM2*, (**ii**) *BAIAP2*, (**iii**) *PTK6*, (**iv**) *TBCD*, (**v**) *SLIT3*, (**vi**) *ADCY9*, (**vii**) *SKI*, (**viii**) *EGFR*, (**ix**) *FAT1*, (**x**) *PLAT*, (**xi**) *SP6*, and (**xii**) *MYLK*. CpG sites ordered by genomic positions. Age-dependent DM adj. *p*-value significance indicated (* < 0.05, ** < 0.01, *** < 0.001, **** < 0.0001) (*limma*). (**I**) Boxplots of gene expression $\log_2$ FPKM values of (**i**) *TRIM2*, (**ii**) *BAIAP2*, (**iii**) *PTK6*, (**iv**) *TBCD*, (**v**) *SLIT3*, (**vi**) *ADCY9*, (**vii**) *SKI*, (**viii**) *EGFR*, (**ix**) *FAT1*, (**x**) *PLAT*, (**xi**) *SP6*, and (**xii**) *MYLK* in the TCGA breast cancer cohort across cancer subtypes. KW test *p*-value between cancer subtypes in TCGA and post-hoc pair-wise Wilcoxon test adj. *p*-value significance levels are annotated (* < 0.05, ** < 0.01, *** < 0.001, **** < 0.0001).

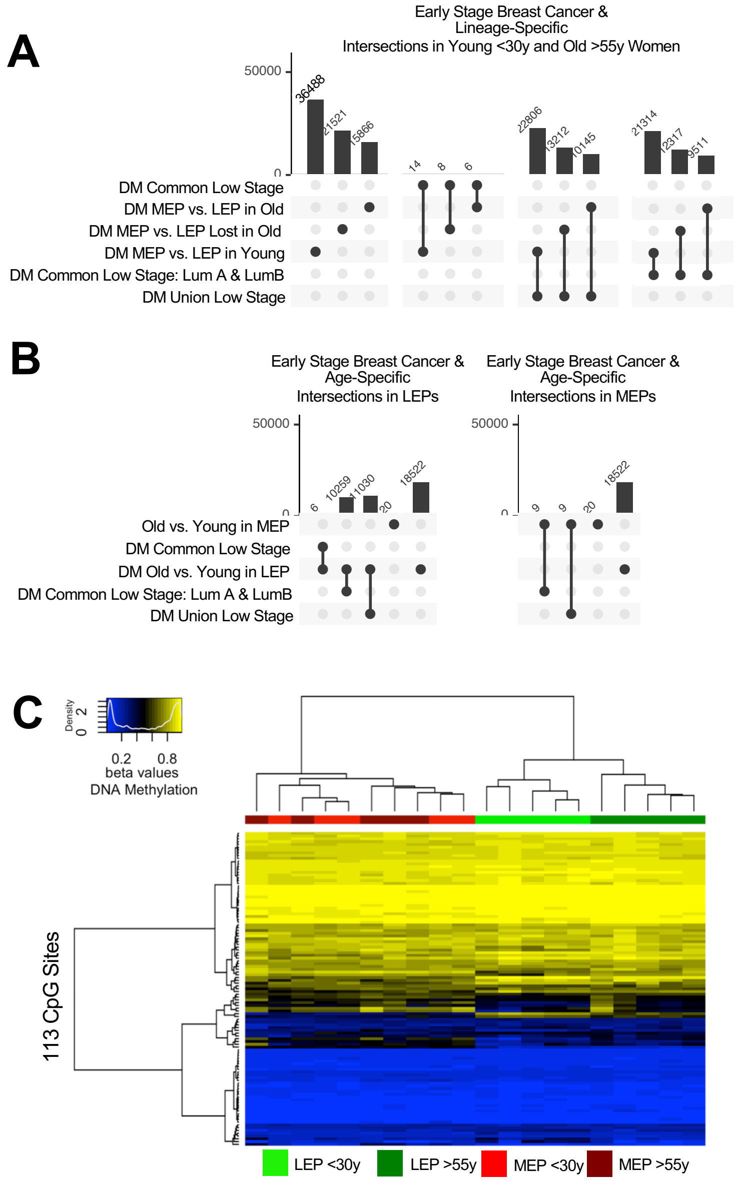


D

| Gene SYM | # Probe | # DM Probes | % DM Probes |
|---|---|---|---|
| DNMT3L | 12 | 3 | 25.0 |
| CXXC5 | 50 | 12 | 24.0 |
| TET1 | 30 | 4 | 13.3 |
| KDM2B | 84 | 9 | 10.7 |
| FBXL19 | 32 | 3 | 9.4 |
| DNMT3A | 79 | 7 | 8.9 |
| ZBTB4 | 38 | 3 | 7.9 |
| CXXC1 | 14 | 1 | 7.1 |
| KDM2A | 35 | 2 | 5.7 |
| ZBTB38 | 43 | 1 | 2.3 |

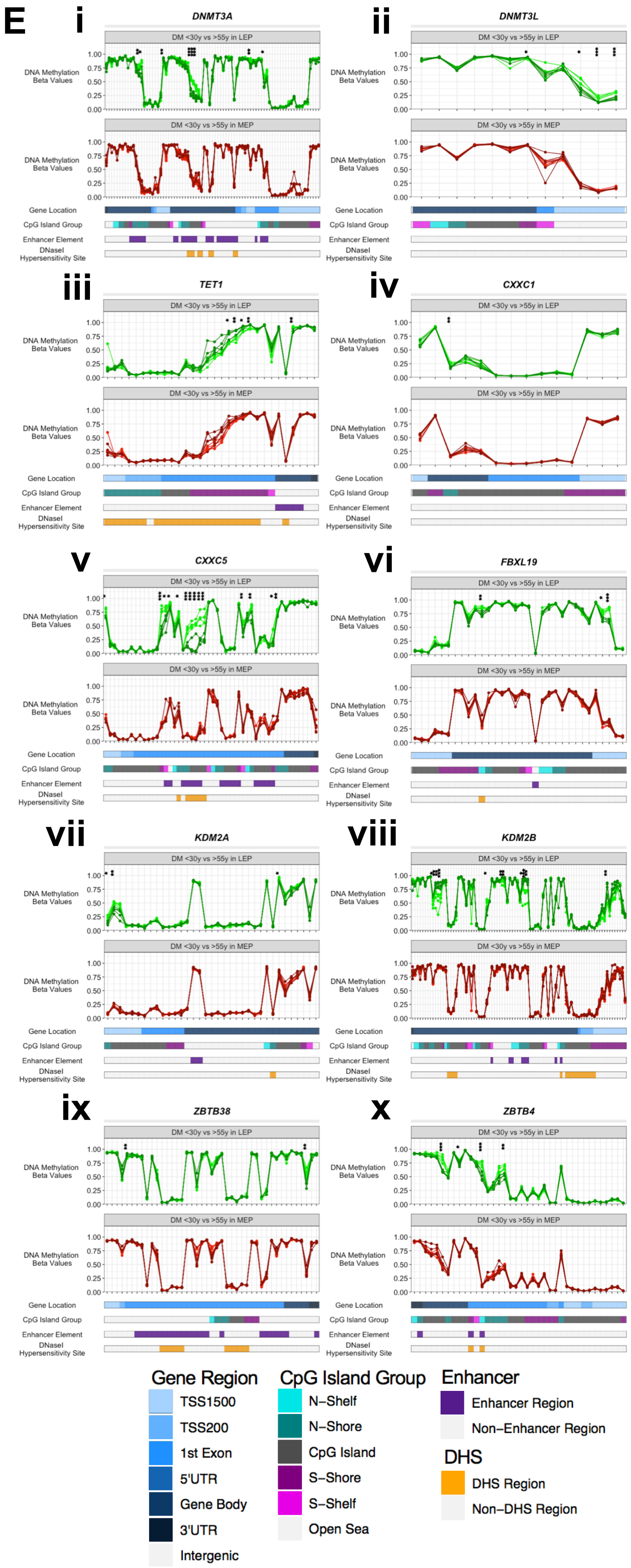

F

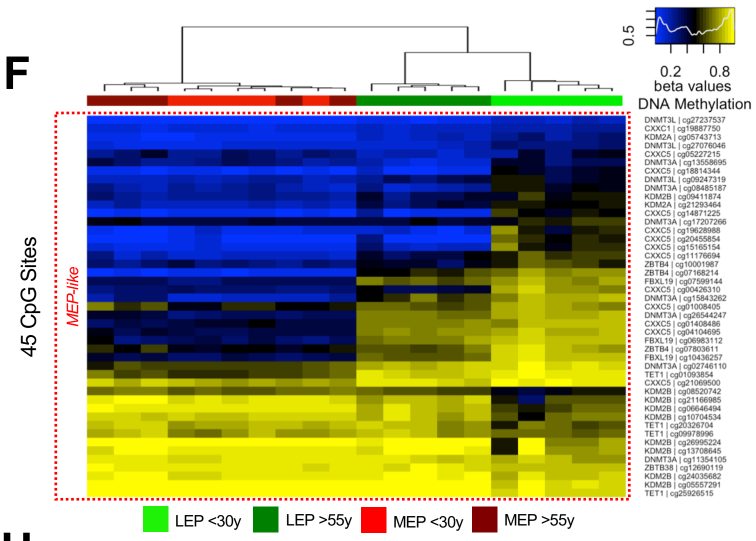


G

| Gene SYM | Gene Region | CpG Island Group | # DM Probes |
|---|---|---|---|
| CXXC5 | PPR | CpG Shelf & Shore | 8 |
| CXXC5 | PPR | CpG Island | 4 |
| KDM2B | Gene Body | CpG Island | 4 |
| DNMT3L | PPR | Open Sea | 3 |
| KDM2B | Gene Body | CpG Shelf & Shore | 3 |
| TET1 | PPR | CpG Shelf & Shore | 3 |
| DNMT3A | Gene Body | CpG Island | 2 |
| DNMT3A | Gene Body | CpG Shelf & Shore | 2 |
| FBXL19 | PPR | CpG Island | 2 |
| KDM2B | Gene Body | Open Sea | 2 |
| ZBTB4 | PPR | CpG Shelf & Shore | 2 |
| CXXC1 | Gene Body | CpG Shelf & Shore | 1 |
| DNMT3A | PPR | CpG Island | 1 |
| DNMT3A | PPR | CpG Shelf & Shore | 1 |
| DNMT3A | PPR | Open Sea | 1 |
| FBXL19 | Gene Body | CpG Shelf & Shore | 1 |
| KDM2A | PPR | CpG Island | 1 |
| KDM2A | PPR | CpG Shelf & Shore | 1 |
| TET1 | Gene Body | Open Sea | 1 |
| ZBTB38 | PPR | Open Sea | 1 |
| ZBTB4 | Gene Body | CpG Island | 1 |

H

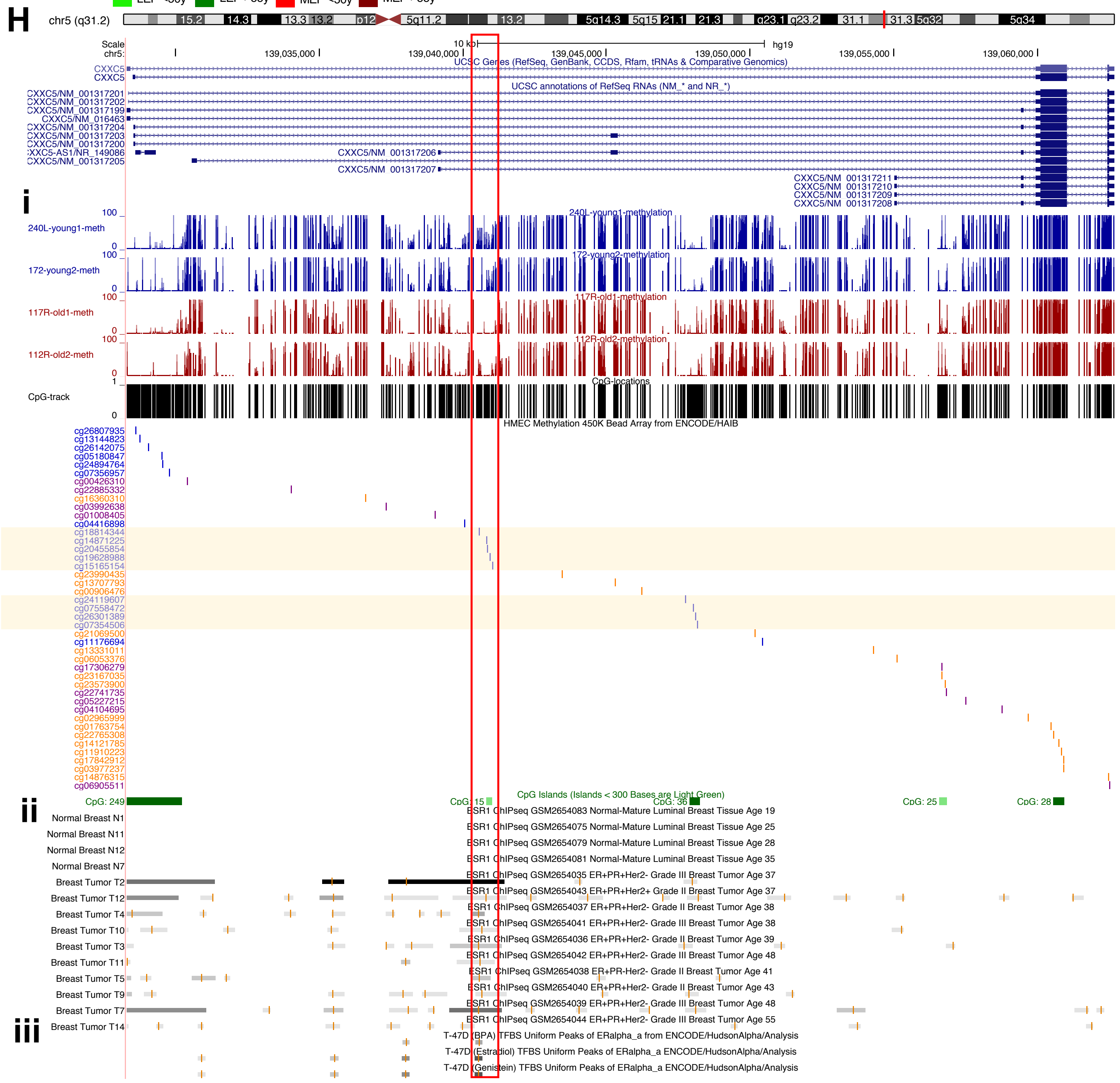

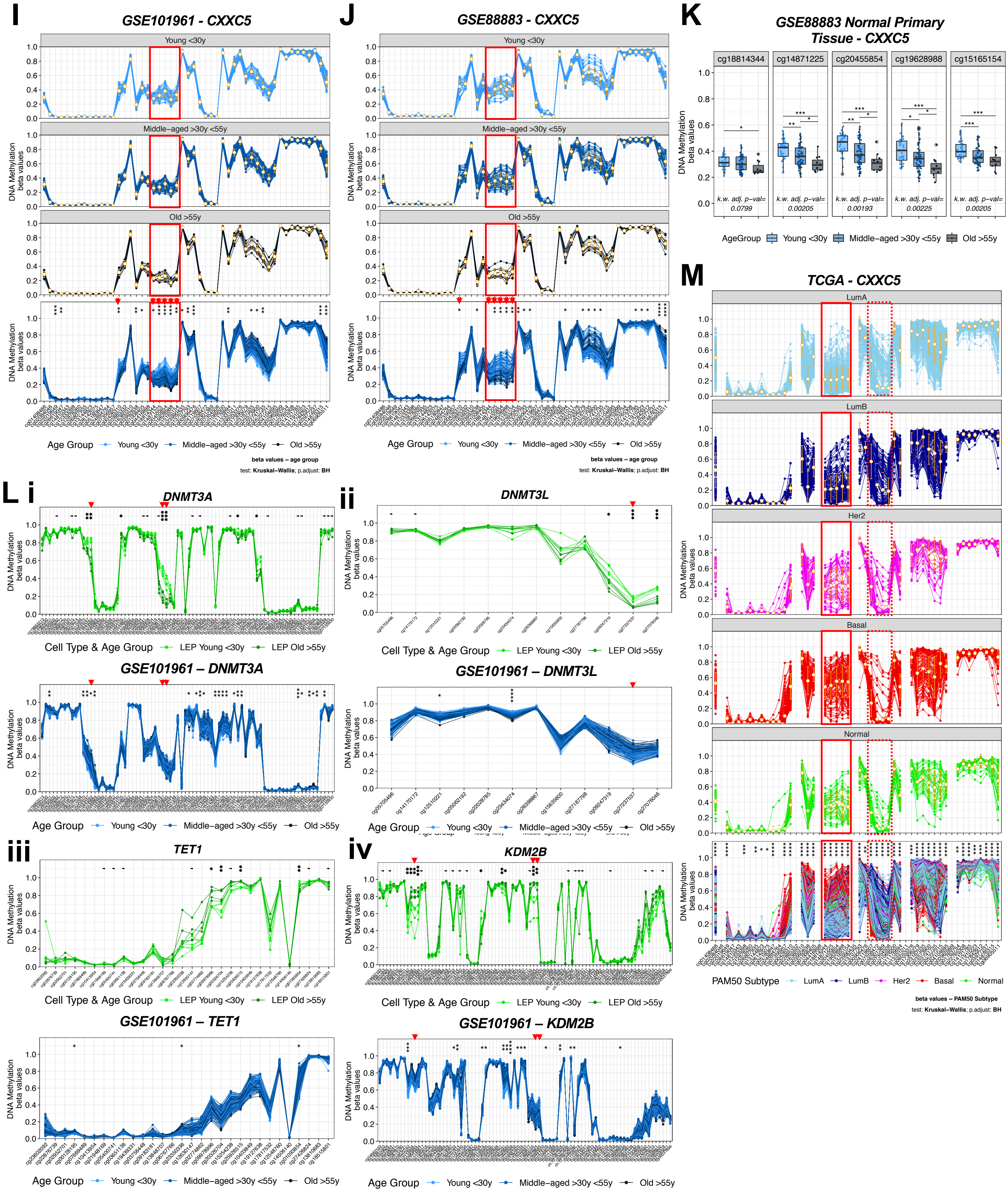
I
GSE101961 - CXXC5
Young <30y
Middle–aged >30y <55y
Old >55y
DNA Methylation beta values
Age Group Young <30y Middle–aged >30y <55y Old >55y
beta values ~ age group
test: Kruskal–Wallis; p.adjust: BH
J
GSE88883 - CXXC5
Young <30y
Middle–aged >30y <55y
Old >55y
DNA Methylation beta values
Age Group Young <30y Middle–aged >30y <55y Old >55y
beta values ~ age group
test: Kruskal–Wallis; p.adjust: BH
K
GSE88883 Normal Primary Tissue - CXXC5
cg18814344
cg14871225
cg20455854
cg19628988
cg15165154
DNA Methylation beta values
k.w. adj. p-val= 0.0799
k.w. adj. p-val= 0.00205
k.w. adj. p-val= 0.00193
k.w. adj. p-val= 0.00225
k.w. adj. p-val= 0.00205
AgeGroup Young <30y Middle–aged >30y <55y Old >55y
M
TCGA - CXXC5
LumA
LumB
Her2
Basal
Normal
DNA Methylation beta values
PAM50 Subtype LumA LumB Her2 Basal Normal
beta values ~ PAM50 Subtype
test: Kruskal–Wallis; p.adjust: BH
L i
DNMT3A
DNA Methylation beta values
Cell Type & Age Group LEP Young <30y LEP Old >55y
GSE101961 – DNMT3A
Age Group Young <30y Middle–aged >30y <55y Old >55y
ii
DNMT3L
Cell Type & Age Group LEP Young <30y LEP Old >55y
GSE101961 – DNMT3L
Age Group Young <30y Middle–aged >30y <55y Old >55y
iii
TET1
Cell Type & Age Group LEP Young <30y LEP Old >55y
GSE101961 – TET1
Age Group Young <30y Middle–aged >30y <55y Old >55y
iv
KDM2B
Cell Type & Age Group LEP Young <30y LEP Old >55y
GSE101961 – KDM2B
Age Group Young <30y Middle–aged >30y <55y Old >55y

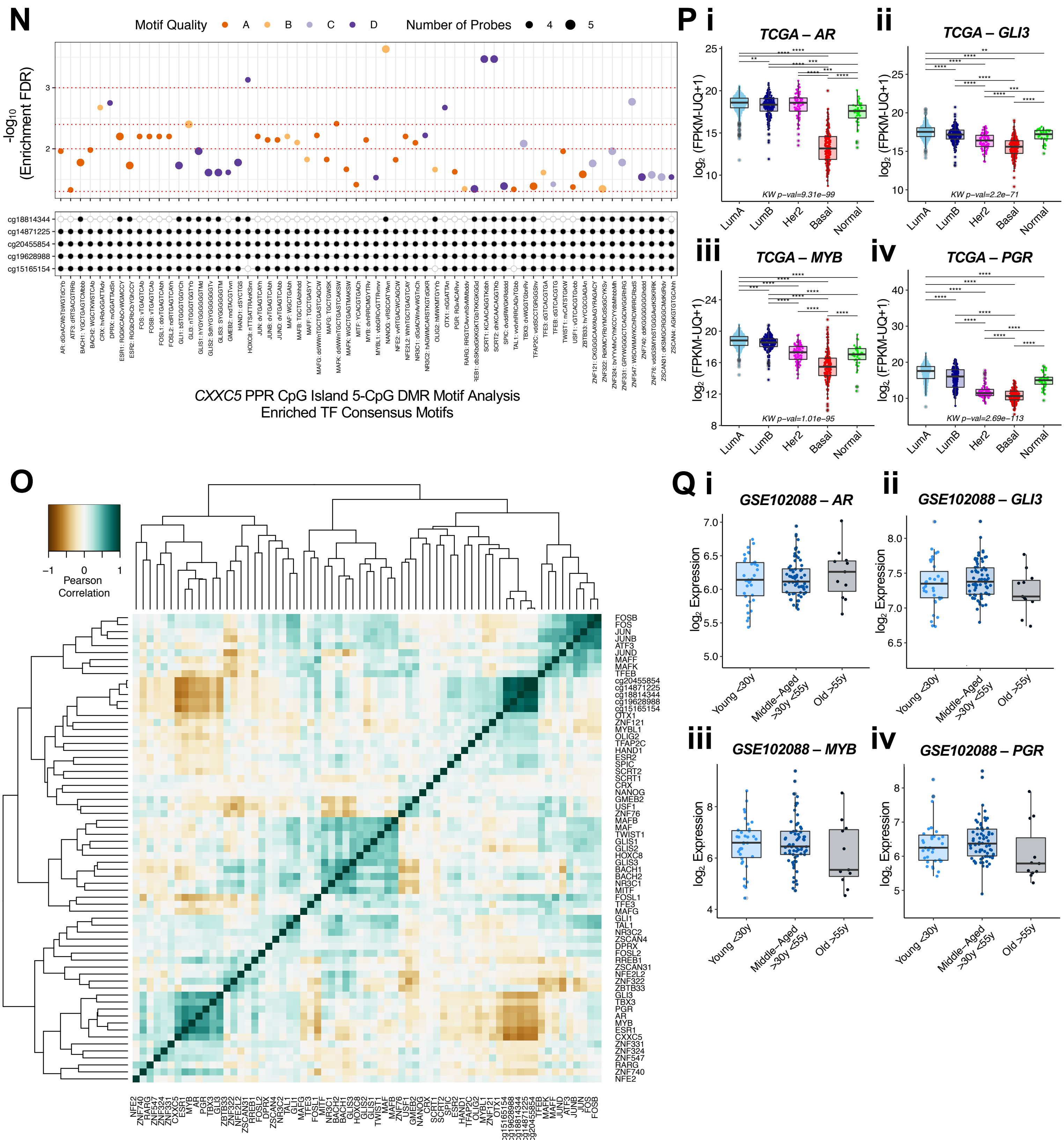
N
Motif Quality A B C D
Number of Probes 4 5
-log10 (Enrichment FDR)
cg18814344
cg14871225
cg20455854
cg19628988
cg15165154
CXXC5 PPR CpG Island 5-CpG DMR Motif Analysis
Enriched TF Consensus Motifs
P i
TCGA – AR
ii
TCGA – GLI3
iii
TCGA – MYB
iv
TCGA – PGR
log2 (FPKM-UQ+1)
KW p-val=9.31e-99
KW p-val=2.2e-71
KW p-val=1.01e-95
KW p-val=2.69e-113
LumA
LumB
Her2
Basal
Normal
O
Pearson Correlation
Q i
GSE102088 – AR
ii
GSE102088 – GLI3
iii
GSE102088 – MYB
iv
GSE102088 – PGR
log2 Expression
Young <30y
Middle-Aged >30y <55y
Old >55y

**Figure S7. Aging-associated hypomethylation of TF-binding sites in *CXXC5* primes epithelia for *CXXC5* dysregulation in luminal-subtype cancers**. **Related to Figure 7.** (**A-B**) Upset plot showing the number of intersecting genes between CpGs mapping to DMGRs in Titus et al., associated with all early stage breast cancers (union low), in common across early stage breast cancers (common low), or in common in early stage PAM50 LumA and LumB breast cancers [Titus et al., 2017a], and (**A**) lineage-specific DM CpG sites (*limma adj. p* < 0.001, fold change ≥ 2) in younger and older women or (**B**) age-dependent DM CpG sites (*limma adj. p* < 0.05) in LEPs and MEPs. (**C**) Hierarchical clustering of LEP and MEP samples based on CpG sites mapping to DMGRs identified by Titus et. al. to be in common across all breast cancer subtypes in early stage breast cancers in the TCGA cohort. DNAm beta values of DM CpG sites are shown as a heatmap. Clustering performed using Euclidean distances and Ward agglomerative method. (**D**) Table of genes encoding for known DNAm-regulatory proteins with the number of mapped CpG probes, and the number and percent that are DM with age in LEPs (*adj. p* < 0.05). (**E**) DNAm beta values of LEPs (top) and MEPs (bottom) from younger and older women at CpG sites mapping to (**i**) *DNMT3A* (**ii**) *DNMT3L* (**iii**) *TET1* (**iv**) *CXXC1* (**v**) *CXXC5* (**vi**) *FBXL19* (**vii**) *KDM2A* (**viii**) *KDM2B* (**ix**) *ZBTB38* and (**x**) *ZBTB4*. CpG sites ordered by genomic positions; gene regions, CpG island groups, enhancer element regions, and DHS indicated. Age-dependent DM adj. *p*-value significance levels (*limma*) indicated (* < 0.05, ** < 0.01, *** < 0.001, **** < 0.0001). (**F**) Hierarchical clustering of LEP and MEP samples based on DM CpG sites mapping to the ten DNAm-regulatory proteins identified to have age-dependent DM (*adj. p* < 0.05). DNAm beta values of DM CpG sites are shown as a heatmap. Clustering performed using Euclidean distances and Ward agglomerative method. (**G**) Table of the DM CpG sites mapping to the ten DNAm-regulatory proteins indicating the number of DM probes in each gene region and CpG island group. (**H**) *CXXC5* gene transcripts shown in UCSC Genome Browser with mapped Infinium 450K CpG probes, CpG island tracks annotated along with (**i**) WGBS data [Senapati et al., 2020] for *CXXC5* in LEPs from younger (blue) and older (red) women; and (**ii-iii**) publicly available *ESR1* ChIP-seq peaks from (**ii**) normal breast tissues and ER+ breast tumors, GSE99680 [Chi et al., 2019]; and (**iii**) T-47D ER+ breast cancer cell line [ENCODE Project Consortium, 2011, 2012] overlapping the *CXXC5* 5-CpG DMR (*DMRcate* FDR *p* < 0.05). (**I-J**) *CXXC5* DNAm beta values in two independent normal primary breast tissue datasets, (**I**) GSE101961 and (**J**) GSE88883 [Song et al., 2017, Johnson et al., 2017] across three age groups: younger <30y, middle-aged >30y <55y, and older >55y, with mean and sd shown (orange). Bottom panel shows DNAm values overlapped across all samples with KW test BH-adj. *p*-values across CpG sites between age groups in normal primary tissue and post-hoc pair-wise Wilcoxon test adj. p-value significance levels are annotated (* < 0.05, ** < 0.01, *** < 0.001, **** < 0.0001). Red arrows indicate corresponding CpGs in bulk tissue that are DM with age in LEPs with lfc > 1.5 (*adj. p* < 0.05) (**Figure 7B**). Red box highlights the *CXXC5* 5-CpG DMR (FDR *p* < 0.05) of interest. (**K**) Boxplots of DNAm beta values at the five contiguous *CXXC5* DM CpG sites in the age-dependent DMR (FDR *p* < 0.05) in a second independent normal primary breast tissue data set, GSE88883 [Johnson et al., 2017], across age groups. KW test BH-adj. p-values across CpG sites between age groups in normal primary tissue and post-hoc pair-wise Wilcoxon test adj. p-value significance levels are annotated (* < 0.05, ** < 0.01, *** < 0.001, **** < 0.0001). (**L**) DNAm beta values in LEPs (top) and in an independent normal primary breast tissue dataset, GSE101961 (bottom) [Song et al., 2017] across age groups for (**i**) *DNMT3A* (**ii**) *DNMT3L* (**iii**) *TET1* and (**iv**) *KDM2B*. Age-dependent DM adj. p-value and significance levels in LEPs (*limma*); and KW test BH-adj. p-values across CpG sites between age groups in normal primary tissue and between cancer subtypes in TCGA, and post-hoc pair-wise Wilcoxon test adj. p-value significance levels are annotated (* < 0.05, ** < 0.01, *** < 0.001, **** < 0.0001). Red arrows indicate CpGs that are DM with age in LEPs with lfc > 1.5 (*adj. p* < 0.05) and corresponding CpGs in bulk tissue. (**M**) *CXXC5* DNAm beta values in TCGA breast cancer dataset across PAM50 cancer subtypes: LumA, LumB, Her2, Basal and Normal, with mean and sd shown (orange). Bottom panel shows

DNAm values overlapped across all samples with KW test BH-adj. *p*-values across CpG sites between cancer subtypes in TCGA and post-hoc pair-wise Wilcoxon test adj. p-value significance levels are annotated (* < 0.05, ** < 0.01, *** < 0.001, **** < 0.0001). Red box highlights the *CXXC5* 5-CpG DMR (FDR $p < 0.05$) of interest. Red dotted box highlights example region consisting of CpG sites: *cg24119607, cg07558472, cg26301389, cg07354506* that is DM between PAM50 subtypes but is unmethylated and does not change with age in normal LEPs or normal tissue. (**N**) TF motif analysis (*ELMER* FDR $p < 0.05$) at the *CXXC5* 5-CpG DMR. Enrichment FDR *p*-values ($-\log_{10}$) for each motif is plotted, with number of probes reflected by point size, and motif quality in color. (**O**) Heatmap of Pearson correlations between *CXXC5* gene expression, *CXXC5* DNAm at the five contiguous DM CpG sites, and expression of TFs with enriched motifs (*ELMER* FDR $p < 0.05$) at the *CXXC5* 5-CpG DMR in the TCGA cohort. Clustering performed using complete agglomerative method with 1-correlation as distance metric. (**P-Q**) Boxplot of gene expression values of (**i**) *AR* (**ii**) *GLI3* (**iii**) *MYB* and (**iv**) *PGR* (**P**) in the TCGA cohort across cancer subtypes, or (**Q**) in normal primary breast tissue, GSE10208 [Song et al., 2017] across age groups. No significant age-dependent DE in normal primary tissue (*limma*) detected. KW test *p*-value between cancer subtypes in TCGA and post-hoc pair-wise Wilcoxon test adj. *p*-value significance levels are annotated (* < 0.05, ** < 0.01, *** < 0.001, **** < 0.0001).

| Subject ID | Culture Condition | Cell Type | Age | Age Group | Tissue Type | Mutation | RNASeq Project ID |
|---|---|---|---|---|---|---|---|
| 160 | p4 | LEP | 16 | Young | RM | None | 170112 |
| 160 | p4 | MEP | 16 | Young | RM | None | 170112 |
| 48R | p4 | LEP | 16 | Young | RM | None | 181212 |
| 48R | p4 | MEP | 16 | Young | RM | None | 181212 |
| 168R | p4 | LEP | 19 | Young | RM | None | 181212 |
| 168R | p4 | MEP | 19 | Young | RM | None | 181212 |
| 240L | p4 | LEP | 19 | Young | RM | None | 170112 |
| 240L | p4 | LEP | 19 | Young | RM | None | 181212 |
| 240L | p4 | LEP | 19 | Young | RM | None | 180928 |
| 240L | p4 | LEP | 19 | Young | RM | None | 200211 |
| 240L | p4 | MEP | 19 | Young | RM | None | 170112 |
| 240L | p4 | MEP | 19 | Young | RM | None | 181212 |
| 240L | p4 | MEP | 19 | Young | RM | None | 180928 |
| 240L | p4 | MEP | 19 | Young | RM | None | 200211 |
| 184D | p4 | LEP | 21 | Young | RM | None | 181212 |
| 184D | p4 | LEP | 21 | Young | RM | None | 180928 |
| 184D | p4 | MEP | 21 | Young | RM | None | 181212 |
| 184D | p4 | MEP | 21 | Young | RM | None | 180928 |
| 356E | p4 | LEP | 21 | Young | RM | None | 181212 |
| 356E | p4 | MEP | 21 | Young | RM | None | 181212 |
| 59L | p4 | LEP | 23 | Young | RM | None | 170112 |
| 59L | p4 | MEP | 23 | Young | RM | None | 170112 |
| 163 | p4 | LEP | 27 | Young | RM | None | 200211 |
| 163 | p4 | MEP | 27 | Young | RM | None | 200211 |
| 51L | p4 | LEP | 27 | Young | RM | None | 170112 |
| 51L | p4 | MEP | 27 | Young | RM | None | 170112 |
| 172L | p4 | LEP | 28 | Young | RM | None | 170112 |
| 172L | p4 | MEP | 28 | Young | RM | None | 170112 |
| 124 | p4 | LEP | 29 | Young | RM | None | 170112 |
| 124 | p4 | LEP | 29 | Young | RM | None | 180921 |
| 124 | p4 | MEP | 29 | Young | RM | None | 170112 |
| 124 | p4 | MEP | 29 | Young | RM | None | 180921 |
| 117R | p4 | LEP | 56 | Old | RM | None | 181212 |
| 117R | p4 | LEP | 56 | Old | RM | None | 180921 |
| 117R | p4 | MEP | 56 | Old | RM | None | 181212 |
| 117R | p4 | MEP | 56 | Old | RM | None | 180921 |
| 191L | p4 | LEP | 56 | Old | RM | None | 170112 |
| 191L | p4 | MEP | 56 | Old | RM | None | 170112 |
| 153L | p4 | LEP | 60 | Old | RM | None | 181212 |
| 153L | p4 | MEP | 60 | Old | RM | None | 181212 |
| 112R | p4 | LEP | 61 | Old | RM | None | 170112 |
| 112R | p4 | LEP | 61 | Old | RM | None | 181212 |
| 112R | p4 | LEP | 61 | Old | RM | None | 200211 |
| 112R | p4 | MEP | 61 | Old | RM | None | 170112 |
| 112R | p4 | MEP | 61 | Old | RM | None | 181212 |
| 112R | p4 | MEP | 61 | Old | RM | None | 200211 |
| 237 | p4 | LEP | 66 | Old | RM | None | 170112 |
| 237 | p4 | MEP | 66 | Old | RM | None | 170112 |
| 122L | p4 | LEP | 66 | Old | RM | None | 170112 |
| 122L | p4 | MEP | 66 | Old | RM | None | 170112 |
| 29 | p4 | LEP | 68 | Old | RM | None | 181212 |
| 29 | p4 | MEP | 68 | Old | RM | None | 181212 |
| 429ER | p4 | LEP | 72 | Old | RM | None | 181212 |
| 429ER | p4 | MEP | 72 | Old | RM | None | 181212 |
| 353P | p4 | LEP | 72 | Old | PTT | None | 170112 |
| 353P | p4 | MEP | 72 | Old | PTT | None | 170112 |
| 249P | p4 | LEP | 31 | Middleaged | PTT | BRCA1 | 170112 |
| 249P | p4 | MEP | 31 | Middleaged | PTT | BRCA1 | 170112 |
| 90P | p4 | LEP | 35 | Middleaged | PTT | BRCA1 | 170112 |
| 90P | p4 | MEP | 35 | Middleaged | PTT | BRCA1 | 170112 |
| 101P | p4 | LEP | 43 | Middleaged | PTT | None | 170112 |
| 101P | p4 | MEP | 43 | Middleaged | PTT | None | 170112 |
| C023R | p4 | LEP | 35 | Middleaged | PM | BRCA1 | 180921 |
| C023R | p4 | MEP | 35 | Middleaged | PM | BRCA1 | 180921 |
| C004R | p4 | LEP | 40 | Middleaged | PM | BRCA1 | 180921 |
| C004R | p4 | MEP | 40 | Middleaged | PM | BRCA1 | 180921 |
| C009R | p4 | LEP | 40 | Middleaged | PM | BRCA1 | 180921 |
| C009R | p4 | MEP | 40 | Middleaged | PM | BRCA1 | 180921 |
| C002R | p4 | LEP | 50 | Middleaged | PM | BRCA1 | 180921 |
| C002R | p4 | MEP | 50 | Middleaged | PM | BRCA1 | 180921 |
| C003R2 | p4 | LEP | 33 | Middleaged | PM | BRCA2 | 180921 |
| C003R2 | p4 | MEP | 33 | Middleaged | PM | BRCA2 | 180921 |
| C001R | p4 | LEP | 52 | Middleaged | PM | BRCA2 | 180921 |
| C001R | p4 | MEP | 52 | Middleaged | PM | BRCA2 | 180921 |
| C017L | p4 | LEP | 36 | Middleaged | PM | None | 180921 |
| C017L | p4 | MEP | 36 | Middleaged | PM | None | 180921 |
| C014R | p4 | LEP | 52 | Middleaged | PM | PALB2 | 180921 |
| C014R | p4 | MEP | 52 | Middleaged | PM | PALB2 | 180921 |
| 247C | p4 | LEP | 25 | Young | CLTT | None | 170112 |
| 247C | p4 | MEP | 25 | Young | CLTT | None | 170112 |
| 211C | p4 | LEP | 64 | Old | CLTT | None | 170112 |
| 211C | p4 | MEP | 64 | Old | CLTT | None | 170112 |
| 71C | p4 | LEP | 65 | Old | CLTT | None | 170112 |
| 71C | p4 | MEP | 65 | Old | CLTT | None | 170112 |
| 257C | p4 | LEP | 67 | Old | CLTT | None | 170112 |
| 257C | p4 | MEP | 67 | Old | CLTT | None | 170112 |
| C039C | p4 | LEP | 55 | Old | CLTT | PALB2 VUS, APC VUS | 180921 |
| C039C | p4 | MEP | 55 | Old | CLTT | PALB2 VUS, APC VUS | 180921 |
| C063C | p4 | LEP | 36 | Middleaged | CLTT | BRCA1 | 180921 |
| C063C | p4 | MEP | 36 | Middleaged | CLTT | BRCA1 | 180921 |
| C008C | p4 | LEP | 53 | Middleaged | CLTT | BRCA1 | 180921 |
| C008C | p4 | MEP | 53 | Middleaged | CLTT | BRCA1 | 180921 |
| C020C | p4 | LEP | 44 | Middleaged | CLTT | BRCA2 | 180921 |
| C020C | p4 | MEP | 44 | Middleaged | CLTT | BRCA2 | 180921 |
| C018C | p4 | LEP | 40 | Middleaged | CLTT | None | 180921 |
| C018C | p4 | MEP | 40 | Middleaged | CLTT | None | 180921 |
| 210C | p4 | LEP | 42 | Middleaged | CLTT | None | 170112 |
| 210C | p4 | MEP | 42 | Middleaged | CLTT | None | 170112 |
| 215C | p4 | LEP | 44 | Middleaged | CLTT | None | 170112 |
| 215C | p4 | MEP | 44 | Middleaged | CLTT | None | 170112 |
| C019C | p4 | LEP | 54 | Middleaged | CLTT | None | 180928 |
| C019C | p4 | MEP | 54 | Middleaged | CLTT | None | 180928 |
| 160 | organoid | LEP | 16 | Young | RM | None | 200211 |
| 160 | organoid | MEP | 16 | Young | RM | None | 200211 |
| 195L | organoid | LEP | 24 | Young | RM | None | 200211 |
| 195L | organoid | MEP | 24 | Young | RM | None | 200211 |
| 51L | organoid | LEP | 27 | Young | RM | None | 200211 |
| 51L | organoid | MEP | 27 | Young | RM | None | 200211 |
| 124 | organoid | LEP | 29 | Young | RM | None | 200211 |
| 124 | organoid | MEP | 29 | Young | RM | None | 200211 |
| 112R | organoid | LEP | 61 | Old | RM | None | 200211 |
| 112R | organoid | MEP | 61 | Old | RM | None | 200211 |
| 96L | organoid | LEP | 62 | Old | RM | None | 200211 |
| 237 | organoid | LEP | 66 | Old | RM | None | 200211 |

**Table S1. RNA-sequencing Sample List.**

| Subject ID | Culture Condition | Cell Type | Age | Age Group | Tissue Type | Mutation | Slide ID | Array ID |
|---|---|---|---|---|---|---|---|---|
| 240L | p4 | MEP | 19 | Young | RM | None | 9970497131 | R01C01 |
| 240L | p4 | LEP | 19 | Young | RM | None | 9970497131 | R02C01 |
| 124 | p4 | MEP | 29 | Young | RM | None | 9970497131 | R03C01 |
| 124 | p4 | LEP | 29 | Young | RM | None | 9970497131 | R04C01 |
| 59L | p4 | MEP | 23 | Young | RM | None | 9970497131 | R05C01 |
| 59L | p4 | LEP | 23 | Young | RM | None | 9970497131 | R06C01 |
| 29 | p4 | MEP | 68 | Old | RM | None | 9970497131 | R01C02 |
| 29 | p4 | LEP | 68 | Old | RM | None | 9970497131 | R02C02 |
| 112R | p4 | MEP | 61 | Old | RM | None | 9970497131 | R03C02 |
| 240L | p4 | MEP | 19 | Young | RM | None | 9970497080 | R01C01 |
| 240L | p4 | LEP | 19 | Young | RM | None | 9970497080 | R02C01 |
| 51L | p4 | MEP | 27 | Young | RM | None | 9970497080 | R03C01 |
| 51L | p4 | LEP | 27 | Young | RM | None | 9970497080 | R04C01 |
| 122L | p4 | MEP | 66 | Old | RM | None | 9970497080 | R05C01 |
| 122L | p4 | LEP | 66 | Old | RM | None | 9970497080 | R06C01 |
| 237 | p4 | MEP | 66 | Old | RM | None | 9970497080 | R01C02 |
| 237 | p4 | LEP | 66 | Old | RM | None | 9970497080 | R02C02 |
| 71C | p4 | MEP | 65 | Old | CLTT | None | 9970497080 | R03C02 |
| 71C | p4 | LEP | 65 | Old | CLTT | None | 9970497080 | R04C02 |
| 112R | p4 | LEP | 61 | Old | RM | None | 9970497080 | R06C02 |

**Table S2. Infinium 450K DNA methylation Array Sample List.**
